# Environmental DNA (eDNA) metabarcoding of pond water as a tool to survey conservation and management priority mammals

**DOI:** 10.1101/546218

**Authors:** Lynsey R. Harper, Lori Lawson Handley, Angus I. Carpenter, Muhammad Ghazali, Cristina Di Muri, Callum J. Macgregor, Thomas W. Logan, Alan Law, Thomas Breithaupt, Daniel S. Read, Allan D. McDevitt, Bernd Hänfling

**Affiliations:** Department of Biological and Marine Sciences, University of Hull, Hull, HU6 7RX, UK; Illinois Natural History Survey, Prairie Research Institute, University of Illinois at Urbana-Champaign, Champaign, Illinois, USA; Wildwood Trust, Canterbury Rd, Herne Common, Herne Bay, CT6 7LQ, UK; The Royal Zoological Society of Scotland, Edinburgh Zoo, 134 Corstorphine Road, Edinburgh, EH12 6TS, UK; Department of Biology, University of York, Wentworth Way, York, YO10 5DD, UK; Biological and Environmental Sciences, University of Stirling, Stirling, FK9 4LA, UK; Centre for Ecology & Hydrology (CEH), Benson Lane, Crowmarsh Gifford, Wallingford, Oxfordshire, OX10 8BB, UK; Ecosystems and Environment Research Centre, School of Science, Engineering and Environment, University of Salford, Salford, M5 4WT, UK

**Keywords:** Key-words, camera traps, field signs, lentic, monitoring, semi-aquatic mammals, terrestrial mammals

## Abstract

Environmental DNA (eDNA) metabarcoding can identify terrestrial taxa utilising aquatic habitats alongside aquatic communities, but terrestrial species’ eDNA dynamics are understudied. We evaluated eDNA metabarcoding for monitoring semi-aquatic and terrestrial mammals, specifically nine species of conservation or management concern, and examined spatiotemporal variation in mammal eDNA signals. We hypothesised eDNA signals would be stronger for semi-aquatic than terrestrial mammals, and at sites where individuals exhibited behaviours. In captivity, we sampled waterbodies at points where behaviours were observed (‘directed’ sampling) and at equidistant intervals along the shoreline (‘stratified’ sampling). We surveyed natural ponds (*N* = 6) where focal species were present using stratified water sampling, camera traps, and field signs. eDNA samples were metabarcoded using vertebrate-specific primers. All focal species were detected in captivity. eDNA signal strength did not differ between directed and stratified samples across or within species, between semi-aquatic or terrestrial species, or according to behaviours. eDNA was evenly distributed in artificial waterbodies, but unevenly distributed in natural ponds. Survey methods deployed at natural ponds shared three species detections. Metabarcoding missed badger and red fox recorded by cameras and field signs, but detected small mammals these tools overlooked, e.g. water vole. Terrestrial mammal eDNA signals were weaker and detected less frequently than semi-aquatic mammal eDNA signals. eDNA metabarcoding could enhance mammal monitoring through large-scale, multi-species distribution assessment for priority and difficult to survey species, and provide early indication of range expansions or contractions. However, eDNA surveys need high spatiotemporal resolution and metabarcoding biases require further investigation before routine implementation.

## 1. Introduction

Mammals are a highly threatened taxon, with 25% of species at risk of extinction globally due to harvesting, habitat degradation/loss, non-native species or perception as pests (Visconti et al., 2011). Most species lack long-term, systematic monitoring, with survey efforts biased towards rare species (Massimino, Harris, & Gillings, 2018). Data deficiency prevents robust estimation of mammalian range expansions/declines and population trends (Bland, Collen, Orme, & Bielby, 2015). Therefore, effective and evidence-based strategies for mammal conservation and management are urgently needed (Mathews et al., 2018).

Many mammals are nocturnal and elusive thus monitoring requires non-invasive, observational methods such as camera traps and field signs, e.g. footprints, scat (Caravaggi et al., 2018; Harris & Yalden, 2004; Kinoshita et al., 2019; Sadlier, Webbon, Baker, & Harris, 2004). Camera trapping is cost-efficient, standardised, reproducible, and produces data suited to site occupancy modelling, but only surveys a fraction of large, heterogeneous landscapes. Trap placement can substantially influence species detection probabilities, and traps often miss small species (Burton et al., 2015; Caravaggi et al., 2018; Ishige et al., 2017; Leempoel, Hebert, & Hadly, 2019). Field sign surveys are inexpensive, but resource-intensive for broad geographic coverage (Kinoshita et al., 2019; Sadlier et al., 2004). Species can have similar footprints and scat, increasing the potential for misidentification (Franklin et al., 2019; Harris & Yalden, 2004). Mammal survey methods can be species-specific, thus multiple methods are necessary for large-scale, multi-species monitoring schemes (Massimino et al., 2018; Sales et al., 2019).

Environmental DNA (eDNA) analysis is a recognised tool for rapid, non-invasive, cost-efficient biodiversity assessment across aquatic and terrestrial ecosystems (Deiner et al., 2017). Organisms transfer genetic material to their environment via secretions, excretions, gametes, blood, or decomposition, which can be isolated from environmental samples (Thomsen & Willerslev, 2015). Studies using eDNA analysis to target specific semi-aquatic and terrestrial mammals have employed PCR or quantitative PCR (qPCR) (e.g. Franklin et al., 2019; Lugg, Griffiths, van Rooyen, Weeks, & Tingley, 2017; Rodgers & Mock, 2015; Thomsen et al., 2012; Williams, Huyvaert, Vercauteren, Davis, & Piaggio, 2018). eDNA metabarcoding can screen entire communities using PCR combined with high-throughput sequencing (Deiner et al., 2017; Thomsen & Willerslev, 2015), but mammalian assessments are uncommon (Klymus, Richter, Thompson, & Hinck, 2017; Kinoshita et al., 2019; Leempoel et al., 2019; Sales et al., 2019; Ushio et al., 2017). Tropical mammal assemblages have been obtained by metabarcoding invertebrate blood meals (e.g. Tessler et al., 2018) and salt licks (Ishige et al., 2017), but samples from the physical environment have tremendous potential to reveal mammal biodiversity over broad spatiotemporal scales (Sales et al., 2019; Ushio et al., 2017).

In aquatic ecosystems, eDNA metabarcoding has predominantly been applied to characterise fish (e.g. Evans et al., 2017; Hänfling et al., 2016; Lawson Handley et al., 2018; Valentini et al., 2016) and amphibian (e.g. Bálint et al., 2018; Valentini et al., 2016) communities. However, mammals also leave eDNA signatures in water that metabarcoding can detect (Harper et al., 2019; Klymus et al., 2017; Sales et al., 2019; Ushio et al., 2017). Ponds in particular provide drinking, foraging, dispersive, and reproductive opportunities for semi-aquatic and terrestrial mammals (Klymus et al., 2017). Samples from these waterbodies could uncover biodiversity present in the wider environment (Deiner et al., 2017; Harper et al., 2019). Drinking is a major source of eDNA deposition due to the release of saliva, but mammals may also swim, wallow, urinate or defecate in water (Rodgers & Mock, 2015; Ushio et al., 2017; Williams et al., 2018). Furthermore, arboreal mammals may use ponds less than semi-aquatic and ground-dwelling species, non-territorial mammals may visit ponds less than territorial species, and group-living species may deposit more eDNA than solitary species (Williams et al., 2018). Despite evidence for eDNA deposition by semi-aquatic and terrestrial mammals in freshwater ecosystems, little is known about the influence of mammal behaviour on the distribution and strength of the eDNA signal left behind (defined here as proportional read counts).

In this study, we conducted two experiments under artificial and natural conditions to evaluate eDNA metabarcoding of pond water as a tool for monitoring semi-aquatic, ground-dwelling, and arboreal mammals of conservation or management concern. The first experiment, carried out on nine focal species housed at two wildlife parks, examined the role of sampling strategy, mammal lifestyle, and mammal behaviour on eDNA detection and signal strength under artificial conditions. Mammal eDNA detection is expected from enclosure water that is frequently used by individuals for drinking, swimming and bathing. We hypothesised that: (1) eDNA would be unevenly distributed, thus directed sampling would yield stronger eDNA signals (i.e. higher proportional read counts) for mammals than stratified sampling; (2) semi-aquatic mammals would have stronger eDNA signals than ground-dwelling or arboreal mammals; and (3) mammal behaviours involving water contact would generate stronger eDNA signals. The second experiment validated eDNA metabarcoding against camera trapping and field sign searches for mammal identification at natural ponds, and investigated spatiotemporal variation in mammal eDNA signals. Mammal eDNA detection is unpredictable at natural waterbodies that can be extensive, subject to environmental fluctuations, and used rarely or not at all by individuals. We hypothesised that: (1) eDNA metabarcoding would detect more mammals than camera trapping or field signs; (2) semi-aquatic mammals would be readily detected and their eDNA evenly distributed in ponds in comparison to terrestrial mammals; and (3) temporal sampling would reveal that terrestrial mammal eDNA is detectable for short periods in comparison to fully aquatic vertebrates.

## 2. Materials and methods

### 2.1 Study species

We studied nine mammal species that are the focus of European conservation or management (Mathews et al., 2018): European water vole (*Arvicola amphibius*), European otter (*Lutra lutra*), Eurasian beaver (*Castor fiber*), European hedgehog (*Erinaceus europaeus*), European badger (*Meles meles*), red deer (*Cervus elaphus*), Eurasian lynx (*Lynx lynx*), red squirrel (*Sciurus vulgaris*), and European pine marten (*Martes martes*). Water vole, otter, red squirrel, pine marten and hedgehog are UK Biodiversity Action Plan species (Joint Nature Conservation Committee, 2018). Water vole, otter, and beaver are semi-aquatic, red squirrel and pine marten are arboreal, and the other species are ground-dwelling. Badger and red deer live in groups whereas the other species are predominantly solitary.

### 2.2 Experiment 1: eDNA detection and signal strength in artificial systems

Behavioural observation and eDNA sampling were conducted between 18^th^ – 21^st^ September 2017 at Wildwood Trust (WT), Kent, England, and 10^th^ – 11^th^ October 2017 at Royal Zoological Society of Scotland (RZSS) Highland Wildlife Park (HWP), Kingussie, Scotland. Sixteen categories of behaviour were defined based on potential contact with waterbodies and species lifestyle, and the frequency and duration of behaviours recorded (Table 1, Appendix A: Table A1). The number of individuals in each enclosure was recorded alongside waterbody size (Table 2). Beaver, lynx, red deer, and red squirrel were present at both wildlife parks, whereas other captive species were only present at WT. Each species was observed for one hour on two separate occasions except nocturnal mammals (badger and beaver), which were observed overnight using camera traps (Bushnell Trophy Cam Standard, Bushnell Corporation, KS, USA). One camera trap per enclosure was positioned perpendicular to the ground (1 m height, 2 m from shoreline) to capture water and shoreline. Cameras took 30 s videos (1920 x 1080) when triggered (30 s interval between triggers) at high sensitivity. Behavioural observation was not undertaken for WT water voles as animals were under quarantine or HWP red squirrels as individuals were wild. Photos of waterbodies in animal enclosures are provided in Appendix B.

**Table 1.** Ethogram used to catalogue mammal behaviours that occur in or near artificial waterbodies in captive enclosures. Importantly, this ethogram was designed to catalogue mammal behaviours potentially leading to eDNA deposition. Therefore, it may not be comparable to ethograms typically used in study of captive animals.

| Behaviour | Definition |
| --- | --- |
| Swimming | Mammal completely submerged in and moving through waterbody using limbs |
| Bodypart in water | Mammal partially submerged in waterbody, e.g. foot or tail in water |
| Drinking | Water taken into mouth and swallowed by mammal |
| Feeding | Food taken into mouth and swallowed by mammal in or near waterbody, e.g. otter and fish |
| Scratching | Bodypart or external object in enclosure used by mammal to relieve itch near waterbody |
| Urinating/scent-marking | Liquid excretion passed by mammal in or near waterbody |
| Pooing | Solid excretion passed by mammal in or near waterbody |
| Sniffing | Air visibly drawn through nose of mammal to detect a smell around waterbody, possibly involving contact with water |
| Standing | Mammal motionless in or near waterbody |
| Walking | Mammal moving around waterbody at a regular pace by lifting and setting down each foot in turn, never having both feet off the ground at once |
| Running | Mammal moving around waterbody at a speed faster than a walk, never having both or all the feet on the ground at the same time |
| Vocalising | Mammal producing sound while in or near waterbody |
| Grooming | Mammal cleaning fur or skin with its tongue while in or near waterbody |
| Resting | Mammal lying down or sitting in or near waterbody |
| Other | Behaviour exhibited in or near waterbody that does not conform to other categories, e.g. chasing tail |
| Not visible | Mammal moved to part of enclosure not visible to the observer |

**Table 2.** Summary of focal species studied at wildlife parks and their lifestyle. The number of individuals present and waterbody size in enclosures is provided.

| Site | Species | Lifestyle | Enclosure | Number of individuals | Waterbody size (m<sup>2</sup>) |
| --- | --- | --- | --- | --- | --- |
| Wildwood Trust | European otter<br>(*Lutra lutra*) | Semi-aquatic | 1 | 2 | 162 |
|  | European water vole<br>(*Arvicola amphibius*) | Semi-aquatic | 1 | 4 | 0.09 |
|  |  |  | 2 | 1 | 0.09 |
|  | European beaver<br>(*Castor fiber*) | Semi-aquatic | 1 | 2 | 100 |
|  |  |  | 2 | 1 | 100 |
|  | European hedgehog<br>(*Erinaceus europaeus*) | Ground-dwelling | 1 | 1 | 0.04 |
|  |  |  | 2 | 2 | 0.04 |
|  | European badger<br>(*Meles meles*) | Ground-dwelling | 1 | 4 | 1.73 |
|  | Red deer<br>(*Cervus elaphus*) | Ground-dwelling | 1 | 8 | 100 |
|  | Eurasian lynx<br>(*Lynx lynx*) | Ground-dwelling | 1 | 2 | 2 |
|  | Red squirrel<br>(*Sciurus vulgaris*) | Arboreal | 1 | 2 | 0.01 |
|  |  |  | 2 | 3 | 0.01 |
|  |  |  | 3 | 3 | 0.01 |
|  |  |  | 4 | 2 | 0.01 |
|  | European pine marten<br>(*Martes martes*) | Arboreal | 1 | 1 | 2 |
|  |  |  | 2 | 1 | 0.375 |
| Highland Wildlife Park | Red squirrel<br>(*Sciurus vulgaris*) | Arboreal | NA | NA | 0.25 |
|  | Eurasian lynx<br>(*Lynx lynx*) | Ground-dwelling | 1 | 8 | 2 |
|  | European beaver<br>(*Castor fiber*) | Semi-aquatic | 1 | 2 | 50 |
|  | Red deer<br>(*Cervus elaphus*) | Ground-dwelling | 1 | 30 | NA |

Water samples were collected from enclosures within 3 hrs of the second behavioural observation period. Up to six directed or stratified samples were collected, but sample number varied by species according to waterbody size and observed behaviours (Tables A1, A2). Enclosure drinking containers were also sampled and classed as ‘other’ samples. Bathing and drinking bowls were sampled where enclosures contained no artificial waterbodies (WT water vole, red squirrel, and hedgehog). The HWP beaver enclosure was empty for 24 hrs before sampling. Water was sampled from a RZSS Edinburgh Zoo (EZ) enclosure containing beavers and classed as ‘other’. A sample was collected from a water bath in the HWP woods to capture wild red squirrels and classed as ‘other’.

Directed samples (2 L surface water taken approximately where behaviours were observed) were collected before stratified samples (2 L surface water [8 x 250 ml pooled subsamples] taken at equidistant points [access permitting] around the waterbody perimeter) to minimise disturbance to the water column and cross-contamination risk. Samples were collected using sterile Gosselin™ HDPE plastic bottles (Fisher Scientific UK Ltd, UK) and disposable gloves. A field blank (1 L molecular grade water [MGW]) was taken into each species enclosure, opened, and closed before artificial water sources were sampled. Samples (*n* = 80) collected from WT and HWP were transported alongside field blanks (*n* = 13) in sterile coolboxes with ice packs to the University of Kent (UoK) and EZ respectively, where ice was added to coolboxes.

Samples and blanks were vacuum-filtered within 6 hrs of collection in a UoK wet laboratory and within 24 hrs of collection in an EZ staff room. Surfaces and equipment were sterilised before, during, and after set-up in temporary work areas. Surfaces and vacuum pumps were wiped with 10% v/v chlorine-based commercial bleach (Elliott Hygiene Ltd, UK) solution. Non-electrical equipment was immersed in 10% bleach solution for 10 minutes, followed by 5% v/v MicroSol detergent (Anachem, UK), and rinsed with purified water. Up to 500 ml of each 2 L sample was vacuum-filtered through sterile 0.45 μm mixed cellulose ester membrane filters with pads (47 mm diameter; Whatman, GE Healthcare, UK) using Nalgene™ filtration units. One hour was allowed for each sample to filter and a second filter used if clogging occurred. A filtration blank (1 L MGW) was processed during each filtration round (*n* = 12), and equipment sterilised after each filtration round. After 500 ml had filtered or one hour had passed, filters were removed from pads using sterile tweezers, placed in sterile 47 mm petri dishes (Fisher Scientific UK Ltd, UK), sealed with parafilm (Sigma-Aldrich Company Ltd, UK), and stored at −20 °C. The total water volume filtered per sample was recorded for downstream analysis (Table A2; Fig. A1).

### 2.3 Experiment 2: eDNA detection and signal strength in natural systems

At three sites where focal species were present based on cumulative survey data, we selected two ponds (range 293-5056 m^2^, average 1471 m^2^) within 4 km of each other. The Bamff Estate (BE), Alyth, Scotland, was selected for beaver, otter, badger, red deer, and red squirrel, but roe deer (*Capreolus capreolus*) and red fox (*Vulpes vulpes*) were also present. Otter, water vole, and badger were present at Tophill Low Nature Reserve (TLNR), Driffield, East Yorkshire, alongside American mink (*Neovison vison*), stoat (*Mustela erminea*), weasel (*Mustela nivalis*), rabbit (*Oryctolagus cuniculus*), brown hare (*Lepus europaeus*), red fox, roe deer, and grey squirrel (*Sciurus carolinensis*). We selected Thorne Moors (TM), Doncaster, South Yorkshire, for red deer and badger, but stoat, weasel, red fox, roe deer, and Reeve’s muntjac (*Muntiacus reevesi*) were also present. Camera traps (Bushnell Trophy Cam Standard/Aggressor, Bushnell Corporation, KS, USA) were deployed at TM (one per pond) and BE (three per pond) one week prior to eDNA sampling and collected once sampling was completed. At TLNR, camera traps (two to three per pond) were deployed one day before a 5-day period of eDNA sampling and collected one week after sampling was completed. Camera traps were positioned perpendicular to the ground (1 m height, 0.3-1 m from shoreline) to capture water and shoreline. Cameras took three photographs (5 megapixel) when triggered (3 s interval between triggers) at high sensitivity.

Ten stratified samples were collected from the shoreline of each pond (TM: 17^th^ April 2018; BE: 20^th^ April 2018; TLNR: 23^rd^ – 27^th^ April 2018) and a field blank (1 L MGW) included as in Experiment 1. TLNR ponds were sampled every 24 hrs over 5 days to investigate spatiotemporal variation in mammal eDNA signals. TM and TLNR samples were transported on ice in sterile coolboxes to the University of Hull (UoH) eDNA facility, and stored at 4 °C. BE samples were transported in sterile coolboxes with ice packs to BE accommodation. Surfaces and equipment were sterilised before, during, and after set-up as in Experiment 1. Samples (*n* = 140) and field blanks (*n* = 14) were vacuum-filtered within 4 hrs of collection as in Experiment 1 with minor modifications to maximise detection probability as follows. The full 2 L of each sample was vacuum-filtered where possible, two filters were used for each sample, and duplicate filters were stored in one petri dish at −20 °C. A filtration blank (1 L MGW) was processed during each filtration round (*n* = 21). The total water volume filtered per sample was recorded (Table A3).

### 2.4 DNA extraction

DNA was extracted within 2 weeks of filtration at the UoH eDNA facility using the Mu-DNA water protocol (Sellers, Di Muri, Gómez, & Hänfling, 2018). The full protocol is available at: https://doi.org/10.17504/protocols.io.qn9dvh6. Duplicate filters from samples in Experiment 1 were lysed independently and the lysate from each loaded onto one spin column. As more samples were collected in Experiment 2, duplicate filters were co-extracted by placing both in a single tube for bead milling. An extraction blank, consisting only of extraction buffers, was included for each round of DNA extraction (*n* = 17). Eluted DNA (100 μl) was stored at −20 °C until PCR amplification.

### 2.5 eDNA metabarcoding

Our eDNA metabarcoding workflow is fully described in Appendix A. Briefly, we performed nested metabarcoding using a two-step PCR protocol, where Multiplex Identification (MID) tags were included in the first and second PCR for sample identification (Kitson et al., 2019). The first PCR amplified eDNA in triplicate with published 12S ribosomal RNA (rRNA) primers 12S-V5-F (5’-ACTGGGATTAGATACCCC-3’) and 12S-V5-R (5’-TAGAACAGGCTCCTCTAG-3’) (Riaz et al., 2011). Harper et al. (2018) validated these primers *in silico* for UK vertebrates, and found 91/112 mammal species listed on the Natural History Museum Checklist of Mammalia v1 (subspecies excluded) could be distinguished. Nine indistinguishable species lacked reference sequences, whereas 12 had reference sequences but did not amplify. PCR positive controls (two per PCR plate; *n* = 16) were exotic cichlid (*Maylandia zebra*) DNA (0.05 ng/µl), and PCR negative controls (two per PCR plate; *n* = 16) were MGW (Fisher Scientific UK Ltd, UK). PCR products were pooled to create sub-libraries (Fig. A2) and purified with Mag-BIND**^®^** RxnPure Plus magnetic beads (Omega Bio-tek Inc, GA, USA), following the double size selection protocol established by Bronner et al. (2009). Ratios of 0.9x and 0.15x magnetic beads to 100 μL of each sub-library were used. Eluted DNA (30 μL) was stored at −20 °C until the second PCR could be performed. The second PCR bound pre-adapters, MID tags, and Illumina adapters to the sub-libraries. PCR products were purified with Mag-BIND**^®^** RxnPure Plus magnetic beads (Omega Bio-tek Inc, GA, USA), following the double size selection protocol established by Bronner et al. (2009). Ratios of 0.7x and 0.15x magnetic beads to 50 μL of each sub-library were used. Eluted DNA (30 μL) was stored at 4 °C until quantification and normalisation. The library was purified again, quantified by qPCR using the NEBNext**^®^** Library Quant Kit for Illumina**^®^** (New England Biolabs**^®^** Inc., MA, USA), and fragment size (330 bp) and removal of secondary product verified using an Agilent 2200 TapeStation and High Sensitivity D1000 ScreenTape (Agilent Technologies, CA, USA). The library (220 eDNA samples, 27 field blanks, 33 filtration blanks, 17 extraction blanks, 16 PCR negative controls, and 16 PCR positive controls) was sequenced on an Illumina MiSeq® using a MiSeq Reagent Kit v3 (600-cycle) (Illumina, Inc, CA, USA). Raw sequence reads were demultiplexed using a custom Python script. metaBEAT v0.97.11 (https://github.com/HullUni-bioinformatics/metaBEAT) was used for quality trimming, merging, chimera removal, clustering, and taxonomic assignment of sequences against our UK vertebrate reference database (Harper et al., 2018) which contains sequences for 103 UK mammals. Taxonomic assignment used a lowest common ancestor approach based on the top 10% BLAST matches for any query that matched a reference sequence across more than 80% of its length at minimum identity of 98%.

### 2.6 Data analysis

Analyses were performed in R v.3.4.3 (R Core Team, 2017). The total unrefined read counts (i.e. raw taxonomically assigned reads) per sample were calculated and retained for downstream analyses. Assignments were corrected: family and genera containing a single UK species were reassigned to that species, species were reassigned to domestic subspecies, and misassignments were corrected, e.g. *Lynx pardinus* and *Lynx lynx*. Manual reassignment duplicated some metaBEAT assignments thus the read count data for these assignments were merged. Taxon-specific sequence thresholds (i.e. maximum sequence frequency of each taxon in PCR positive controls) were used to mitigate cross-contamination and false positives (Table A4, Fig. A3), and remnant contaminants and higher taxonomic assignments removed excluding the following genera. *Anas* (Dabbling ducks) was retained because potential for hybridisation reduced confidence in species-level assignments, and *Emberiza* (Buntings) and *Larus* (White-headed gulls) were retained because reference sequences were missing for several common species. Dataset refinement is fully described in Appendix A. Taxonomic assignments remaining in the refined dataset were predominantly of species resolution and considered true positives. We split the refined dataset by Experiment 1 (artificial waterbodies) and Experiment 2 (natural ponds). Proportional read counts for each species were calculated from the total unrefined read counts per sample. Our proportional read count data were not normally distributed (Shapiro–Wilk normality test: *W* = 0.915, *P* < 0.001), thus we used a Mann-Whitney *U* test to compare the median proportional read count of stratified and directed samples across species.

We employed binomial Generalized Linear Mixed-effects Models (GLMMs) with the logit link function using the package glmmTMB (development version; Brooks et al., 2017) for the following tests. First, we compared the eDNA signals from stratified and directed samples for each mammal species using a hierarchical model including sample type nested within species (fixed) and wildlife park (random) as effects. We tested the influence of species lifestyle on mammal eDNA signals using a model with species lifestyle (fixed) and species nested within wildlife park (random) as effects. Using directed samples, we tested the influence of behaviour on mammal eDNA signals using two hierarchical models, including species nested within wildlife park (random) and specific (e.g. swimming, drinking) or generic (i.e. water contact versus no water contact) behaviour(s) respectively (fixed) as effects. We assessed model fit using diagnostic plots and performed validation checks to ensure model assumptions were met and overdispersion was absent (Zuur, Ieno, Walker, Saveliev, & Smith, 2009).

For Experiment 2, we qualitatively compared mammal presence-absence records generated by eDNA metabarcoding, camera trapping, and field signs. TLNR ponds were sampled every 24 hrs for 5 days, thus proportional read counts were averaged across days for comparison to BE and TM ponds (sampled once each). We qualitatively compared the distribution and persistence of eDNA signals between semi-aquatic and terrestrial mammals using tile plots and heat maps of the unaveraged proportional read counts for identified species at TLNR over the 5-day period. All figures were produced using the package ggplot2 v3.0.0 (Wickham, 2016).

## 3. Results

### 3.1 eDNA metabarcoding

The sequencing run generated 47,713,656 raw sequence reads, of which 37,590,828 remained following trimming, merging, and length filter application. After removal of chimeras and redundancy via clustering, the library contained 21,127,061 sequences (average read count of 64,215 per sample including controls), of which 16,787,750 (79.46%) were assigned a taxonomic rank. Contamination (Fig. A4) was observed in the field blanks (badger, beaver, lynx, pine marten, red squirrel, and water vole) as well as in the filtration and extraction blanks (human [*Homo sapiens*] and cichlid). PCR negative controls were contaminated to different extents with human, cichlid, beaver, and pine marten as well as non-focal species. After threshold application, contaminants remaining in eDNA samples included Gentoo penguin (*Pygoscelis papua*), reindeer (*Rangifer tarandus*), cichlid, and human. The refined dataset contained 59 vertebrate species, including six amphibians, 10 fish, 19 birds, and 24 mammals (Table A5).

### 3.2 Experiment 1: eDNA detection and signal strength in artificial systems

All nine focal species were detected in captivity, of which seven were detected in all water samples taken from their respective enclosures. HWP red deer were not detected in 2 of 5 stratified samples, and WT hedgehog was not detected in 1 of 2 drinking bowl samples (Fig. 1). ‘Other’ samples (neither directed nor stratified) were excluded from further comparisons, thus hedgehog, red squirrel, and water vole were omitted in downstream analyses. Across species, stratified samples (0.406) had a higher median proportional read count than directed samples (0.373), but this difference was not significant (Mann-Whitney *U* test: *U* = 1181.5, *P* = 0.829). Proportional read counts for directed and stratified samples did not significantly differ (𝜒^2^_6_ = 0.364, *P =* 0.999) within species either (Fig. 2a; GLMM: θ = 0.168, 𝜒^2^_53_ = 8.915, *P* = 1.000, pseudo-*R*^2^ = 39.21%). Otter proportional read counts were lower than other species, but not significantly so. Similarly, species lifestyle (semi-aquatic, ground-dwelling, arboreal) did not influence (𝜒^2^_2_ = 0.655, *P* = 0.721) proportional read counts (Fig. 2b; GLMM: θ = 0.213, 𝜒^2^_61_ = 13.002, *P* = 1.000, pseudo-*R*^2^ = 11.85%). Proportional read counts did not differ (𝜒^2^_11_ = 1.369, *P* = 0.999) according to specific behaviours exhibited by species (Fig. 3a; GLMM: θ = 0.355, 𝜒^2^_31_ = 11.013, *P* = 0.999, pseudo-*R*^2^ = 9.17%). Likewise, generic behaviour (i.e. water contact versus no water contact) did not influence (𝜒^2^_11_ = 0.002, *P* = 0.964) proportional read counts (Fig. 3b; GLMM: θ = 0.217, 𝜒^2^_41_ = 8.897, *P* = 1.000, pseudo-*R*^2^ = 8.50%).

**Figure 1.**
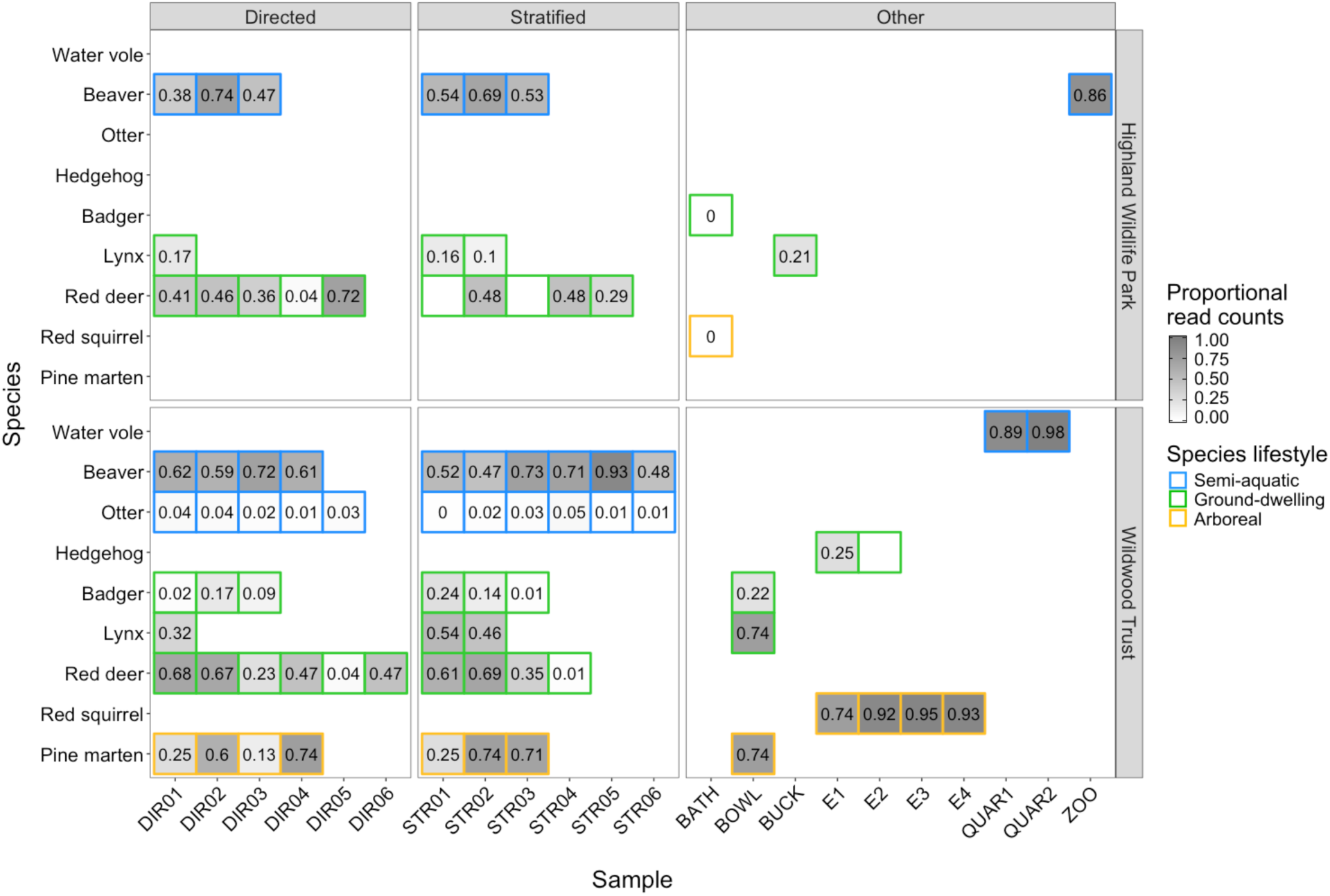
Heatmap showing proportional read counts for eDNA samples (*n* = 81) from Experiment 1. The heatmap is faceted by sample type (directed, stratified or other) and wildlife park (Highland Wildlife Park or Wildwood Trust). Each cell represents an individual sample taken from an enclosure containing the focal species in that row. Directed (DIR01-DIR06) and stratified (STR01-STR06) samples were collected for each species from artificial waterbodies. Samples were also collected from drinking containers (E1, E2, E3, E4, BOWL, BUCK), water vole (QUAR1, QUAR2) and RZSS Edinburgh Zoo beaver (ZOO) enclosures, and a water bath (BATH) in RZSS Highland Wildlife Park woods. The maximum proportional read count for each cell (i.e. sample) is 1, if all reads from a particular sample belonged to the focal species. Cells containing 0 represent samples with proportional read counts less than 0.01 whereas empty cells are samples with proportional read counts of exactly 0.

**Figure 2.**
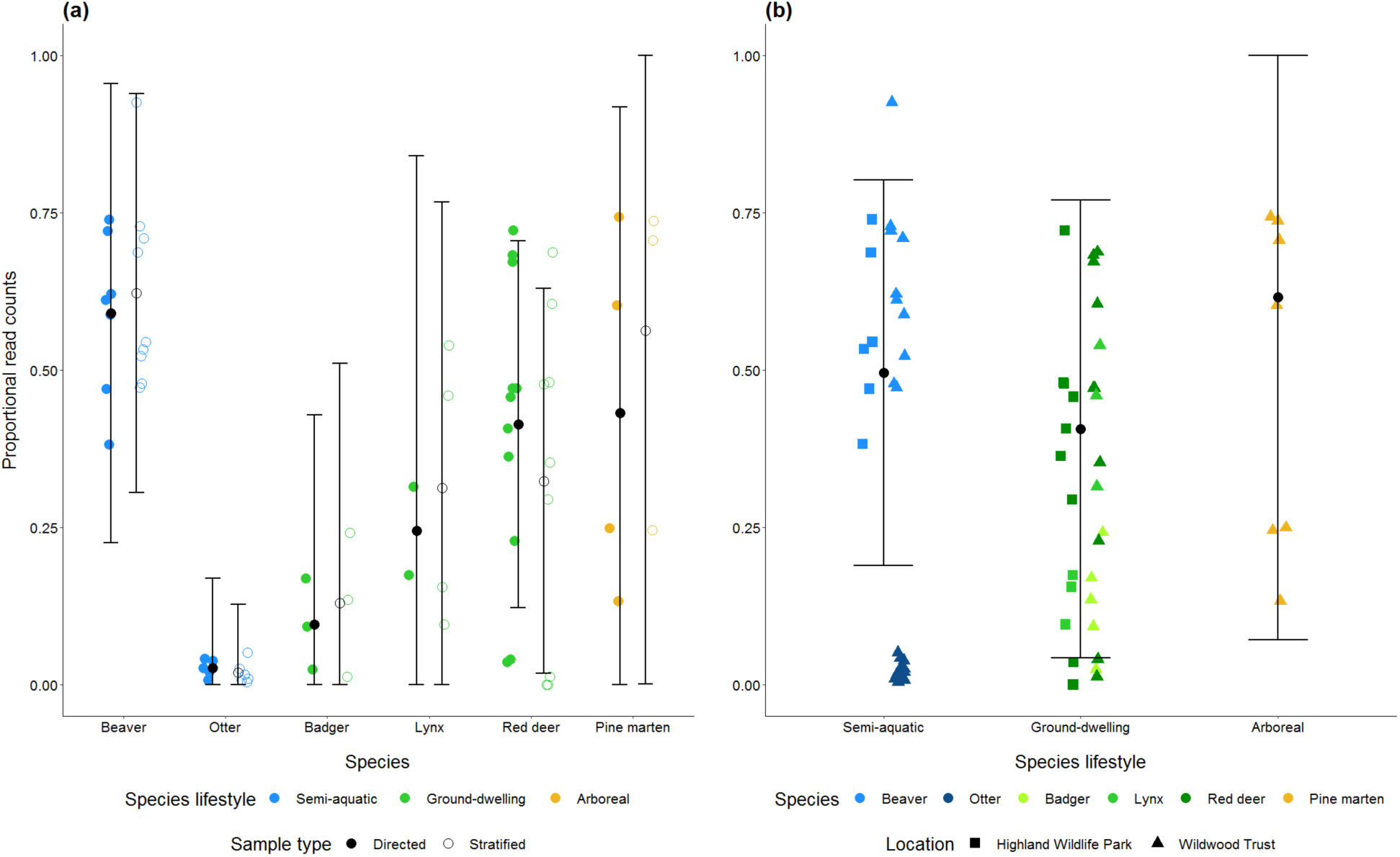
Relationships predicted by the binomial GLMMs between proportional read counts and sample type nested within species **(a)** or species lifestyle **(b)** for Experiment 1. The observed data (coloured points) are displayed against the predicted relationships (black points with error bars) for each species **(a)** or species lifestyle **(b)**. Points are shaped by sample type **(a)** or wildlife park **(b)**, and coloured by species lifestyle. Error bars represent the standard error around the predicted means.

**Figure 3.**
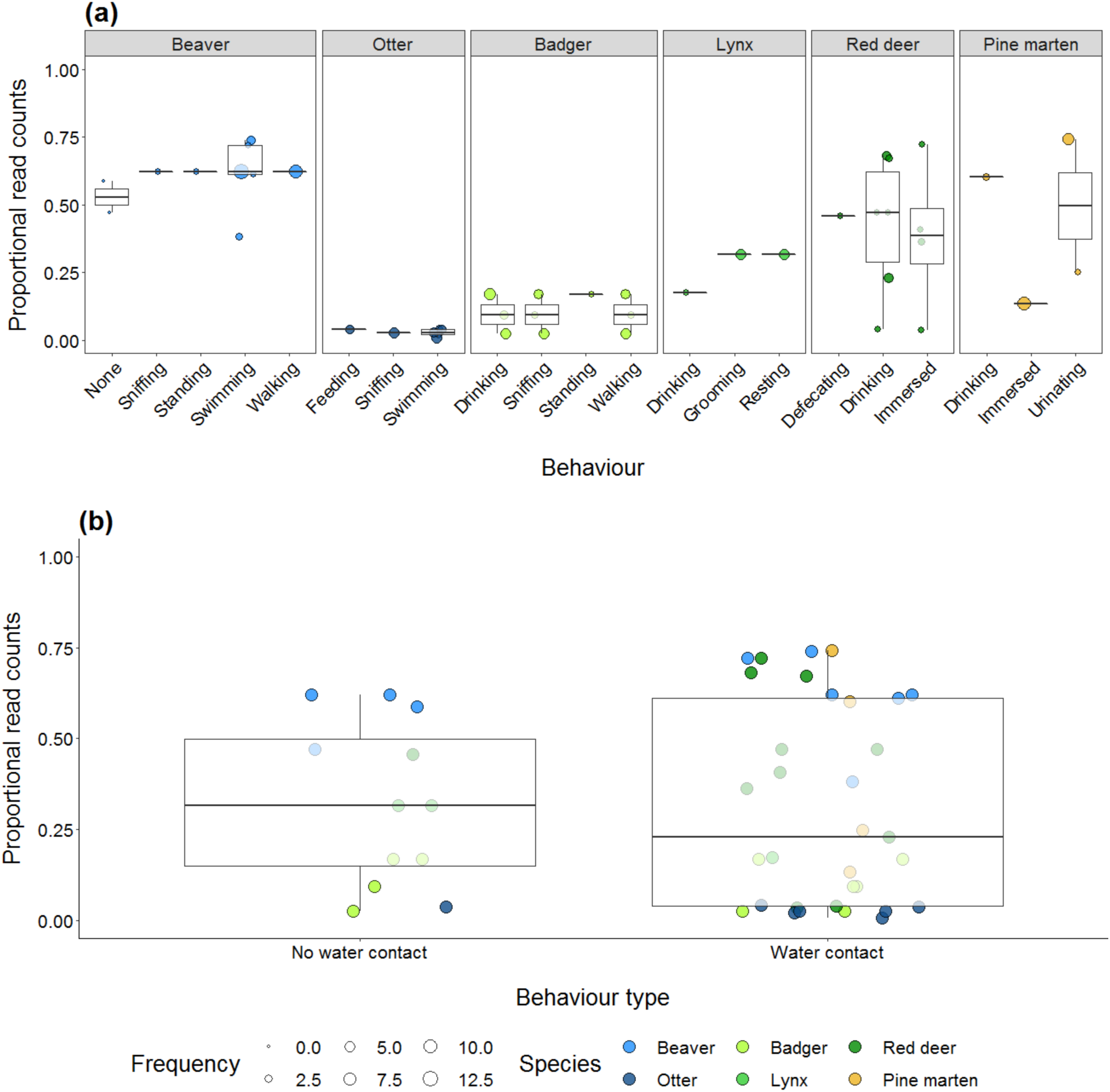
Boxplots showing the mean proportional read counts for specific **(a)** and generic **(b)** behaviour(s) exhibited by focal species in Experiment 1. Boxes show 25th, 50th, and 75th percentiles, and whiskers show 5th and 95th percentiles. Points are coloured by species lifestyle, and each point in **(a)** represents a directed sample sized by frequency of behaviour. The behaviour ‘none’ for beaver represents occurrences of beaver in water but out of view of camera traps.

### 3.3 Experiment 2: eDNA detection and signal strength in natural systems

At natural ponds, eDNA metabarcoding, camera trapping, and field signs all detected beaver, red deer, and roe deer. Camera traps (Fig. 4) and field signs recorded red fox and badger when eDNA metabarcoding did not (Fig. 5). However, eDNA metabarcoding revealed small mammals missed by cameras and field signs, including water vole, water shrew (*Neomys fodiens*), bank vole (*Myodes glareolus*), common shrew (*Sorex araneus*), brown rat (*Rattus norvegicus*), rabbit, grey squirrel, and common pipistrelle (*Pipistrellus pipistrellus*). We observed mice or vole footprints at BE Pond 1, but could not ascertain species. Fig. 5 summarises mammals recorded by different methods at each site with reference to cumulative survey data. Notably, only beaver was found at the same ponds by all methods. Although methods shared species at site level, species were not always detected at the same pond. Detection rates for species captured by at least one survey method are summarised in Table A6.

**Figure 4.**
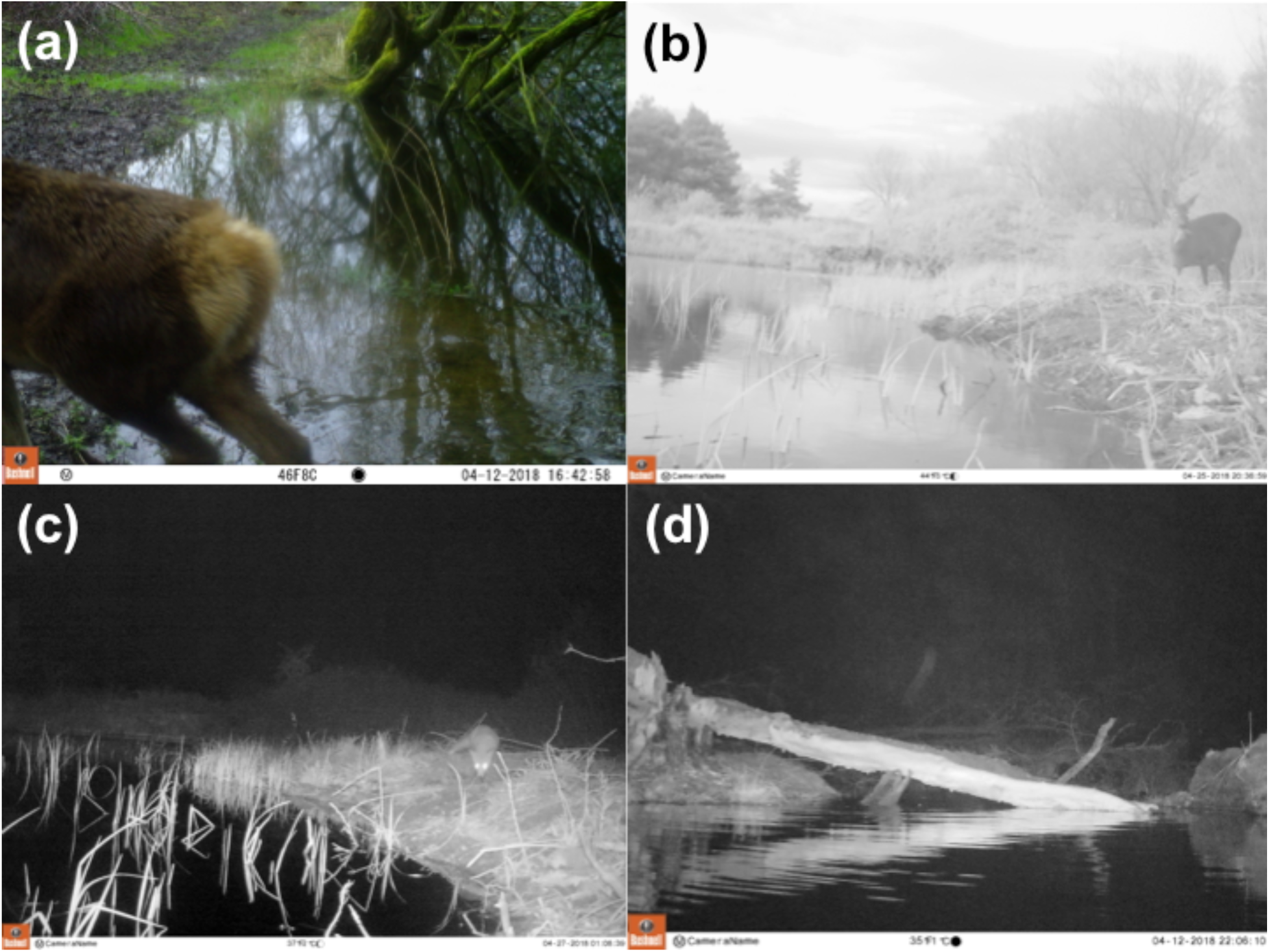
Exemplar camera trap photos taken at natural ponds where focal species were present in Experiment 2. Red deer was recorded at Thorne Moors **(a)**, roe deer **(b)** and red fox **(c)** were recorded at Tophill Low Nature Reserve, and beaver was recorded at the Bamff Estate **(d)**.

**Figure 5.**
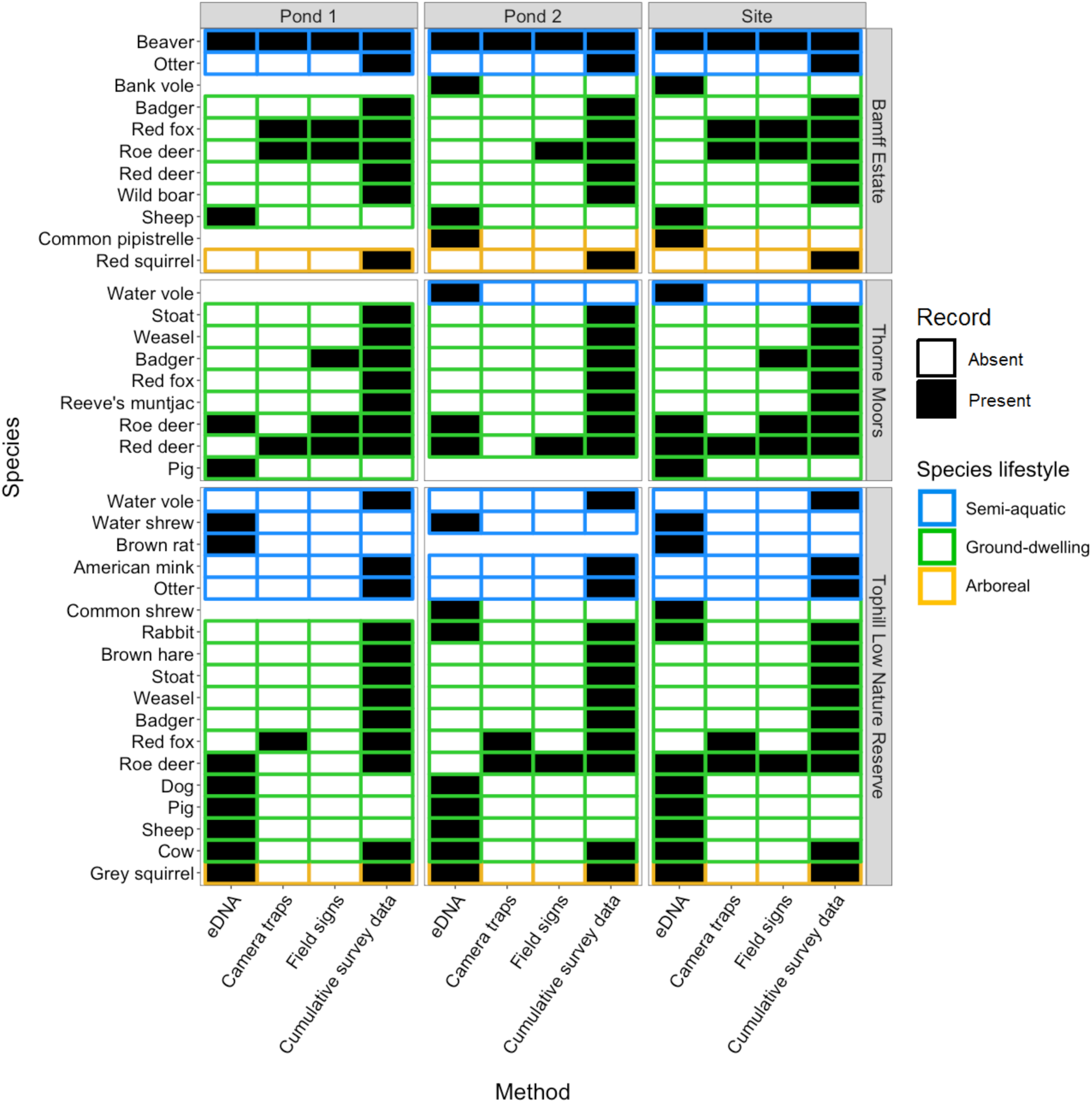
Tile plot showing species presence-absence at individual pond and site-level as indicated by field signs, camera trapping, and eDNA metabarcoding in Experiment 2. Surveys were performed at sites where focal species presence was confirmed by cumulative survey data.

Sampling of natural ponds revealed spatial patterns in eDNA detection and signal strength. eDNA from non-domestic terrestrial mammals (i.e. mammals excluding dog [*Canis lupus familiaris*], pig [*Sus scrofa domesticus*], sheep [*Ovis aries*] and cow [*Bos taurus*]) was unevenly dispersed compared with semi-aquatic mammals (Fig. A5). Semi-aquatic beaver and water vole were detected in at least 90% and 60% respectively of water samples (*n* = 10) collected from single ponds, albeit water shrew was only detected in 10% of samples. Non-domestic terrestrial mammals were routinely detected in <20% of water samples collected from a pond and left relatively weak eDNA signals. Overall, beaver was the most consistently detected mammal with the highest proportional read counts. However, the strongest and most evenly distributed signals belonged to amphibians, particularly common frog (*Rana temporaria*) and great crested newt (*Triturus cristatus*) (Fig. A5).

TLNR samples collected over a 5-day period (D01-05) revealed that mammal detection heavily depends on the spatial and temporal resolution of eDNA metabarcoding surveys (Fig. A6). Mammal eDNA signals in pond water were ephemeral, often disappearing within 24-48 hrs of initial detection, as opposed to amphibians that were detected for multiple days and whose eDNA signal increased in strength. The majority of semi-aquatic or terrestrial mammals were only detected in a single sample on each day.

## 4. Discussion

We have demonstrated the potential of eDNA metabarcoding for monitoring conservation and management priority mammals, but species detection rates are variable. Our experiments have validated this molecular approach and provided new insights that will inform the development and application of mammal eDNA metabarcoding. Sampling strategy, mammal lifestyle, and mammal behaviour did not influence eDNA detection and signal strength in captivity, but all played vital roles in natural ponds. Although semi-aquatic and terrestrial mammals were detected from pond water, their eDNA signals were temporary and weak in comparison to aquatic amphibians and fishes. Nonetheless, this suggests that eDNA is representative of contemporary and local mammal diversity.

### 4.1 Influence of sampling strategy and mammal behaviour on eDNA detection

In Experiment 1, all nine focal species were detected in captivity, and seven were detected in all water samples taken from their respective enclosures. This demonstrates that our method can successfully detect a variety of mammals from pond and drinking water. Surprisingly, we found that neither sampling strategy nor mammal lifestyle nor mammal behaviour influenced eDNA detectability and signal strength in captivity. This included behaviours associated with eDNA deposition, e.g. swimming, drinking, urination, and defecation (Rodgers & Mock, 2015; Ushio et al., 2017; Williams et al., 2018). Enclosures were permanently occupied and artificial waterbodies likely saturated with eDNA, which possibly masked behavioural signals. Modest replication may have limited experimental power, preventing patterns being detected statistically. Nonetheless, our results show that mammal contact with water enables eDNA deposition and detection.

Unsurprisingly, given the nature of wild mammal interactions with natural systems versus those in captivity, Experiment 2 results highlight the challenges of mammal eDNA detection. We recorded 17 mammals using three monitoring tools, comparable to the 17 mammals expected from cumulative survey data despite discordance. Field signs and camera trapping detected red fox and badger where eDNA metabarcoding did not, but eDNA metabarcoding identified water vole and other small mammals missed on camera or with ambiguous field signs, i.e. mice, voles, shrews. Importantly, camera trap deployment period, height, and positioning may have influenced small mammal detection by this method (Caravaggi et al., 2018). Ishige et al. (2017) achieved comparable mammal detection at salt licks with eDNA metabarcoding and camera trapping, but species presence was inconsistent between salt licks surveyed. Using multi-species occupancy modelling for three mammal species, Sales et al. (2019) observed water-based eDNA metabarcoding provided comparable detection probabilities to conventional survey methods and actually outperformed camera trapping. Similarly, Leempoel et al. (2019) found soil-based eDNA metabarcoding identified the same mammals as camera trapping as well as small mammals rarely seen on camera, albeit the methods differed between sites. Our own results echo all three studies, where despite some inconsistencies, eDNA metabarcoding enhanced species inventories and identified smaller, cryptic taxa.

Notably, no survey method captured semi-aquatic otter despite presence at study sites and successful detection in eDNA metabarcoding studies of UK ponds (Harper et al., 2019), lakes (Hänfling et al., 2017), and rivers/streams (Sales et al., 2019). Captive otter also had a weaker eDNA signal than other semi-aquatic mammals studied here. Lower eDNA detection rates for otter, badger, and red fox may stem from species’ ecologies (Sales et al., 2019). These mammals are wide-ranging (Gaughran et al., 2018; Thomsen et al., 2012) and may not readily release DNA in water. Otters often spraint on grass or rock substrata outside water and use latrines associated with caves and dens (Ruiz-Olmo & Gosálbez, 1997). As terrestrial mammals, red fox and badger must drink from or enter ponds for eDNA deposition to occur (Rodgers & Mock, 2015; Ushio et al., 2017; Williams et al., 2018). Otter, badger, and red fox detection may require greater spatiotemporal resolution of eDNA sampling. This is reinforced by other eDNA metabarcoding studies where mammal detection was highly variable across sites surveyed (Ishige et al., 2017; Klymus et al., 2017; Leempoel et al., 2019; Sales et al., 2019; Ushio et al., 2017). False negatives may instead be symptomatic of metabarcoding bias, but this is unlikely in our study (section 4.2).

eDNA from other semi-aquatic mammals was evenly distributed, being found in most or all samples collected on fine spatial scales within natural ponds, whereas terrestrial mammal eDNA was highly localised and detected in few (<20%) samples. Mammal eDNA signals varied temporally, being detectable for two consecutive days maximum. Depending on the species, mammal eDNA may be spatially and temporally clumped in lentic ecosystems due to the nature and frequency of water contact. Unless non-domestic mammals exhibit behaviours involving prolonged water contact (e.g. swimming, wallowing), they may only be detected at drinking sites (Klymus et al., 2017; Ushio et al., 2017; Williams et al., 2018). Conversely, domestic mammals may have elevated detection rates in ponds due to high occurrence of these waterbodies in agricultural landscapes as well as eDNA transport by rainfall and run-off (Staley et al., 2018). eDNA detection and persistence are further influenced by group size, where eDNA from multiple individuals endures for longer periods in water than eDNA from single individuals (Williams et al., 2018). Detailed investigations incorporating biotic (e.g. population size, body mass, behaviour) and abiotic (e.g. temperature, pH, rainfall) factors are needed to understand the longevity of mammal eDNA signals in aquatic ecosystems (Rodgers & Mock, 2015; Sales et al., 2019; Williams et al., 2018).

Our two experiments have shown that sampling strategy influences mammal eDNA detection. Mammal eDNA was evenly distributed in closed, artificial waterbodies, but locally distributed in open, natural ponds. Captive mammal enclosures contained one species (excluding HWP red deer) and a drinking container(s) and/or small waterbody (range 0.01-162 m^2^, mean 27.4 m^2^). Some enclosures housed more individuals of a species than others, thereby increasing eDNA deposition and detection probability (Williams et al., 2018). Wild mammals have an array of freshwater habitats at their disposal and can hold vast territories. Therefore, rates of pond visitation and eDNA deposition are more irregular (Klymus et al., 2017; Ushio et al., 2017), possibly leading to between-sample variation (Williams et al., 2018).

### 4.2 Accounting for false positives and false negatives in metabarcoding

eDNA metabarcoding has potential for inclusion in mammal monitoring schemes (section 4.3), but like existing monitoring tools, may produce false negatives or false positives. Our process controls identified low-level contamination at all stages of metabarcoding, but primarily during sampling or PCR (Appendix A). We applied taxon-specific sequence thresholds to our data to mitigate false positives as in Harper et al. (2019). Remnant contaminants were cichlid (laboratory), Gentoo penguin (environment), reindeer (environment), and human (environment/laboratory). Gentoo penguin is housed at EZ and was identified from EZ beaver enclosure water. The WT red squirrel and reindeer enclosures are in close proximity. DNA transport by wildlife (e.g. waterfowl [Hänfling et al., 2016]) and park staff/visitors may explain this environmental contamination. Human DNA was present across process controls corresponding to artificial and natural waterbodies. Human DNA may be amplified and sequenced instead of focal species, potentially resulting in false negative detections for rare and/or less abundant species. Human DNA blocking primers can prevent this bias, but may impair PCR amplification efficiency (Klymus et al., 2017; Ushio et al., 2017; Valentini et al., 2016). Sequence thresholds are one method of accounting for contamination in metabarcoding datasets, but this is a topic that warrants deeper investigation aimed at researching and refining standardised methods for false positive identification and mitigation, e.g. the R package microDecon (McKnight et al., 2019).

In our study, eDNA metabarcoding produced false negatives for otter, badger, and red fox at natural ponds. We selected a 12S metabarcode designed to amplify vertebrate DNA (Riaz et al., 2011). One of four fox reference sequences (NCBI Accession: KF387633.1) possessed one mismatch to the forward primer, and one of three otter reference sequences (NCBI Accession: EF672696.1) possessed one mismatch to the reverse primer. These mismatches did not occur within the first or last four bases of either primer sequence, and there were no primer mismatches with the badger reference sequences (Harper et al., 2018). Therefore, amplification bias was not responsible for these false negatives. DNA from aquatic and more abundant species may have overwhelmed otter, badger, and red fox DNA during amplification and sequencing, i.e. species-masking (Kelly, Port, Yamahara, & Crowder, 2014; Klymus et al., 2017). Species-masking may also arise from use of proportional read counts as an index of eDNA signal strength. High proportional read counts for a species may translate to a weak eDNA signal if the total mammalian eDNA concentration is highly variable between samples or lower than the total eDNA concentration for other taxonomic groups in a sample. Metabarcoding primers targeting mammals (Ushio et al., 2017) or multi-marker (e.g. 12S, 16S, COI) investigations (Evans et al., 2017; Hänfling et al., 2016; Kelly et al., 2014; Klymus et al., 2017) may improve mammal detection in systems with competition from non-target aquatic species and where total mammalian eDNA concentration varies between samples. Similarly, more biological and technical replication may improve species detection probabilities (Evans et al., 2017; Lawson Handley et al., 2019; Sales et al., 2019; Valentini et al., 2016). Importantly, otter also had lower qPCR detection than amphibians and fish (Thomsen et al., 2012). A metabarcoding and qPCR comparison (e.g. Harper et al., 2018; Lacoursière-Roussel, Dubois, Normandeau, & Bernatchez, 2016) would confirm whether poor amplification efficiency for otter arises from technical bias or species ecology, and whether eDNA metabarcoding can reliably monitor otter alongside the wider mammalian community.

### 4.3 Scope of eDNA metabarcoding for mammal monitoring

Mammal population assessments are hindered by lack of data and systematic monitoring for many species (Mathews et al., 2018). Distribution and occupancy data are poor for most species, with ongoing survey effort biased toward rare species. Surveys heavily rely on citizen science and casual records (Massimino et al., 2018). Tools that provide standardised, systematic monitoring of mammal populations are needed (Mathews et al., 2018). Despite issues inherent to metabarcoding for biodiversity monitoring (Deiner et al., 2017), this tool has enormous potential to enhance mammal monitoring, conservation, and management. eDNA metabarcoding generates distribution data for multiple species, whether rare, invasive, or abundant, and could track conflicting species simultaneously, e.g. water vole, American mink, and otter (Bonesi & Macdonald, 2004) or red squirrel, grey squirrel, and pine marten (Sheehy, Sutherland, O’Reilly, & Lambin, 2018).

eDNA metabarcoding can rapidly survey multitudes of aquatic sites at landscape-scale where camera traps might be resource-intensive, cost-inefficient, and susceptible to theft/damage (Ushio et al., 2017). Field signs require volunteer time and skill (Sadlier et al., 2004) to be employed at comparable spatial scales to eDNA metabarcoding which could provide accurate data for species misidentified from field signs, e.g. mice and voles, otter and mink (Franklin et al., 2019; Harris & Yalden, 2004). However, camera traps and field signs both recorded species that eDNA metabarcoding missed. Therefore, eDNA metabarcoding is complementary and should be incorporated into, not replace, existing monitoring schemes (Leempoel et al., 2019; Sales et al., 2019). This tool could be most effective in mammal monitoring if deployed at the edges of known species distributions, in areas where species presence is unknown, and in areas with isolated species records (Mathews et al., 2018).

### 4.4 Recommendations for mammal survey using eDNA metabarcoding

Water-based eDNA metabarcoding shows great promise for mammal monitoring encompassing conservation and management priority species (Sales et al., 2019). However, there are factors to be considered when designing and conducting mammal eDNA surveys that may not be problematic for surveys of fishes or amphibians. Mammal eDNA detection probabilities from natural ponds will likely be high when areas with dense populations are studied, but rigorous sampling strategies will be required to track mammals in areas sparsely populated by individuals. Multiple ponds must be sampled repeatedly, and samples taken at multiple locations within ponds without pooling to enable site occupancy inferences. Importantly, we sampled natural ponds in spring but sampling in other seasons may produce different results, reflective of species’ ecologies (Lawson Handley et al., 2019). To account for differential mammal visitation rates and maximise eDNA detection probabilities, we recommend that researchers and practitioners using eDNA metabarcoding for mammal monitoring channel their efforts into extensive sampling of numerous waterbodies in a given area over prolonged timescales. Water-based eDNA appears to be indicative of contemporary mammal presence, with most mammal eDNA signals lost within 1-2 days. Therefore, eDNA metabarcoding could provide valuable mammalian community “snapshots” that may not be obtained with other survey methods (Ushio et al., 2017). Different sample types (e.g. water, soil, snow, salt licks, feeding traces, faeces, hair, and blood meals) may also offer new insights to mammal biodiversity (Franklin et al., 2019; Ishige et al., 2017; Kinoshita et al., 2019; Leempoel et al., 2019; Sales et al., 2019; Tessler et al., 2018; Ushio et al., 2017).

## Data accessibility

Raw sequence reads have been archived on the NCBI Sequence Read Archive (Study: SRP164740; BioProject: PRJNA495011; BioSamples: SAMN10195928 - SAMN10196255; SRA accessions: SRR7986451 - SRR7986778). Jupyter notebooks, R scripts and corresponding data are archived online (https://doi.org/10.5281/zenodo.2561415).

## Acknowledgements

The authors declare that they have no conflicts of interest in presenting this work for publication. This work was funded by the University of Hull, and D.S.R was supported by the Natural Environment Research Council award number NE/R016429/1 as part of the UK-SCAPE programme delivering National Capability. We would like to thank Gill Murray-Dickson (National Museums Scotland) and Helen Senn (RZSS Edinburgh Zoo) for feedback on the study design, and Richard Griffiths (DICE, University of Kent) for support with filtration of water samples from Wildwood Trust. We are grateful to staff at Wildwood Trust and RZSS Highland Wildlife Park for assisting with eDNA sampling from animal enclosures. We thank Richard Hampshire and volunteers at Tophill Low Nature Reserve for assisting with camera trap deployment. Tim Kohler (Natural England) and Paul Ramsay kindly gave permission to sample at Thorne Moors and the Bamff Estate respectively.

## Author contributions

L.R.H, B.H, and L.L.H conceived and designed the study. A.I.C and M.G coordinated sampling at Wildwood Trust and RZSS Highland Wildlife Park respectively. L.R.H, C.D.M, C.J.M, and T.L collected and filtered water samples. A.L, T.L, and T.B helped select natural ponds to be surveyed using eDNA, camera trapping, and field signs, and provided camera traps for the study. L.R.H, A.L, and T.L deployed camera traps, which were then collected and footage analysed by L.R.H. L.R.H processed samples in the laboratory with advice from C.D.M and A.M. D.S.R sequenced the final library. L.R.H completed bioinformatic processing of samples, and subsequent data analysis. L.R.H wrote the manuscript, which all authors contributed critically to drafts of and gave final approval for publication.

## Notes

#### Summary of Updates

This corresponding author's contact information has been updated.

