## Appendix A for "Environmental DNA (eDNA) metabarcoding of pond water as a tool to survey conservation and management priority mammals"

### Contents

1. Materials and methods 3
- 1.1 eDNA metabarcoding workflow 3
- 1.2 Data analysis 5
  - 1.2.1 Dataset refinement 5
  - 1.2.2 Effect of water volume and filtration method 7

2. Results and discussion 8
- 2.1 Effect of water volume and filtration method 8
- 2.2 Technical pitfalls of eDNA metabarcoding for mammal monitoring 8

3. Tables 9
- Table A1 9
- Table A2 13
- Table A3 15
- Table A4 21
- Table A5 22
- Table A6 25

4. Figures 27
- Figure A1 27
- Figure A2 28
- Figure A3 29
- Figure A4 30
- Figure A5 31
- Figure A6 32

References 33

### 1. Materials and methods

#### 1.1 eDNA metabarcoding workflow

A two-step PCR protocol was performed on eDNA samples at the University of Hull. Dedicated rooms were available for pre-PCR and post-PCR processes. Pre-PCR processes were performed in a dedicated eDNA laboratory, with separate rooms for filtration, DNA extraction, and PCR preparation of sensitive environmental samples. PCR reactions were set up in a ultraviolet (UV) and bleach sterilised laminar flow hood. To minimise cross-contamination risk between samples, eight-strip PCR tubes with individually attached lids were used instead of 96-well plates (Port et al., 2016) and PCR reactions were sealed with mineral oil (Sigma-Aldrich Company Ltd, Dorset, UK) droplets (Harper et al., 2018). PCR positive ($n$ = 2) and negative controls ($n$ = 2) were included on each PCR run to screen for sources of potential contamination. The DNA (0.05 ng/µL) used for the PCR positive control ($n$ = 16) was *Maylandia zebra* as this is an exotic cichlid not found in UK freshwater habitats. The negative controls ($n$ = 16) substituted molecular grade sterile water (Fisher Scientific UK Ltd, Loughborough, UK) for template DNA.

During the first PCR, the target region was amplified using published 12S rRNA primers 12S-V5-F (5'-ACTGGGATTAGATACCCC-3') and 12S-V5-R (5'-TAGAACAGGCTCCTCTAG-3') (Riaz et al., 2011) that were validated *in silico* for all UK vertebrates by Harper *et al*. (2018). Primers were modified to include MID tags, heterogeneity spacers, sequencing primers, and pre-adapters. During the first PCR, three replicates were performed for each sample to combat amplification bias. PCR reactions were performed in 25 µL volumes, consisting of 3 µL of template DNA, 1.5 µL of each 10 µM primer (Integrated DNA Technologies, Belgium), 12.5 µL of Q5® High-Fidelity 2x Master Mix (New England Biolabs® Inc., MA, USA) and 6.5 µL molecular grade water (Fisher Scientific UK Ltd, UK). PCR was performed on an Applied Biosystems® Veriti Thermal Cycler (Life Technologies, CA, USA) with the following thermocycling profile: 98 °C for 5 mins, 35 cycles of 98 °C for 10 s, 58 °C for 20 s and 72 °C for 30 s, followed by a final elongation step at 72 °C for 7 mins. PCR products were stored at 4 °C until replicates for each sample were pooled, and 2 µL of pooled PCR product was added to 0.5 µL of 5x DNA Loading Buffer Blue (Bioline®, London, UK). PCR product was visualised on 2% agarose gels (1.6 g Bioline® Agarose in 80 mL 1x sodium borate) stained with ethidium bromide, and gels were imaged using Image Lab Software (Bio-Rad Laboratories Ltd, Watford, UK). A PCR product was deemed positive where there was an amplification band on the gel that was of the expected size (200-300 bp). PCR products were stored at -20 °C until they were pooled according to PCR plate to create sub-libraries for purification with Mag-BIND® RxnPure Plus magnetic beads (Omega Bio-tek Inc, GA, USA), following the double size selection protocol established by Bronner et al. (2009). Ratios of

0.9x and 0.15x magnetic beads to 100 µL of each sub-library were used. Eluted DNA (30 µL) was stored at -20 °C until the second PCR could be performed.

The second PCR bound pre-adapters, MID tags, and Illumina adapters to the purified sub-libraries. Two replicates were performed for each sub-library in 50 µL volumes, consisting of 6 µL of template DNA, 3 µL of each 10 µM primer (Integrated DNA Technologies, Belgium), 25 µL of Q5® High-Fidelity 2x Master Mix (New England Biolabs® Inc., MA, USA) and 13 µL molecular grade water (Fisher Scientific UK Ltd, UK). PCR was performed on an Applied Biosystems® Veriti Thermal Cycler (Life Technologies, CA, USA) with the following thermocycling profile: 95 °C for 3 mins, 8 cycles of 98 °C for 20 s and 72 °C for 1 min, followed by a final elongation step at 72 °C for 5 mins. PCR products were stored at 4 °C until duplicates for each sub-library were pooled, and 2 µL of pooled product was added to 0.5 µL of 5x DNA Loading Buffer Blue (Bioline®, London, UK). PCR products were visualised on 2% agarose gels (1.6 g Bioline® Agarose in 80 mL 1x Sodium borate) stained with ethidium bromide, and gels were imaged using Image Lab Software (Bio-Rad Laboratories Ltd, Watford, UK). Again, PCR products were deemed positive where there was an amplification band on the gel that was of the expected size (300-400 bp). Sub-libraries were stored at 4 °C until purification with Mag-BIND® RxnPure Plus magnetic beads (Omega Bio-tek Inc, GA, USA), following the double size selection protocol established by Bronner et al. (2009). Ratios of 0.7x and 0.15x magnetic beads to 50 µL of each sub-library were used. Eluted DNA (30 µL) was stored at 4 °C until normalisation and final purification.

Sub-libraries were quantified on a Qubit™ 3.0 fluorometer using a Qubit™ dsDNA HS Assay Kit (Invitrogen, UK) and pooled proportional to sample size and concentration. The pooled library was purified using the same ratios, volumes, and protocol as second PCR purification. Based on Qubit™ concentration, the library was diluted to 6 nM for quantification by real-time quantitative PCR (qPCR) using the NEBNext® Library Quant Kit for Illumina® (New England Biolabs® Inc., MA, USA). The library was also checked using an Agilent 2200 TapeStation and High Sensitivity D1000 ScreenTape (Agilent Technologies, CA, USA) to verify secondary product had been removed successfully and a fragment of the expected size (330 bp) remained. The library was frozen and transported in a sterile portable freezer (-20 °C) to Centre for Ecology & Hydrology (CEH), Wallingford, for sequencing. The library was sequenced at 11.5 pM with 10% PhiX Control on an Illumina MiSeq® using 2 x 300 bp V3 chemistry (Illumina Inc., CA, USA).

Raw sequence reads were demultiplexed using a custom Python script then processed using metaBEAT (metaBarcoding and Environmental Analysis Tool) v0.97.11 (https://github.com/HullUni-bioinformatics/metaBEAT). Raw reads were quality trimmed from the read ends (minimum per base phred score Q30) and across sliding windows (window size 5bp; minimum average phred score Q30) using Trimmomatic v0.32 (Bolger, Lohse, & Usadel, 2014). Reads were cropped to a maximum length of 110 bp and reads shorter than 90 bp after quality trimming were discarded. The first 18 bp of remaining reads were also removed to ensure no locus primer remained. Sequence pairs were merged into single high quality reads using FLASH v1.2.11 (Magoč & Salzberg, 2011), provided there was a minimum overlap of 10 bp and no more than 10% mismatch between pairs. Only forward reads were kept for pairs that could not be merged. A final length filter was applied to ensure sequences were reflected the expected fragment size (90-110 bp). Retained sequences were screened for chimeric sequences against a custom reference database for UK vertebrates (Harper et al., 2018) using the uchime algorithm (Edgar, Haas, Clemente, Quince, & Knight, 2011), as implemented in vsearch v1.1 (Rognes, Flouri, Nichols, Quince, & Mahé, 2016). Redundant sequences were removed by clustering at 100% identity ('--cluster_fast' option) in vsearch v1.1 (Rognes et al., 2016). Clusters were considered sequencing error and omitted from further processing if they were represented by less than three sequences. Non-redundant sets of query sequences were then compared against the UK vertebrate reference database (Harper et al., 2018) using BLAST (Zhang, Schwartz, Wagner, & Miller, 2000). Putative taxonomic identity was assigned using a lowest common ancestor (LCA) approach based on the top 10% BLAST matches for any query that matched a reference sequence across more than 80% of its length at minimum identity of 98%. Unassigned sequences were subjected to a separate BLAST search against the complete NCBI nucleotide (nt) database at 98% identity to determine the source via LCA as described above. The bioinformatic analysis has been deposited in the GitHub repository for reproducibility (permanently archived at: https://doi.org/10.5281/zenodo.2561415).

#### 1.2 Data analysis

##### *1.2.1 Dataset refinement*

Assignments from different databases were merged, and spurious assignments (i.e. non-UK species, invertebrates and bacteria) removed from the dataset. The family Cichlidae was reassigned to *Maylandia zebra*. The genera *Bison*, *Bos*, *Buteo*, *Castor*, *Meleagris*, *Pelophylax*, *Sprattus*, *Strix*, and *Triturus* were reassigned to European bison (*Bison bonasus*), cow (*Bos taurus*), common buzzard (*Buteo buteo*), Eurasian beaver (*Castor fiber*), marsh frog (*Pelophylax ridibundus*), turkey (*Meleagris gallopavo*), European sprat (*Sprattus sprattus*), tawny owl (*Strix aluco*), and great crested newt (*Triturus cristatus*) respectively based on local knowledge of sampling sites and UK distribution maps (National Biodiversity Network Atlas, 2019). The species *Sus scrofa* and *Canis lupus* were reassigned to pig (*Sus scrofa domesticus*) and (*Canis lupus familiaris*) given the restricted distribution of wild boar (*S. scrofa*) and absence of grey wolf (*C. lupus*) in the UK.

Misassignments included the cichlids *Haplochromis burtoni*, *Oreochromis niloticus*, and *Pundamilia nyererei* which were reassigned to *M. zebra*, and Iberian lynx (*Lynx pardinus*) which was reassigned to Eurasian lynx (*Lynx lynx*). Other potential misassignments were green-winged teal (*Anas carolinensis*), yellow-browed bunting (*Emberiza chrysophrys*), and Iceland gull (*Larus glaucoides*). These are rare migrants that have been infrequently recorded in the UK (British Trust for Ornithology, 2019) but may have been assigned due to potential for hybridisation within the genus *Anas* and missing reference sequences for several common species within the genera *Emberiza* and *Larus*. These species were reassigned to the genera *Anas*, *Emberiza*, and *Larus*. Reads from corrected assignments were then merged with unaltered assignments.

Of 89 process controls included throughout the metabarcoding workflow, 39 produced no reads. Reads generated for 50 of 89 process controls ranged from 3 to 4930, and strength of each contaminant varied (mean = 62.4%, range = 0.3 - 100.0% of the total reads per process control). Environmental contamination was observed in the field blanks (European badger [*Meles meles*], beaver, lynx, European pine marten [*Martes martes*], red squirrel [*Sciurus vulgaris*], and European water vole [*Arvicola amphibius*]) as well as environmental and/or laboratory contamination in the filtration and extraction blanks (human [*Homo sapiens*] and cichlid). PCR negative controls were also contaminated with human, cichlid, beaver, and pine marten as well as non-focal species (Fig. S3). Consequently, we evaluated different sequence thresholds to minimise the risk of false positives in our dataset. These included the maximum sequence frequency of cichlid DNA in eDNA samples (0.308%), maximum sequence frequency of any DNA except cichlid in PCR positive controls (0.064%), and taxon-specific thresholds (maximum sequence frequency of each taxon in PCR positive controls). The different thresholds were applied to the eDNA samples and the results from each compared (Fig. S4). The taxon-specific thresholds (Table S4) retained the most biological information, thus these were selected for downstream analysis. Consequently, taxa were only classed as present at sites if their sequence frequency exceeded taxon-specific thresholds.

Contaminants remaining in eDNA samples after threshold application included Gentoo penguin (*Pygoscelis papua*) and reindeer (*Rangifer tarandus*) which were likely sourced from the environment, cichlid sourced from the laboratory, and human which may have originated from the environment or the laboratory. Gentoo penguin was only detected in water from the beaver quarantine enclosure at RZSS Edinburgh Zoo, and reindeer was only detected in water sampled from a red squirrel enclosure at Wildwood Trust. Human DNA was detected in the majority of eDNA samples, and cichlid DNA was also present at low frequency in some samples. These contaminants and assignments higher than species level were removed from the dataset, excluding the genera *Anas*, *Emberiza*, and *Larus*. Therefore, all taxonomic assignments in the final dataset were predominantly of species resolution and considered real detections. European bison, which is present in the red deer (*Cervus elaphus*) enclosure at RZSS Highland Wildlife Park, was detected in two samples taken from this enclosure. However, these detections were excluded from downstream analyses as bison was not one of our focal species. Samples belonging to focal species in Experiment 1 were contaminated with DNA of other focal species to different extents (mean = 6.7%, range = 0.0 - 100.0% of the total refined reads per sample). Therefore, any proportional reads for incorrect focal species in each enclosure were set to 0 for the purposes of downstream analysis.

*1.2.2 Effect of water volume and filtration method*

We tested the hypothesis that volume of water filtered or number of filters used may affect read counts. A hierarchical binomial Generalized Linear Mixed Model (GLMM) with the logit link function from the development version of the R package glmmTMB (Brooks et al., 2017), including volume and number of filters as fixed effects and species nested within wildlife park as a random effect, was used. Validation checks were performed to ensure all model assumptions were met where possible and absence of overdispersion (Zuur, Ieno, Walker, Saveliev, & Smith, 2009). Model fit was assessed visually and with the Hosmer and Lemeshow Goodness of Fit Test (Hosmer & Lemeshow, 2000) using the R package ResourceSelection v0.3-2 (Lele, Keim, & Solymos, 2016). Predictions for each model were obtained using the *predict* function and upper and lower 95% CIs were calculated from the standard error of the predictions. Figures were produced using the R package ggplot2 v3.3.0 (Wickham, 2016).

### 2. Results and discussion

#### 2.1 Effect of water volume and filtration method

Neither volume of water filtered ($\chi^2_1$ = 2.141 ± 0.143) or number of filters used ($\chi^2_1$ = 0.108 ± 0.742) had a significant effect on the proportional read counts based on the hierarchical model (θ = 0.221, $\chi^2_{76}$ = 16.798, $P$ = 1.000, pseudo-$R^2$ = 20.57%). Proportional read counts somewhat decreased (-0.003 ± 0.002, $Z$ = -1.389, $P$ = 0.165) as water volume filtered increased (Fig. S1a), and marginally increased (0.232 ± 0.703, $Z$ = 0.331, $P$ = 0.741) where two filters were used for water filtration (Fig. S1b).

#### 2.2 Technical pitfalls of eDNA metabarcoding for mammal monitoring

Our process controls identified low-level contamination from domestic and wild species at all stages of metabarcoding, but contaminants primarily occurred in field and PCR negative controls. We also identified cichlid sequences in field, filtration, and extraction controls, despite this DNA not being handled before PCR. Contaminants found in process controls may stem from PCR contamination (Kelly, Port, Yamahara, & Crowder, 2014) or sequencing error (Hänfling et al., 2016). Indeed, PCR negative controls corresponding to Tophill Low Nature Reserve samples were contaminated with great crested newt (*Triturus cristatus*), a highly abundant pond species at the reserve. This suggests that concentrated samples can contaminate negative controls during metabarcoding. Negative controls at each stage of metabarcoding can identify contaminant introduction (Klymus, Richter, Thompson, & Hinck, 2017; Ushio et al., 2017), but contaminants in these controls can amplify exponentially with no competition affecting inferences (Harper et al., 2018). Innovative approaches are needed to mitigate metabarcoding contamination, such as false positive estimation using occupancy models (Ficetola et al., 2015) or sequencing thresholds, i.e. the number of sequence reads required for a sample to be species-positive (Evans et al., 2017; Hänfling et al., 2016).

### 3. Tables

**Table A1.** Behavioural observation data for species housed at wildlife parks, including date, time, weather conditions, behaviour, and frequency and duration of behaviour.

| Location | Date | Start time | End time | Weather | Air temperature (°C) | Species | Enclosure | Behaviour | Frequency | Duration (hrs) |
|---|---|---|---|---|---|---|---|---|---|---|
| Wildwood Trust | 18/09/2017 | 10:29 | 11:29 | Cloudy | 13 | Red deer (*Cervus elaphus*) | 1 | Drinking | 9 | 9.37 |
| | | | | | | | | Feeding | 3 | 22.37 |
| | | | | | | | | Defecating | 1 | 0.5 |
| | | | | | | | | Sniffing | 1 | 0.53 |
| | | | | | | | | Standing | 6 | 30 |
| | | | | | | | | Walking | 12 | 22.27 |
| Wildwood Trust | 19/09/2017 | 09:50 | 10:50 | Sunny | 17 | Red deer (*Cervus elaphus*) | 1 | Drinking | 2 | 4.47 |
| | | | | | | | | Feeding | 2 | 40.68 |
| | | | | | | | | Standing | 1 | 34.8 |
| | | | | | | | | Walking | 1 | 5.75 |
| | | | | | | | | Resting | 1 | 6.7 |
| Wildwood Trust | 18/09/2017 | 11:38 | 12:38 | Partial sun | 16 | Eurasian lynx (*Lynx lynx*) | 1 | Scratching | 1 | 0.08 |
| | | | | | | | | Urinating | 2 | 3.42 |
| | | | | | | | | Standing | 1 | 0.38 |
| | | | | | | | | Walking | 9 | 18.12 |
| | | | | | | | | Walking | 2 | 9.17 |
| | | | | | | | | Running | 1 | 0.38 |
| | | | | | | | | Vocalising | 4 | 4.58 |
| | | | | | | | | Resting | 4 | 29.6 |
| | | | | | | | | Resting | 1 | 1.32 |
| | | | | | | | | Grooming | 4 | 5.03 |
| | | | | | | | | Grooming | 1 | 0.08 |
| | | | | | | | | Not visible | 1 | 1.88 |
| | | | | | | | | Not visible | 2 | 48.25 |

| Location | Date | Start time | End time | Weather | Air temperature (°C) | Species | Enclosure | Behaviour | Frequency | Duration (hrs) |
|---|---|---|---|---|---|---|---|---|---|---|
| Wildwood Trust | 19/09/2017 | 09:30 | 10:30 | Cloudy | 12 | Eurasian lynx (*Lynx lynx*) | 1 | Drinking | 3 | 1.13 |
| | | | | | | | | Urinating | 1 | 0.01 |
| | | | | | | | | Not visible | 1 | 10.52 |
| | | | | | | | | Resting | 1 | 2.42 |
| | | | | | | | | Walking | 3 | 46.8 |
| | | | | | | | | Walking | 3 | 27.92 |
| | | | | | | | | Resting | 1 | 0.77 |
| | | | | | | | | Grooming | 1 | 0.45 |
| | | | | | | | | Other | 1 | 0.97 |
| | | | | | | | | Not visible | 2 | 30.95 |
| Wildwood Trust | 18/09/2017 | 11:36 | 12:32 | Partial sun | 16 | European pine marten (*Martes martes*) | 1 | Immersed | 10 | 0.35 |
| | | | | | | | | Drinking | 2 | 0.03 |
| | | | | | | | | Urinating | 1 | 0.03 |
| | | | | | | | | Other | 19 | 0.67 |
| Wildwood Trust | 21/09/2017 | 09:26 | 09:56 | Sunny | 17 | European pine marten (*Martes martes*) | 1 | Drinking | 1 | 0.12 |
| | | | | | | | | Sniffing | 7 | 6.83 |
| | | | | | | | | Standing | 1 | 0.82 |
| | | | | | | | | Walking | 10 | 8.45 |
| | | | | | | | | Resting | 10 | 12.18 |
| | | | | | | | | Playing | 9 | 5.22 |
| | | | | | | | | Not visible | 1 | 0.98 |
| Wildwood Trust | 21/09/2017 | 09:56 | 10:26 | Sunny | 17 | European pine marten (*Martes martes*) | 2 | Immersed | 3 | 0.2 |
| | | | | | | | | Urinating | 9 | 0.27 |
| | | | | | | | | Standing | 1 | 0.23 |
| | | | | | | | | Running | 4 | 29.57 |
| | | | | | | | | Resting | 1 | 0.2 |

| Location | Date | Start time | End time | Weather | Air temperature (°C) | Species | Enclosure | Behaviour | Frequency | Duration (hrs) |
|---|---|---|---|---|---|---|---|---|---|---|
| Wildwood Trust | 18/09/2017 | 13:25 | 13:55 | Partial sun | 17 | Red squirrel (*Sciurus vulgaris*) | 1 | Feeding | 5 | 3.77 |
| | | | | | | | | Walking | 1 | 1.15 |
| | | | | | | | | Running | 4 | 3.68 |
| | | | | | | | | Resting | 2 | 0.8 |
| | | | | | | | | Drinking | 2 | 0.93 |
| | | | | | | | | Running | 4 | 23.83 |
| | | | | | | | | Resting | 2 | 5.27 |
| Wildwood Trust | 18/09/2017 | 14:00 | 14:30 | Partial sun | 17 | Red squirrel (*Sciurus vulgaris*) | 2 | Drinking | 1 | 1.4 |
| | | | | | | | | Feeding | 4 | 15.5 |
| | | | | | | | | Running | 3 | 13.1 |
| | | | | | | | | Drinking | 2 | 0.57 |
| | | | | | | | | Feeding | 3 | 1.07 |
| | | | | | | | | Running | 6 | 27.37 |
| Wildwood Trust | 18/09/2017 | 13:45 | 14:45 | Sunny | 16 | European otter (*Lutra lutra*) | 1 | Swimming | 12 | 14.25 |
| | | | | | | | | Standing | 1 | 0.35 |
| | | | | | | | | Playing | 2 | 5.28 |
| Wildwood Trust | 19/09/2017 | 09:21 | 10:21 | Cloudy | 12 | European otter (*Lutra lutra*) | 1 | Swimming | 8 | 16.4 |
| | | | | | | | | Feeding | 3 | 5.7 |
| | | | | | | | | Sniffing | 5 | 8.33 |
| | | | | | | | | Not visible | 4 | 35.43 |
| Wildwood Trust | 19/09/2017 | 19:54 | 05:04 | Cloudy | 9 | European badger (*Meles meles*) | 1 | Drinking | 14 | 0.95 |
| | | | | | | | | Sniffing | 36 | 8.43 |
| | | | | | | | | Walking | 36 | 8.43 |
| [illegible] | 19/09/2017 | 20:47 | 06:58 | Clear | 9 | Eurasian beaver | 1 | Swimming | 40 | 13.3 |

| Location | Date | Start time | End time | Weather | Air temperature (°C) | Species | Enclosure | Behaviour | Frequency | Duration (hrs) |
|---|---|---|---|---|---|---|---|---|---|---|
| | | | | | | (*Castor fiber*) | | Sniffing | 1 | 0.33 |
| | | | | | | | | Standing | 1 | 0.15 |
| | | | | | | | | Walking | 12 | 1.52 |
| Highland Wildlife Park | 09/10/17 | 15:09 | 16:09 | Partial sun | 13 | Eurasian lynx (*Lynx lynx*) | 1 | Drinking | 1 | 46 |
| | | | | | | | | Feeding | 2 | 8.77 |
| | | | | | | | | Walking | 6 | 19.82 |
| | | | | | | | | Resting | 2 | 13.88 |
| | | | | | | | | Grooming | 1 | 2 |
| | | | | | | | | Other | 2 | 0.92 |
| Highland Wildlife Park | NA | NA | NA | NA | NA | Eurasian beaver (*Castor fiber*) | 1 | Swimming | 6 | 1.07 |
| | | | | | | | | Standing | 2 | 0.5 |
| | | | | | | | | Sniffing | 1 | 0.08 |
| | | | | | | | | Feeding | 1 | 0.27 |
| | | | | | | | | Other | 1 | 0.27 |
| Highland Wildlife Park | 09/10/17 | 09:55 | 12:25 | Cloudy | 12 | Red deer (*Cervus elaphus*) | 1 | Drinking | 1 | 0.02 |
| | | | | | | | | Feeding | 4 | 32.32 |
| | | | | | | | | Walking | 7 | 16.62 |
| | | | | | | | | Resting | 1 | 23 |
| | | | | | | | | Other | 1 | 0.08 |
| | | | | | | | | Not visible | 5 | 42.78 |

**Table A2.** Summary of directed, random, or other samples collected for each species at wildlife parks.

| Site | Species | Enclosure | Sample type | Number of samples | Volume filtered (ml) |
|---|---|---|---|---|---|
| Wildwood Trust | European otter (*Lutra lutra*) | 1 | Targeted | 5 | 500 |
| | | | Passive | 6 | 500 |
| | European water vole (*Arvicola amphibius*) | 1 | Other | 1 | 250 |
| | | 2 | Other | 1 | 250 |
| | Eurasian beaver (*Castor fiber*) | 1 | Targeted | 4 | 150 |
| | | | Passive | 5 | 150-200 |
| | | 2 | Passive | 1 | 150 |
| | European hedgehog (*Erinaceus europaeus*) | 1 | Other | 1 | 250 |
| | | 2 | Other | 1 | 250 |
| | European badger (*Meles meles*) | 1 | Targeted | 3 | 500 |
| | | | Passive | 3 | 500 |
| | | | Other | 1 | 500 |
| | Red deer (*Cervus elaphus*) | 1 | Targeted | 6 | 10-75 |
| | | | Passive | 4 | 25-150 |
| | Eurasian lynx (*Lynx lynx*) | 1 | Targeted | 1 | 500 |
| | | | Passive | 2 | 500 |
| | | | Other | 1 | 500 |
| | Red squirrel (*Sciurus vulgaris*) | 1 | Other | 1 | 250 |
| | | 2 | Other | 1 | 250 |
| | | 3 | Other | 1 | 250 |
| | | 4 | Other | 1 | 250 |
| | European pine marten | 1 | Targeted | 3 | 500 |

| **Site** | **Species** | **Enclosure** | **Sample type** | **Number of samples** | **Volume filtered (ml)** |
|---|---|---|---|---|---|
| | (*Martes martes*) | | Passive | 2 | 500 |
| | | | Other | 1 | 500 |
| | | 2 | Targeted | 1 | 500 |
| | | | Passive | 1 | 500 |
| Highland Wildlife Park | Red squirrel (*Sciurus vulgaris*) | NA | Other | 1 | 500 |
| | Eurasian lynx (*Lynx lynx*) | 1 | Targeted | 1 | 500 |
| | | | Passive | 2 | 500 |
| | | | Other | 1 | 500 |
| | Eurasian beaver (*Castor fiber*) | 1 | Targeted | 3 | 500 |
| | | | Passive | 3 | 500 |
| | | | Other | 1 | 100 |
| | Red deer (*Cervus elaphus*) | 1 | Targeted | 5 | 50-200 |
| | | | Passive | 5 | 125-500 |

**Table A3.** Summary of samples collected from natural ponds at locations where target species were confirmed as present.

| Site | Date | Pond | Sample | Volume filtered (L) |
|---|---|---|---|---|
| Thorne Moors | 17/04/2018 | 1 | 1 | 1.5 |
| | | | 2 | 1 |
| | | | 3 | 0.65 |
| | | | 4 | 0.8 |
| | | | 5 | 0.85 |
| | | | 6 | 0.8 |
| | | | 7 | 1 |
| | | | 8 | 1.5 |
| | | | 9 | 0.15 |
| | | | 10 | 0.1 |
| | | 2 | 1 | 0.175 |
| | | | 2 | 0.4 |
| | | | 3 | 0.4 |
| | | | 4 | 0.5 |
| | | | 5 | 0.75 |
| | | | 6 | 0.75 |
| | | | 7 | 0.3 |
| | | | 8 | 0.3 |
| | | | 9 | 0.4 |
| | | | 10 | 0.6 |
| Bamff Estate | 20/04/2018 | 1 | 1 | 0.75 |
| | | | 2 | 0.45 |

| Site | Date | Pond | Sample | Volume filtered (L) |
|---|---|---|---|---|
| | | | 3 | 0.55 |
| | | | 4 | 1 |
| | | | 5 | 2 |
| | | | 6 | 0.9 |
| | | | 7 | 1 |
| | | | 8 | 0.95 |
| | | | 9 | 1.05 |
| | | | 10 | 0.6 |
| | | 2 | 1 | 0.95 |
| | | | 2 | 0.95 |
| | | | 3 | 0.85 |
| | | | 4 | 1 |
| | | | 5 | 0.95 |
| | | | 6 | 0.95 |
| | | | 7 | 1.1 |
| | | | 8 | 0.95 |
| | | | 9 | 0.95 |
| | | | 10 | 0.95 |
| Tophill Low Nature Reserve | 23/04/2018 | 1 | 1 | 0.6 |
| | | | 2 | 0.625 |
| | | | 3 | 0.625 |
| | | | 4 | 0.675 |
| | | | 5 | 1.1 |
| | | | 6 | 0.85 |
| | | | 7 | 1 |

| Site | Date | Pond | Sample | Volume filtered (L) |
|---|---|---|---|---|
| | | | 8 | 0.625 |
| | | | 9 | 0.6 |
| | | | 10 | 0.65 |
| | | 2 | 1 | 1 |
| | | | 2 | 1 |
| | | | 3 | 1 |
| | | | 4 | 1 |
| | | | 5 | 0.9 |
| | | | 6 | 0.9 |
| | | | 7 | 0.95 |
| | | | 8 | 0.95 |
| | | | 9 | 0.95 |
| | | | 10 | 0.85 |
| | 24/04/2018 | 1 | 1 | 0.625 |
| | | | 2 | 0.8 |
| | | | 3 | 0.7 |
| | | | 4 | 0.65 |
| | | | 5 | 0.9 |
| | | | 6 | 0.8 |
| | | | 7 | 0.75 |
| | | | 8 | 0.65 |
| | | | 9 | 0.65 |
| | | | 10 | 0.625 |
| | | 2 | 1 | 1 |
| | | | 2 | 0.9 |

| Site | Date | Pond | Sample | Volume filtered (L) |
|---|---|---|---|---|
| | | | 3 | 0.9 |
| | | | 4 | 1 |
| | | | 5 | 0.875 |
| | | | 6 | 0.825 |
| | | | 7 | 0.85 |
| | | | 8 | 1.1 |
| | | | 9 | 1.2 |
| | | | 10 | 1.1 |
| | 25/04/2018 | 1 | 1 | 0.65 |
| | | | 2 | 0.75 |
| | | | 3 | 0.825 |
| | | | 4 | 0.55 |
| | | | 5 | 1 |
| | | | 6 | 0.8 |
| | | | 7 | 0.8 |
| | | | 8 | 0.65 |
| | | | 9 | 0.725 |
| | | | 10 | 0.55 |
| | | 2 | 1 | 0.9 |
| | | | 2 | 0.9 |
| | | | 3 | 1 |
| | | | 4 | 0.925 |
| | | | 5 | 0.85 |
| | | | 6 | 0.775 |
| | | | 7 | 0.875 |

| Site | Date | Pond | Sample | Volume filtered (L) |
|---|---|---|---|---|
| | | | 8 | 1.1 |
| | | | 9 | 1.1 |
| | | | 10 | 1 |
| | 26/04/2018 | 1 | 1 | 0.55 |
| | | | 2 | 0.775 |
| | | | 3 | 0.7 |
| | | | 4 | 0.7 |
| | | | 5 | 0.75 |
| | | | 6 | 0.85 |
| | | | 7 | 0.8 |
| | | | 8 | 0.65 |
| | | | 9 | 0.65 |
| | | | 10 | 0.725 |
| | | 2 | 1 | 0.9 |
| | | | 2 | 0.9 |
| | | | 3 | 1 |
| | | | 4 | 0.9 |
| | | | 5 | 0.9 |
| | | | 6 | 0.775 |
| | | | 7 | 0.65 |
| | | | 8 | 0.75 |
| | | | 9 | 0.85 |
| | | | 10 | 0.7 |
| | 27/04/2018 | 1 | 1 | 0.7 |
| | | | 2 | 0.65 |

| Site | Date | Pond | Sample | Volume filtered (L) |
|---|---|---|---|---|
| | | | 3 | 0.75 |
| | | | 4 | 0.65 |
| | | | 5 | 0.85 |
| | | | 6 | 0.8 |
| | | | 7 | 0.85 |
| | | | 8 | 0.7 |
| | | | 9 | 0.7 |
| | | | 10 | 0.75 |
| | | 2 | 1 | 0.85 |
| | | | 2 | 0.8 |
| | | | 3 | 0.95 |
| | | | 4 | 0.75 |
| | | | 5 | 0.8 |
| | | | 6 | 0.85 |
| | | | 7 | 0.8 |
| | | | 8 | 0.85 |
| | | | 9 | 1.1 |
| | | | 10 | 0.9 |

**Table A4.** List of taxa detected in PCR positive controls by eDNA metabarcoding and corresponding taxon-specific false positive sequence threshold applied.

| **Taxonomic assignment** | **Common name** | **Threshold** |
|---|---|---|
| *Anas* spp. | Dabbling ducks | 0.00067132 |
| Anatidae | Ducks, geese, swans | 0.000100995 |
| *Arvicola amphibius* | European water vole | 0.000342575 |
| Aves | Birds | 0.000054 |
| *Castor fiber* | European beaver | 0.003023912 |
| *Columba* | Pigeons | 0.0000877 |
| Corvidae | Corvids | 0.000081 |
| Gasterosteidae | Sticklebacks | 0.001862034 |
| *Homo sapiens* | Human | 0.000873784 |
| *Lynx lynx* | Eurasian lynx | 0.0000585 |
| *Martes martes* | European pine marten | 0.000906857 |
| *Mus musculus* | Mouse | 0.000107263 |
| Passeriformes | Songbirds | 0.0000202 |
| *Pelophylax ridibundus* | Marsh frog | 0.0000743 |
| Phasianidae | Gamebirds | 0.000107263 |
| *Phoxinus phoxinus* | Common minnow | 0.000092 |
| *Pungitius pungitius* | Ninespine stickleback | 0.026399055 |
| *Rana temporaria* | Common frog | 0.064393287 |
| *Sus scrofa domesticus* | Pig | 0.000148423 |
| *Triturus cristatus* | Great crested newt | 0.001758274 |
| unassigned | NA | 0.009074043 |

**Table A5.** Summary of species detected using eDNA metabarcoding across all samples collected in this study.

| Common name | Binomial name | Number of samples (*N* = 220) |
|---|---|---|
| Red-legged partridge | *Alectoris rufa* | 2 |
| Dabbling ducks | *Anas* spp. | 38 |
| European eel | *Anguilla anguilla* | 6 |
| Grey heron | *Ardea cinerea* | 14 |
| European water vole | *Arvicola amphibius* | 12 |
| European bison | *Bison bonasus* | 2 |
| Cow | *Bos taurus* | 44 |
| Common toad | *Bufo bufo* | 22 |
| Common buzzard | *Buteo buteo* | 3 |
| Dog | *Canis lupus familiaris* | 4 |
| Roe deer | *Capreolus capreolus* | 4 |
| European beaver | *Castor fiber* | 50 |
| Red deer | *Cervus elaphus* | 36 |
| Atlantic herring | *Clupea harengus* | 24 |
| Rock dove | *Columba livia* | 6 |
| Stock dove | *Columba oenas* | 6 |
| Common quail | *Coturnix coturnix* | 8 |
| Grass carp | *Ctenopharyngodon idella* | 1 |
| Buntings | *Emberiza* spp. | 1 |
| Horse | *Equus caballus* | 28 |
| European hedgehog | *Erinaceus europaeus* | 1 |
| European robin | *Erithacus rubecula* | 8 |
| Common moorhen | *Gallinula chloropus* | 10 |

| **Common name** | **Binomial name** | **Number of samples (*N* = 220)** |
|---|---|---|
| Eurasian jay | *Garrulus glandarius* | 1 |
| Three-spined stickleback | *Gasterosteus aculeatus* | 18 |
| White-headed gulls | *Larus* spp. | 3 |
| Palmate newt | *Lissotriton helveticus* | 4 |
| Smooth newt | *Lissotriton vulgaris* | 80 |
| European otter | *Lutra lutra* | 16 |
| Eurasian lynx | *Lynx lynx* | 22 |
| European pine marten | *Martes martes* | 16 |
| Turkey | *Meleagris gallopavo* | 3 |
| European badger | *Meles meles* | 25 |
| Mouse | *Mus musculus* | 11 |
| Bank vole | *Myodes glareolus* | 2 |
| Eurasian water shrew | *Neomys fodiens* | 7 |
| Red-crested pochard | *Netta rufina* | 18 |
| European rabbit | *Oryctolagus cuniculus* | 37 |
| European smelt | *Osmerus eperlanus* | 2 |
| Sheep | *Ovis aries* | 9 |
| Great tit | *Parus major* | 10 |
| Marsh frog | *Pelophylax ridibundus* | 11 |
| Common pheasant | *Phasianus colchicus* | 19 |
| Common minnow | *Phoxinus phoxinus* | 10 |
| Eurasian magpie | *Pica pica* | 7 |
| Common pipistrelle | *Pipistrellus pipistrellus* | 1 |
| Ninespine stickleback | *Pungitius pungitius* | 7 |
| Common frog | *Rana temporaria* | 20 |

| **Common name** | **Binomial name** | **Number of samples (*N* = 220)** |
|---|---|---|
| Brown rat | *Rattus norvegicus* | 10 |
| Brown trout | *Salmo trutta* | 12 |
| Grey squirrel | *Sciurus carolinensis* | 9 |
| Red squirrel | *Sciurus vulgaris* | 13 |
| Common shrew | *Sorex araneus* | 2 |
| European sprat | *Sprattus sprattus* | 5 |
| Tawny owl | *Strix aluco* | 1 |
| Common starling | *Sturnus vulgaris* | 2 |
| Pig | *Sus scrofa domesticus* | 45 |
| European mole | *Talpa europaea* | 1 |
| Great crested newt | *Triturus cristatus* | 100 |
| Eurasian wren | *Troglodytes troglodytes* | 1 |
| Song thrush | *Turdus philomelos* | 10 |
| Northern lapwing | *Vanellus vanellus* | 1 |

**Table A6.** Summary of detection rates for species which were detected by at least one survey method performed at six ponds across three sites in this study.

| Species | Lifestyle | Field signs | Camera trapping | eDNA metabarcoding |
|---|---|---|---|---|
| Common pipistrelle (*Pipistrellus pipistrellus*) | Arboreal | 0/6 (0%) | 0/6 (0%) | 1/6 (16.67%) |
| Grey squirrel (*Sciurus carolinensis)* | Arboreal | 0/6 (0%) | 0/6 (0%) | 2/6 (33.33%) |
| Cow (*Bos taurus*) | Ground-dwelling | 0/6 (0%) | 0/6 (0%) | 3/6 (50%) |
| Sheep (*Ovis aries*) | Ground-dwelling | 0/6 (0%) | 0/6 (0%) | 4/6 (66.67%) |
| Pig (*Sus scrofa domesticus*) | Ground-dwelling | 0/6 (0%) | 0/6 (0%) | 3/6 (50%) |
| Dog (*Canis lupus familiaris*) | Ground-dwelling | 0/6 (0%) | 0/6 (0%) | 2/6 (33.33%) |
| Roe deer (*Capreolus capreolus*) | Ground-dwelling | 4/6 (66.67%) | 2/6 (33.33%) | 3/6 (50%) |
| Red deer (*Cervus elaphus*) | Ground-dwelling | 2/6 (33.33%) | 1/6 (16.67%) | 1/6 (16.67%) |
| Red fox (*Vulpes vulpes*) | Ground-dwelling | 1/6 (16.67%) | 3/6 (50%) | 0/6 (0%) |
| Badger (*Meles meles*) | Ground-dwelling | 1/6 (16.67%) | 0/6 (0%) | 0/6 (0%) |
| Bank vole (*Myodes glareolus*) | Ground-dwelling | 0/6 (0%) | 0/6 (0%) | 1/6 (16.67%) |
| Common shrew (*Sorex araneus*) | Ground-dwelling | 0/6 (0%) | 0/6 (0%) | 1/6 (16.67%) |
| Rabbit (*Oryctolagus cuniculus*) | Ground-dwelling | 0/6 (0%) | 0/6 (0%) | 1/6 (16.67%) |
| Water vole (*Arvicola amphibius*) | Semi-aquatic | 0/2 (0%) | 0/2 (0%) | 1/6 (16.67%) |

| | | | | |
|---|---|---|---|---|
| Water shrew (*Neomys fodiens*) | Semi-aquatic | 0/6 (0%) | 0/6 (0%) | 2/6 (33.33%) |
| Brown rat (*Rattus norvegicus*) | Semi-aquatic | 0/6 (0%) | 0/6 (0%) | 1/6 (16.67%) |
| Beaver (*Castor fiber*) | Semi-aquatic | 2/6 (33.33%) | 2/6 (33.33%) | 2/6 (33.33%) |

#### 4. Figures

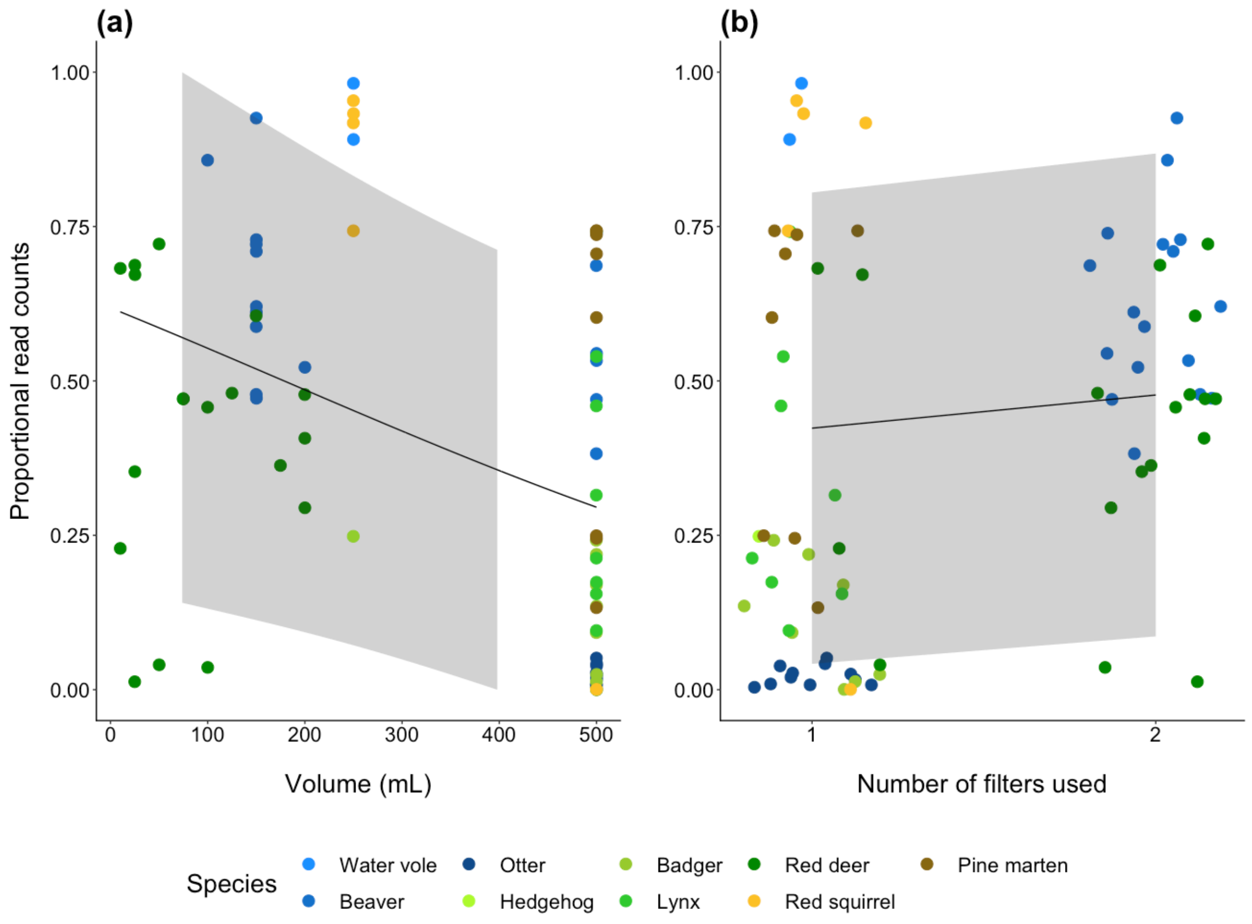


**Figure A1.** Relationship between the fixed effects (volume and number of filters) and response variable (proportional read counts) as predicted by the binomial GLMM. The 95% CIs, as calculated using the predicted proportional read counts and standard error for these predictions, are given for each relationship. The observed data (points) are displayed against the predicted relationships (lines). Proportional read count marginally decreased as volume of water filtered increased **(a)**, but increased as number of filters used increased **(b)**.

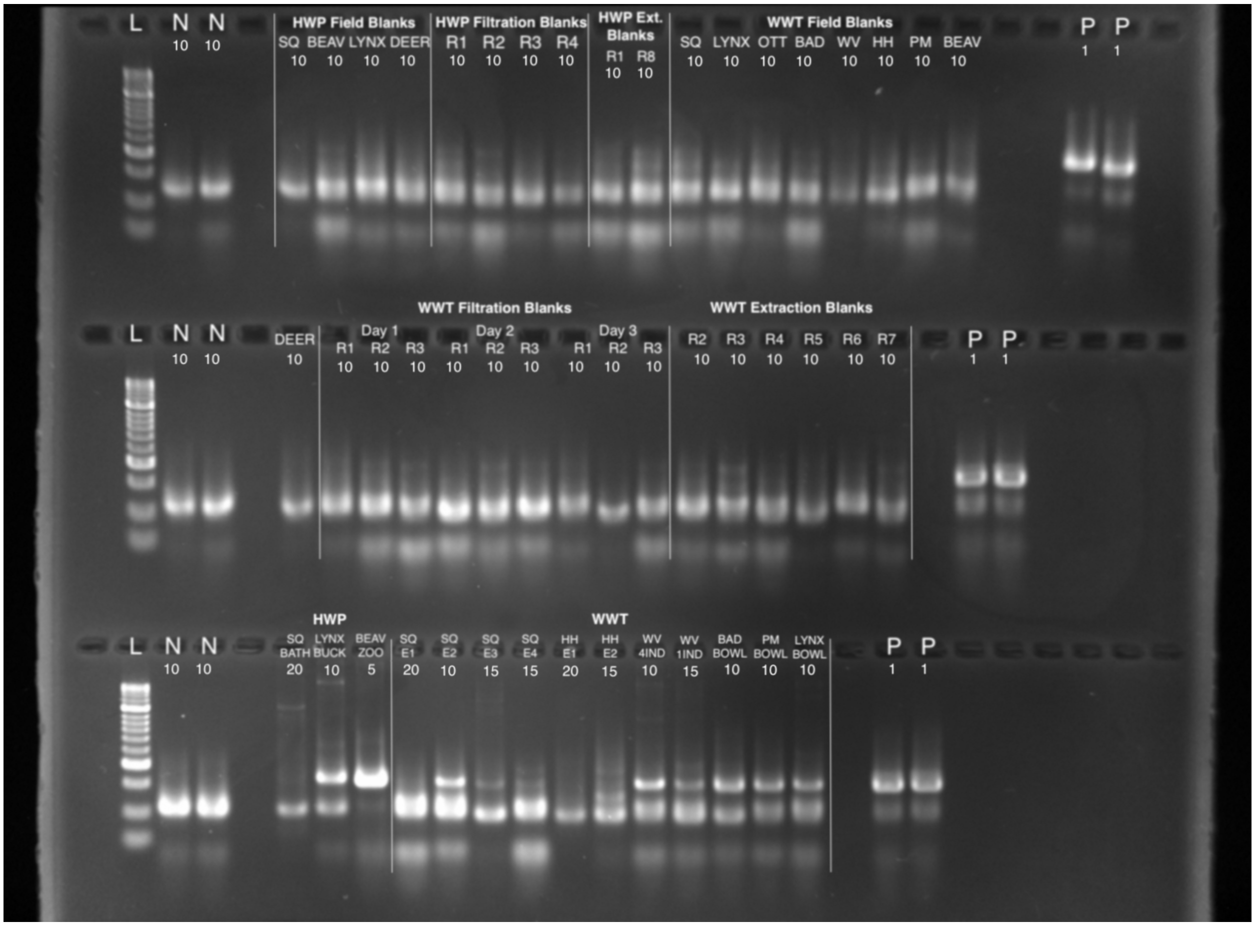


**Figure A2.** Example gel image of pooled first PCR products which were run on 2% agarose gels with Hyperladder™ 50bp (Bioline®, London, UK) molecular weight marker (L). PCR products were assigned an amplification score based on band strength (0 = no band, 1 = faint band, 2 = bright band, 3 = very bright band). These scores were used to determine how much product should be pooled to create each sub-library (0 = 20 µL, 1 = 15 µL, 2 = 10 µL, 3 = 5 µL). All blanks and PCR negative controls were pooled in consistent volumes (10 µL). Only 1 µL of each PCR positive control was pooled.

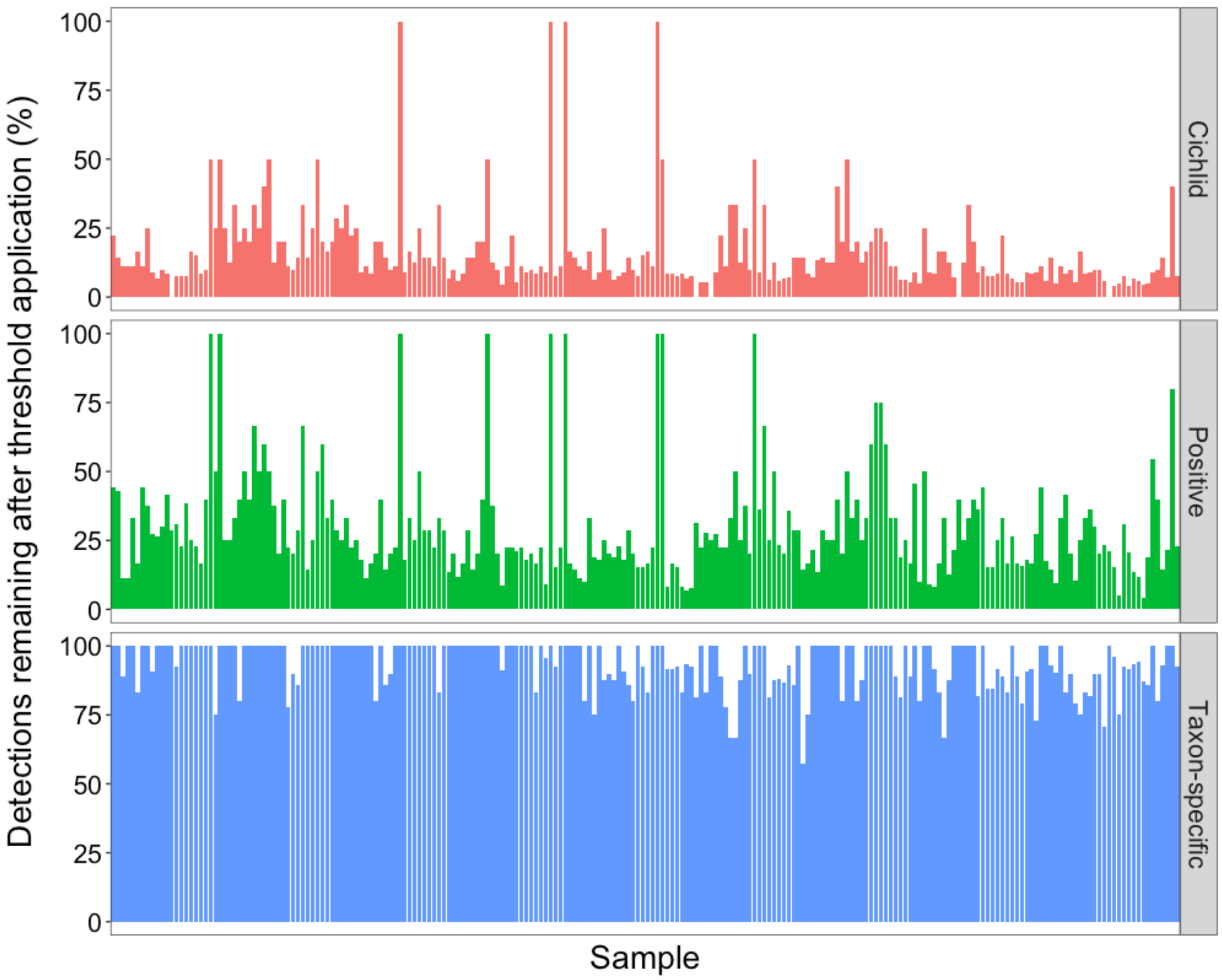


**Figure A3.** Barplot showing the impact of different false positive sequence thresholds on the proportion of taxa detected in each sample. The taxon-specific thresholds retained the most biological information, thus these were applied to the eDNA metabarcoding data for downstream analyses.

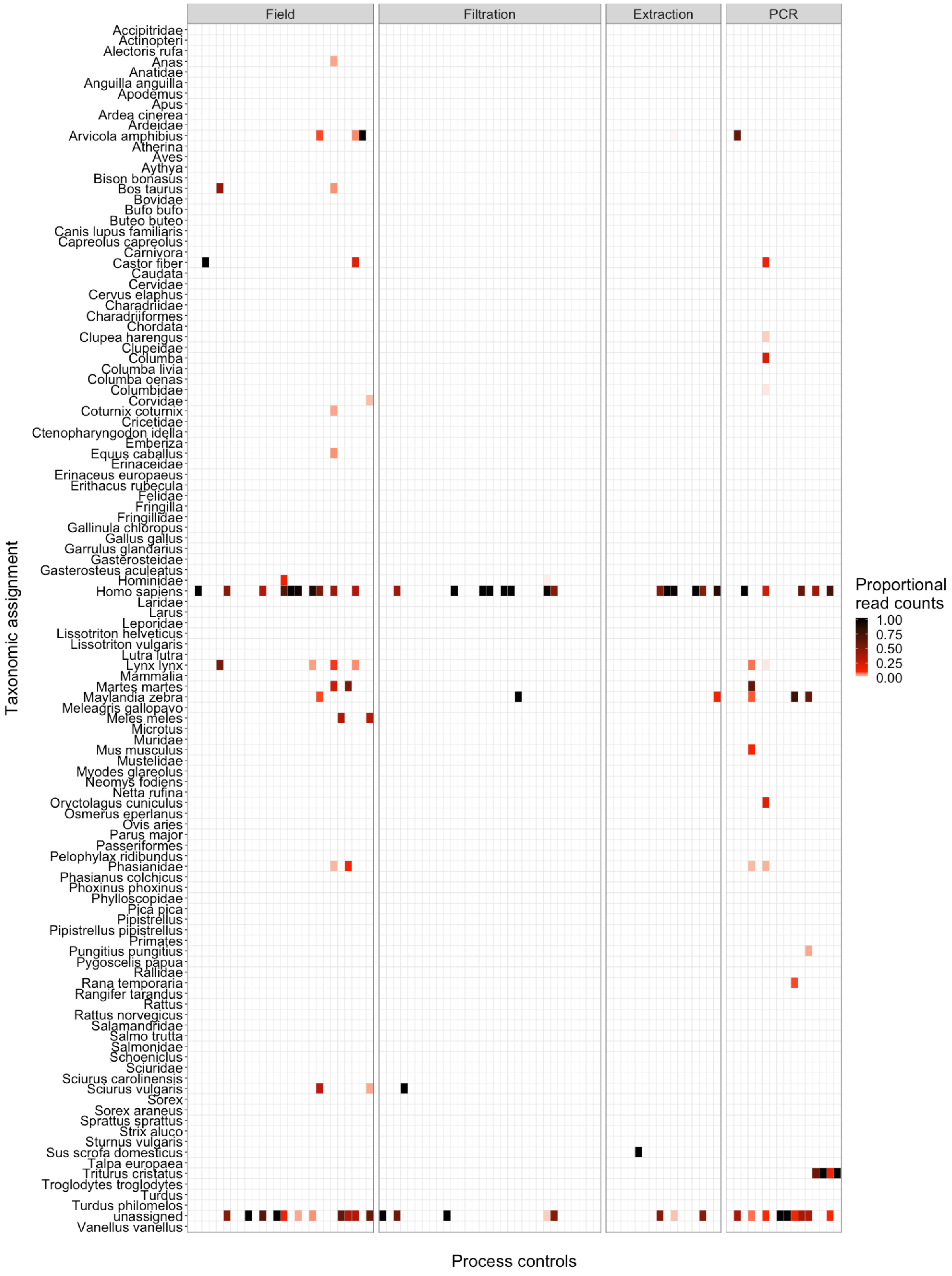


**Figure A4.** Heatmap showing the frequency of contamination in negative process controls (field blanks, filtration blanks, extraction blanks, and PCR negative controls). Assignments that were not detected in a given process control are coloured white.

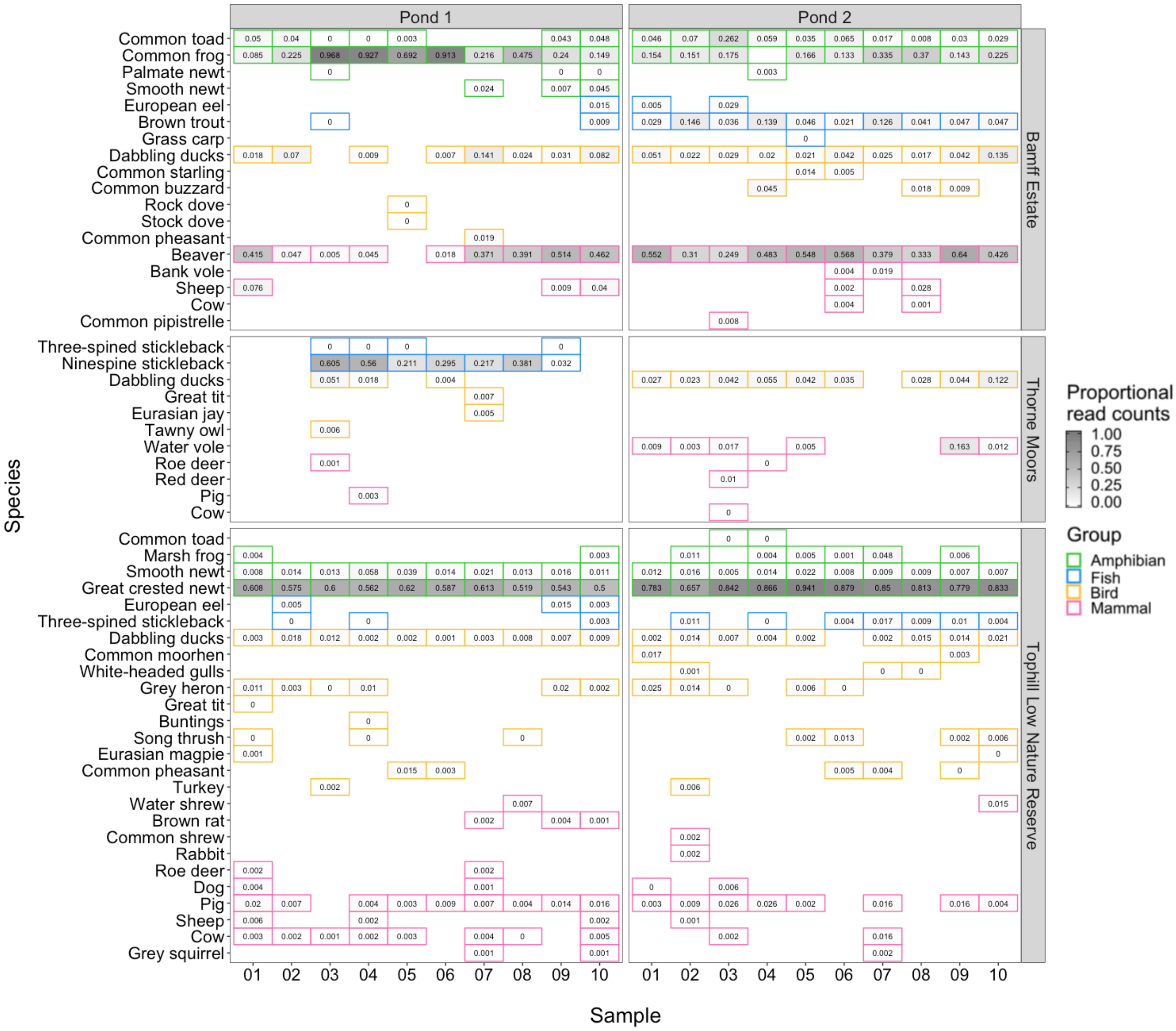


**Figure A5.** Heatmap showing proportional read counts for samples collected from natural ponds at sites where focal species were present. Each square represents a sample that had reads assigned to a vertebrate species. Species with low proportional read counts (i.e. more than 3 decimal places) are labeled 0.

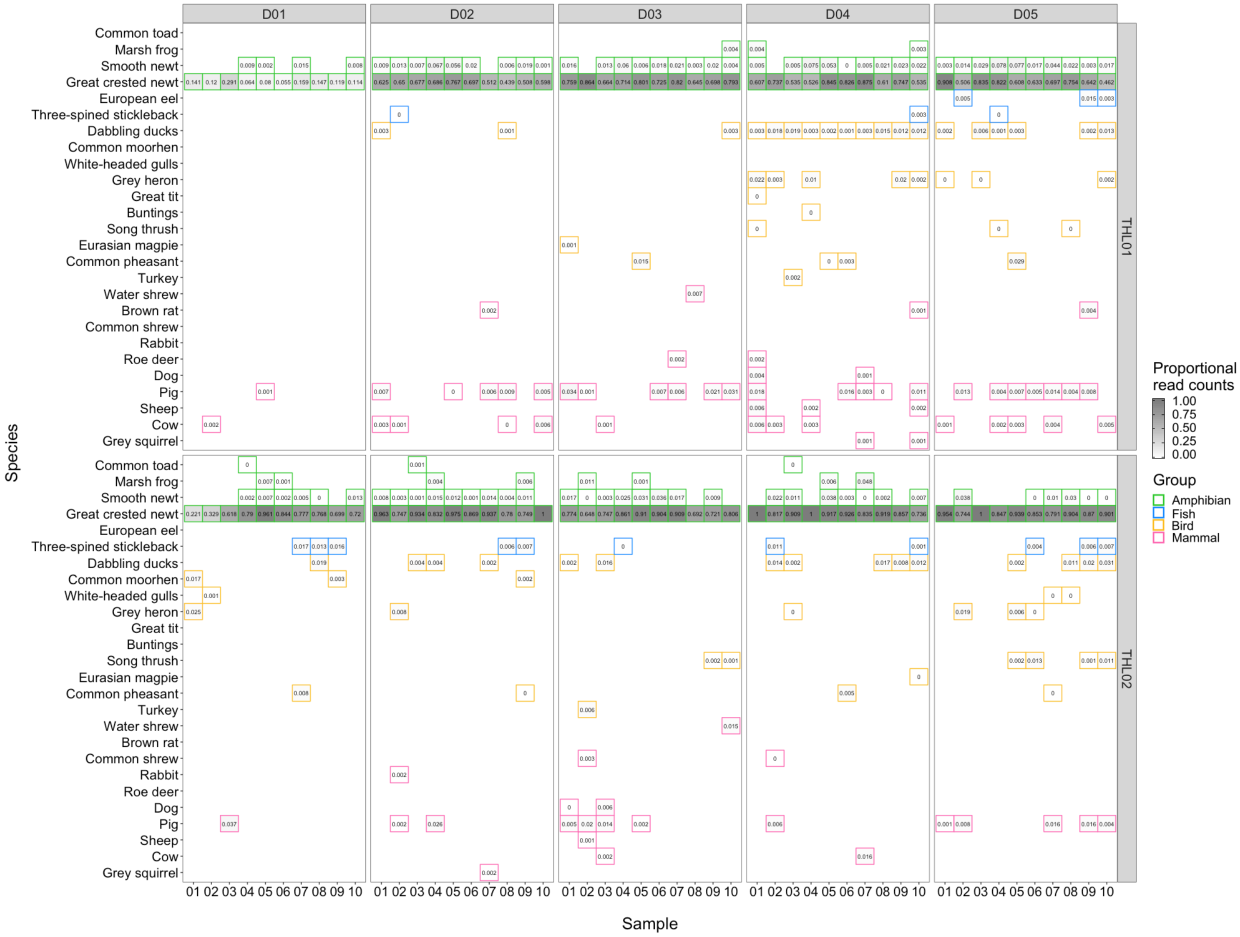


**Figure A6.** Heatmap showing species detected from samples collected at ponds (THL01 and THL02) within Tophill Low Nature Reserve every 24 hrs over a 5-day period (D01 - D05). Each square represents a sample that had reads assigned to a vertebrate species. Species with low proportional read counts (i.e. more than 3 decimal places) are labeled 0.
