## Appendix B for "Environmental DNA (eDNA) metabarcoding of pond water as a tool to survey conservation and management priority mammals"

**Wildwood Trust, Canterbury, Kent**

Otter (*Lutra lutra*)

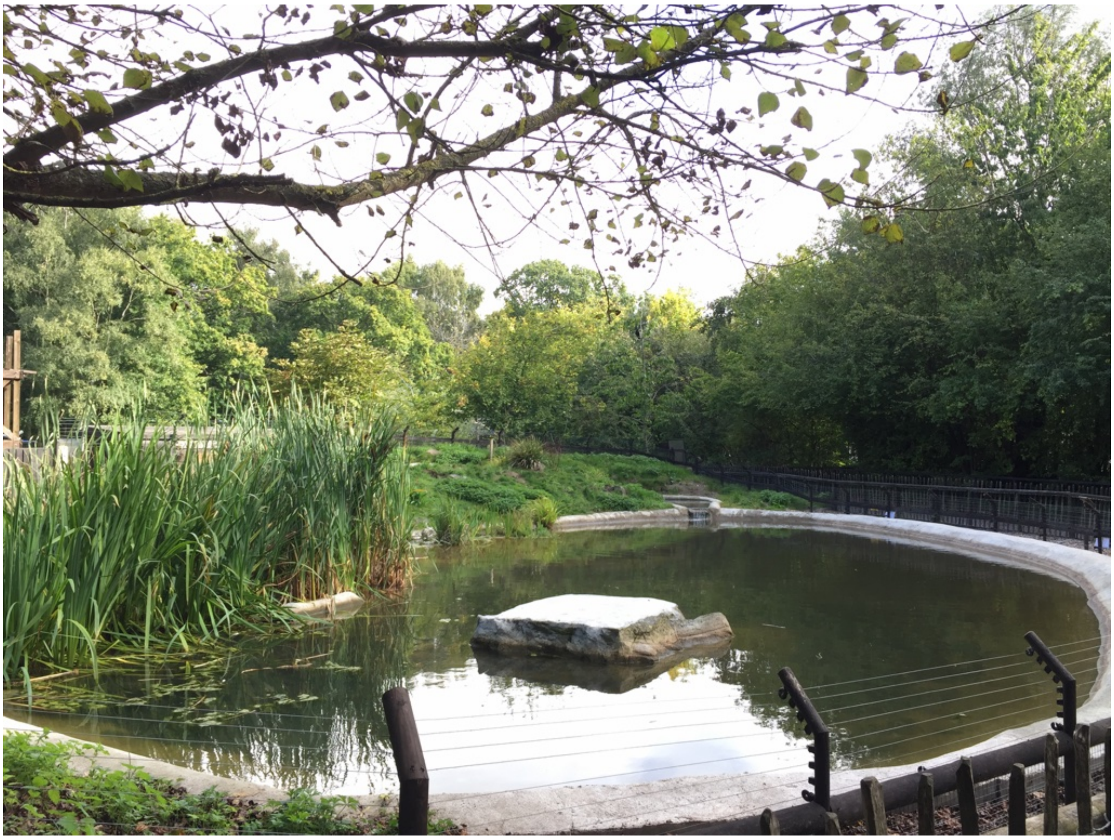

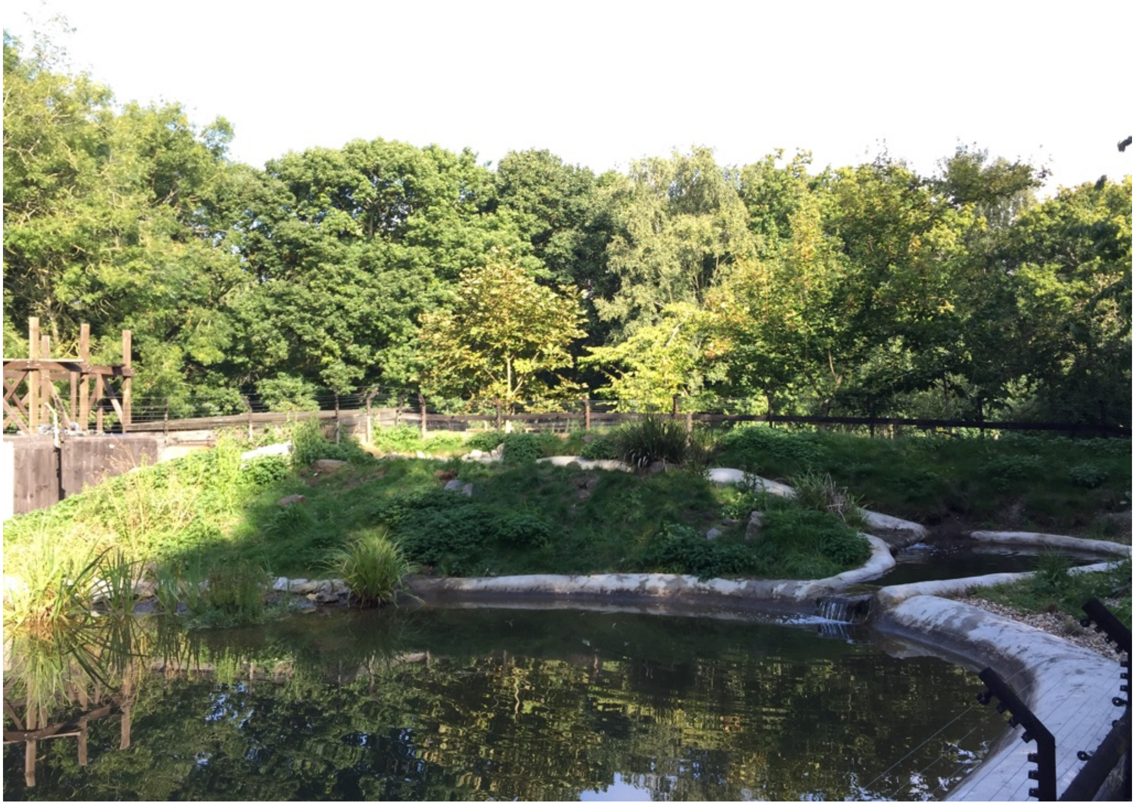

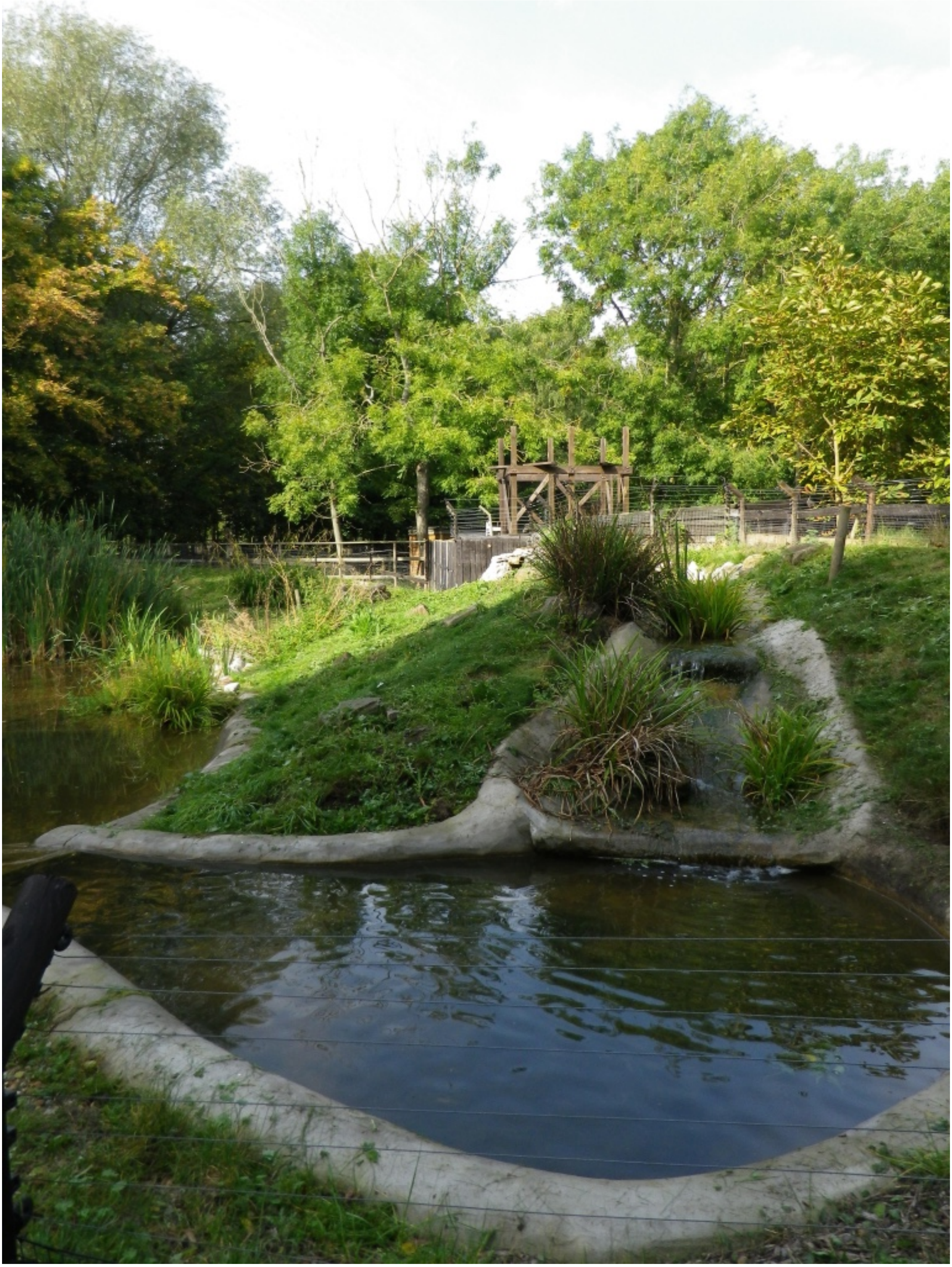

Beaver (*Castor fiber*)

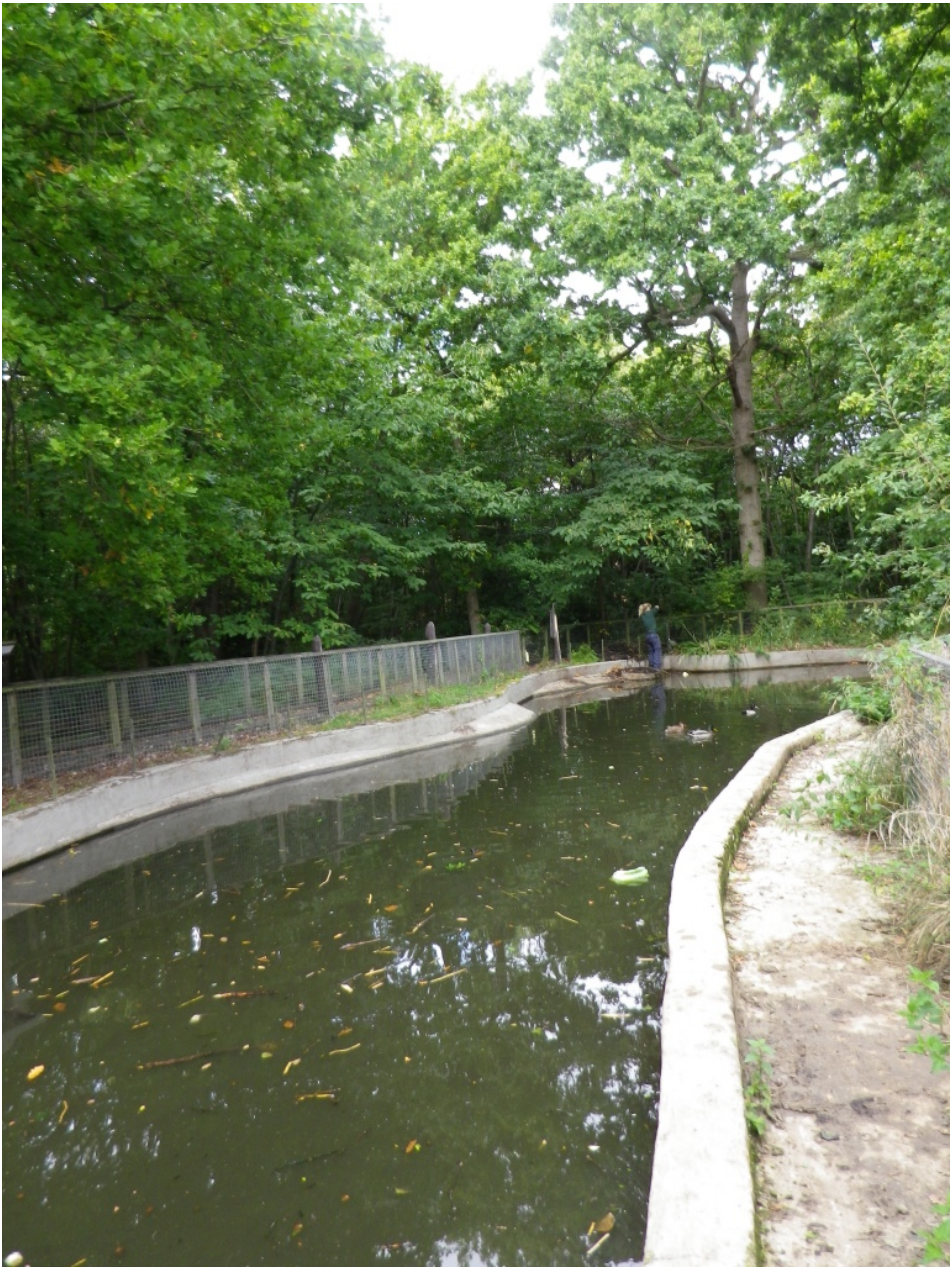

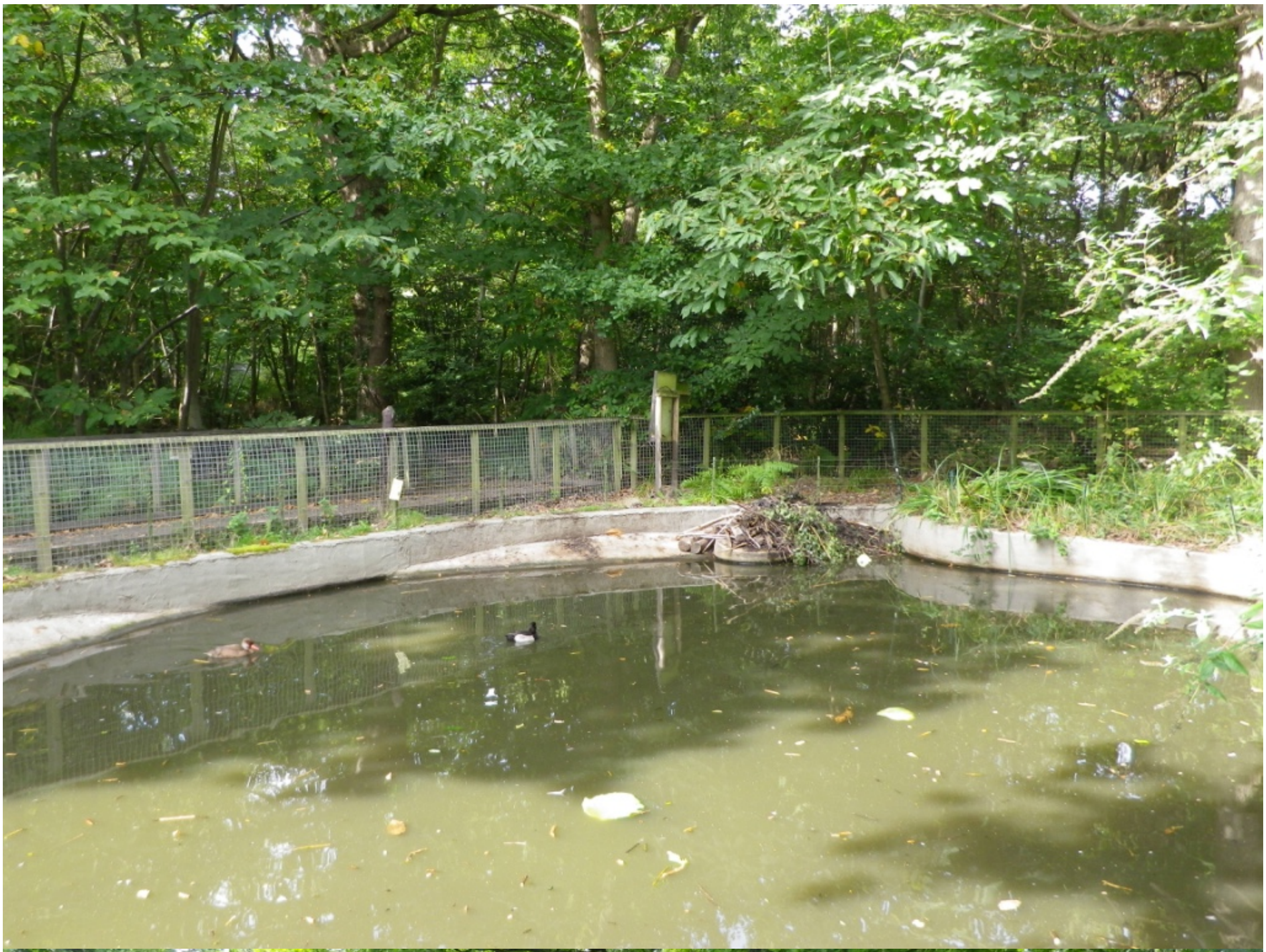

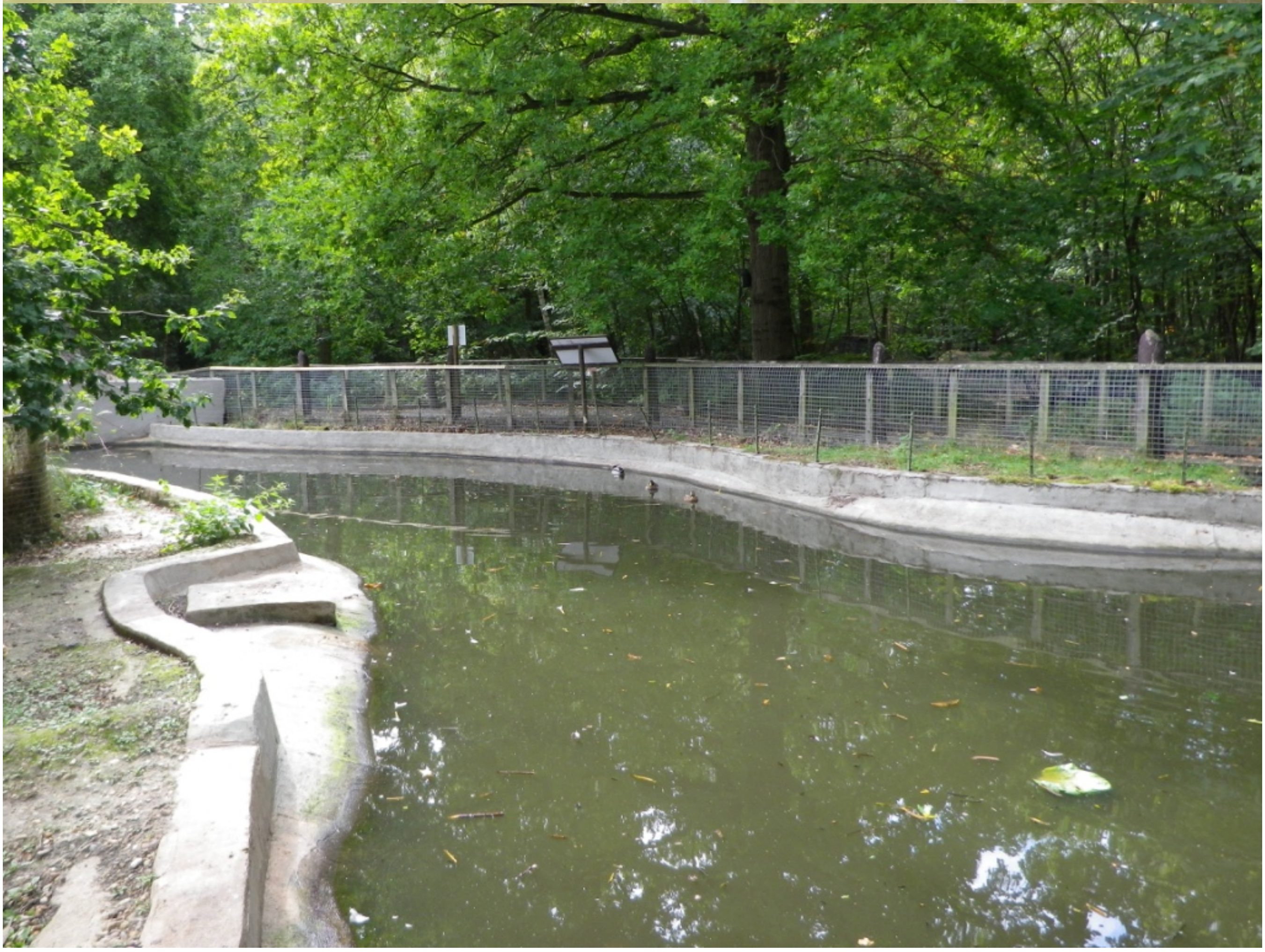

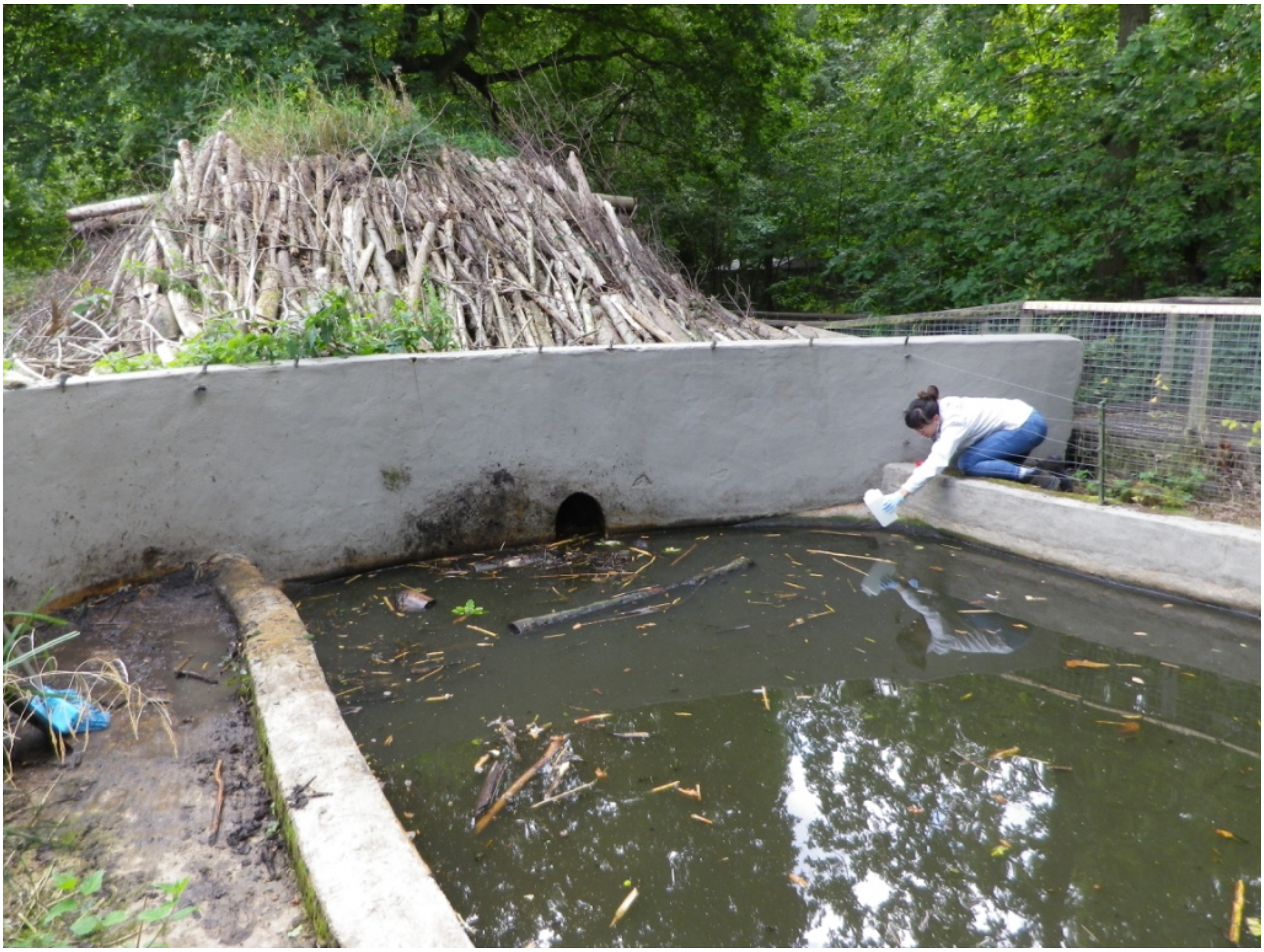

Water vole (*Arvicola amphibius*)

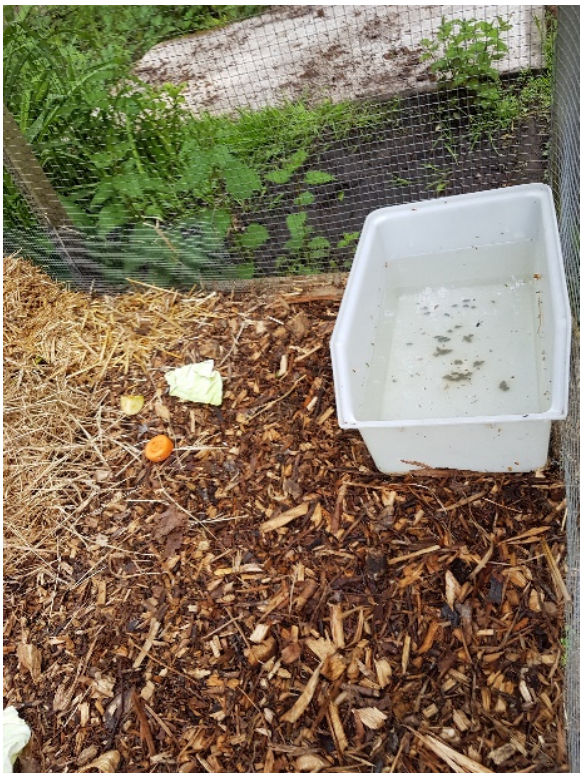

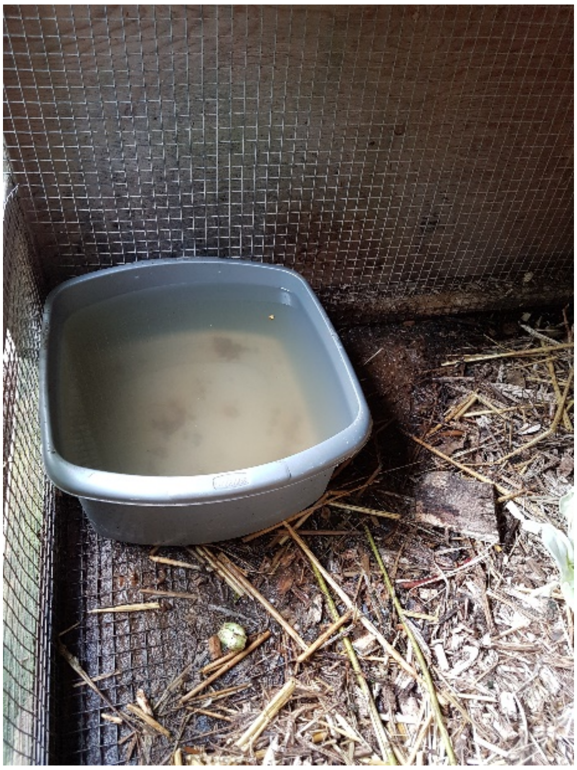

Hedgehog (*Erinaceus europaeus*)

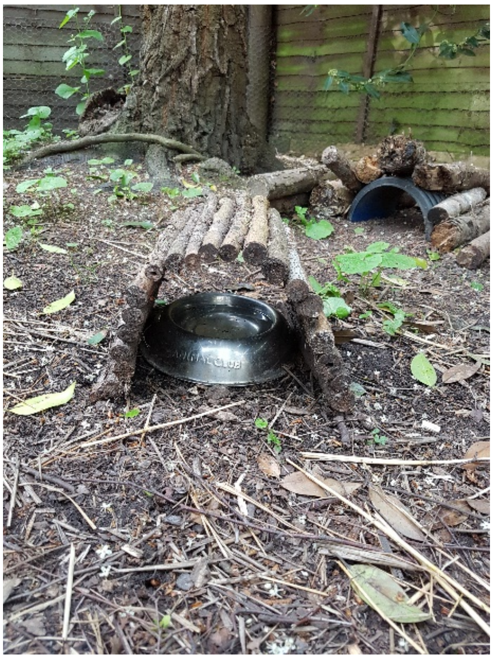

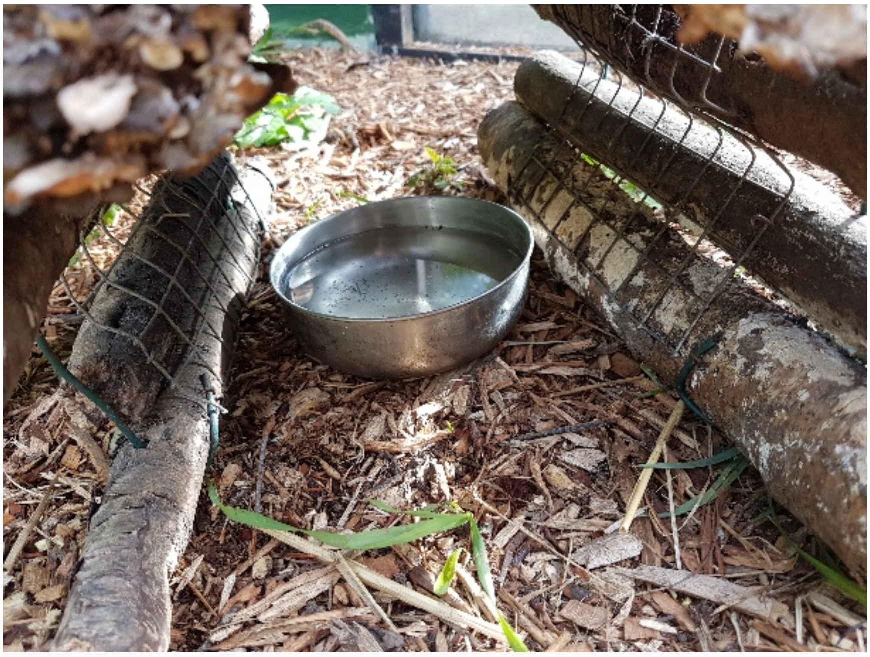

Badger (*Meles meles*)

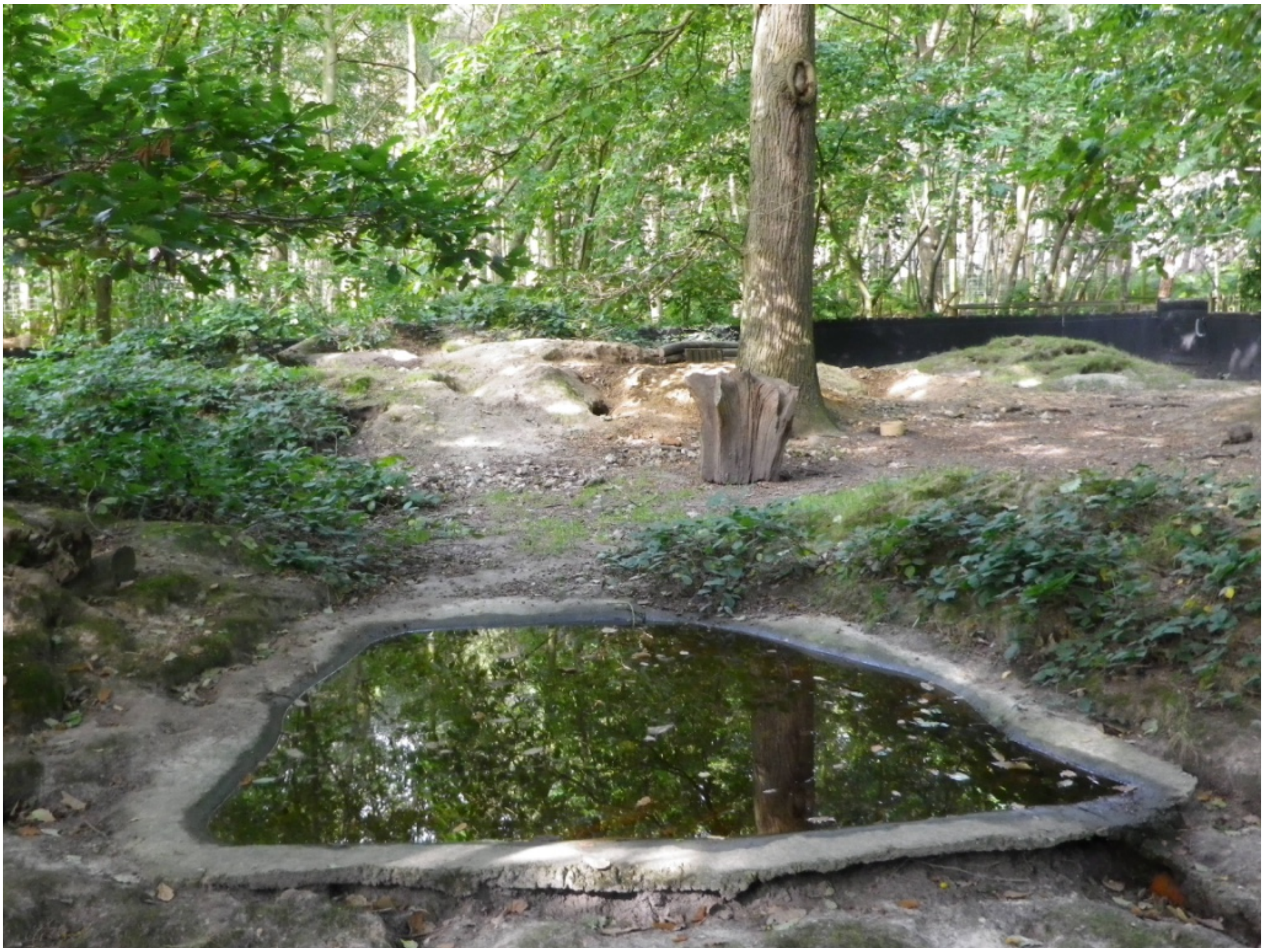

Lynx (*Lynx lynx*)

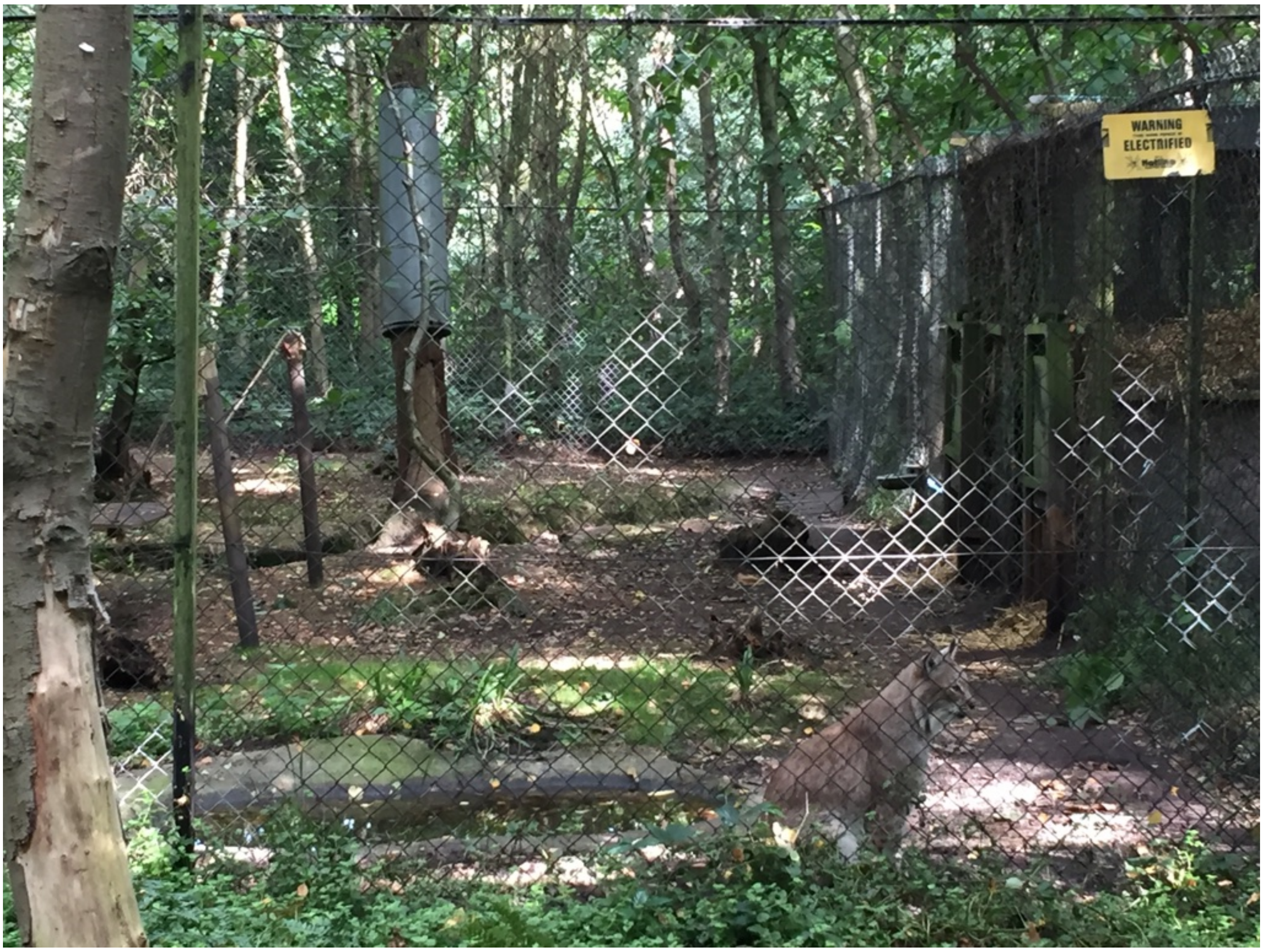


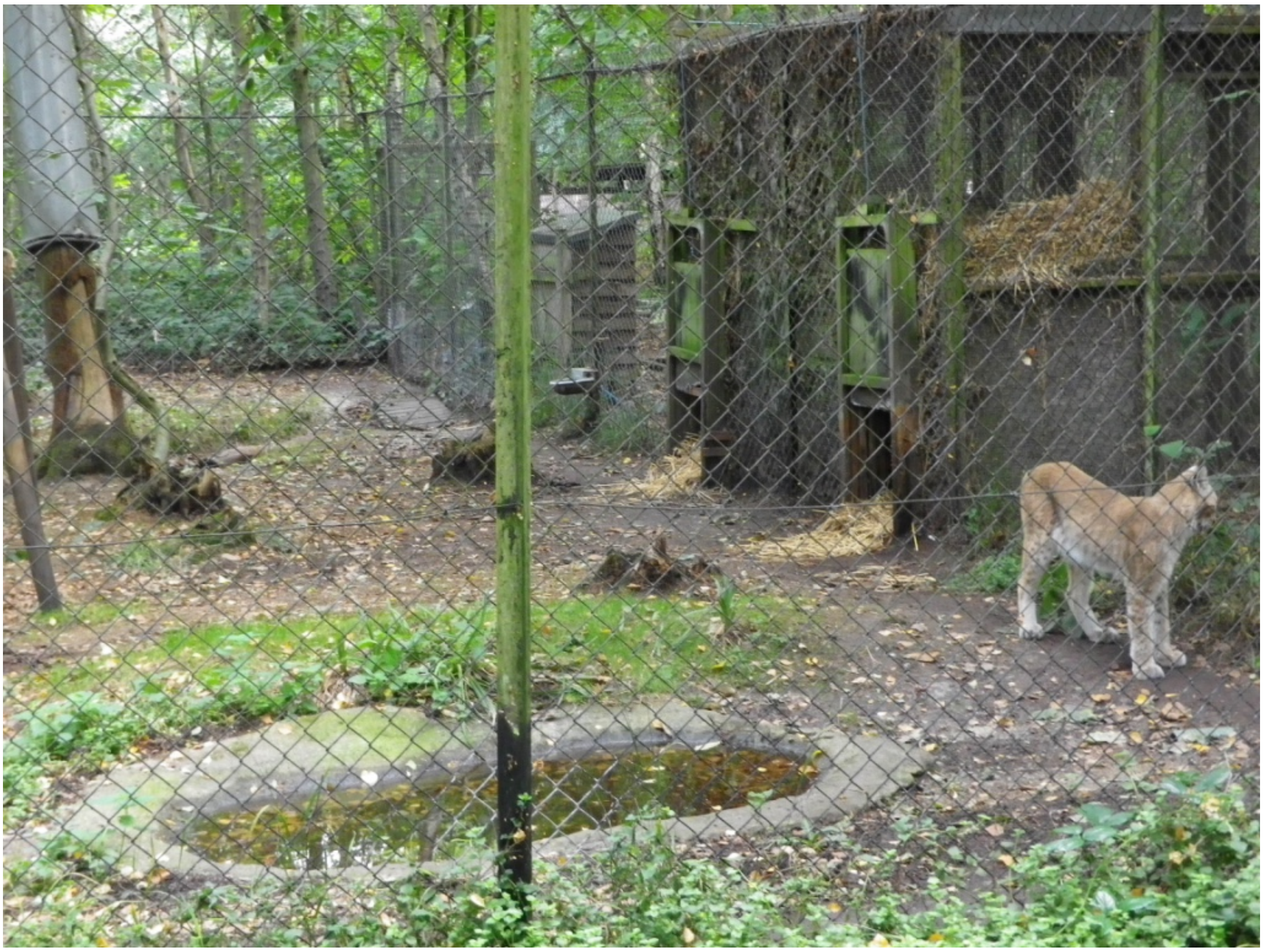

Red deer (*Cervus elaphus*)

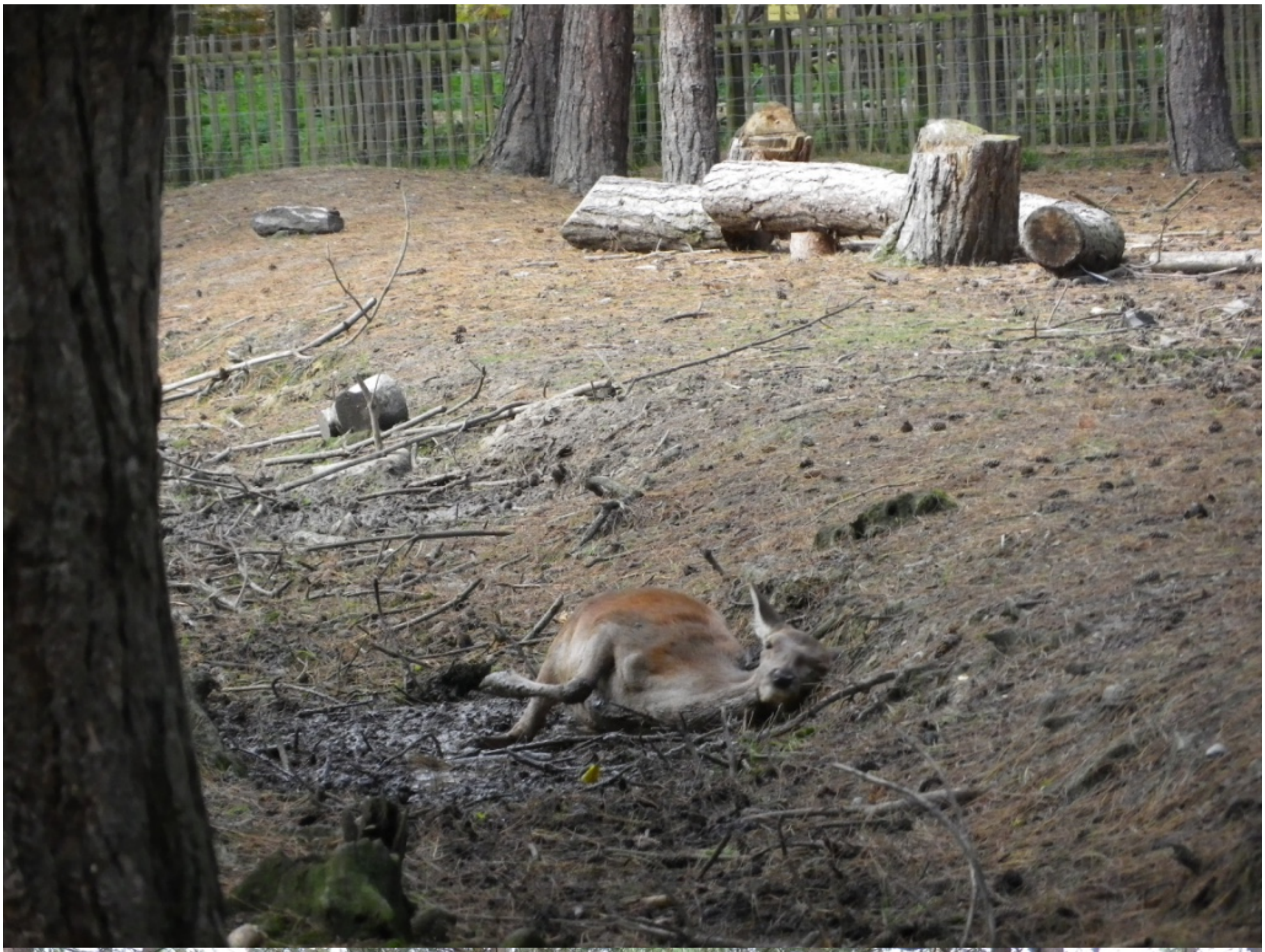

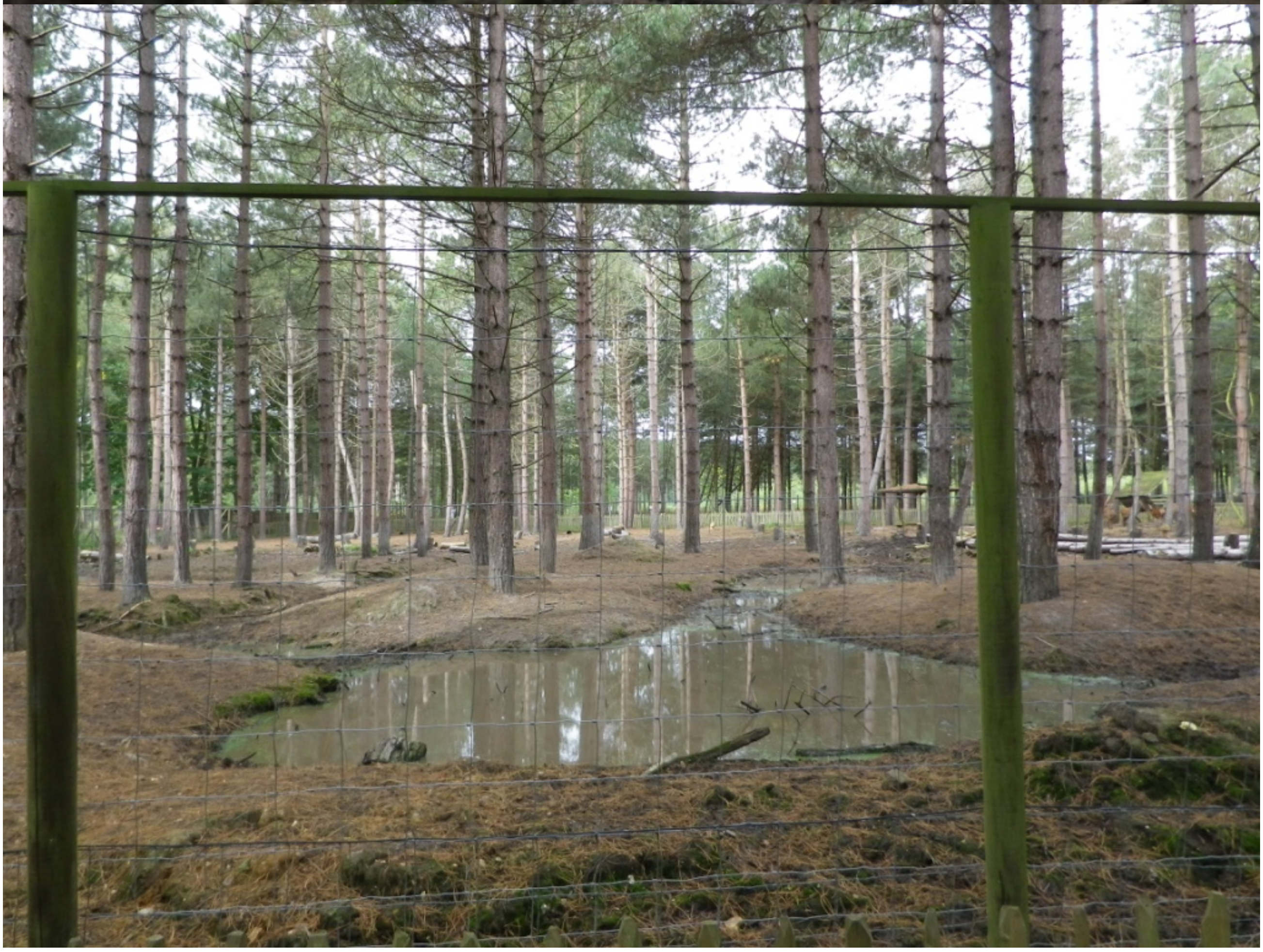

Red squirrel (*Sciurus vulgaris*)

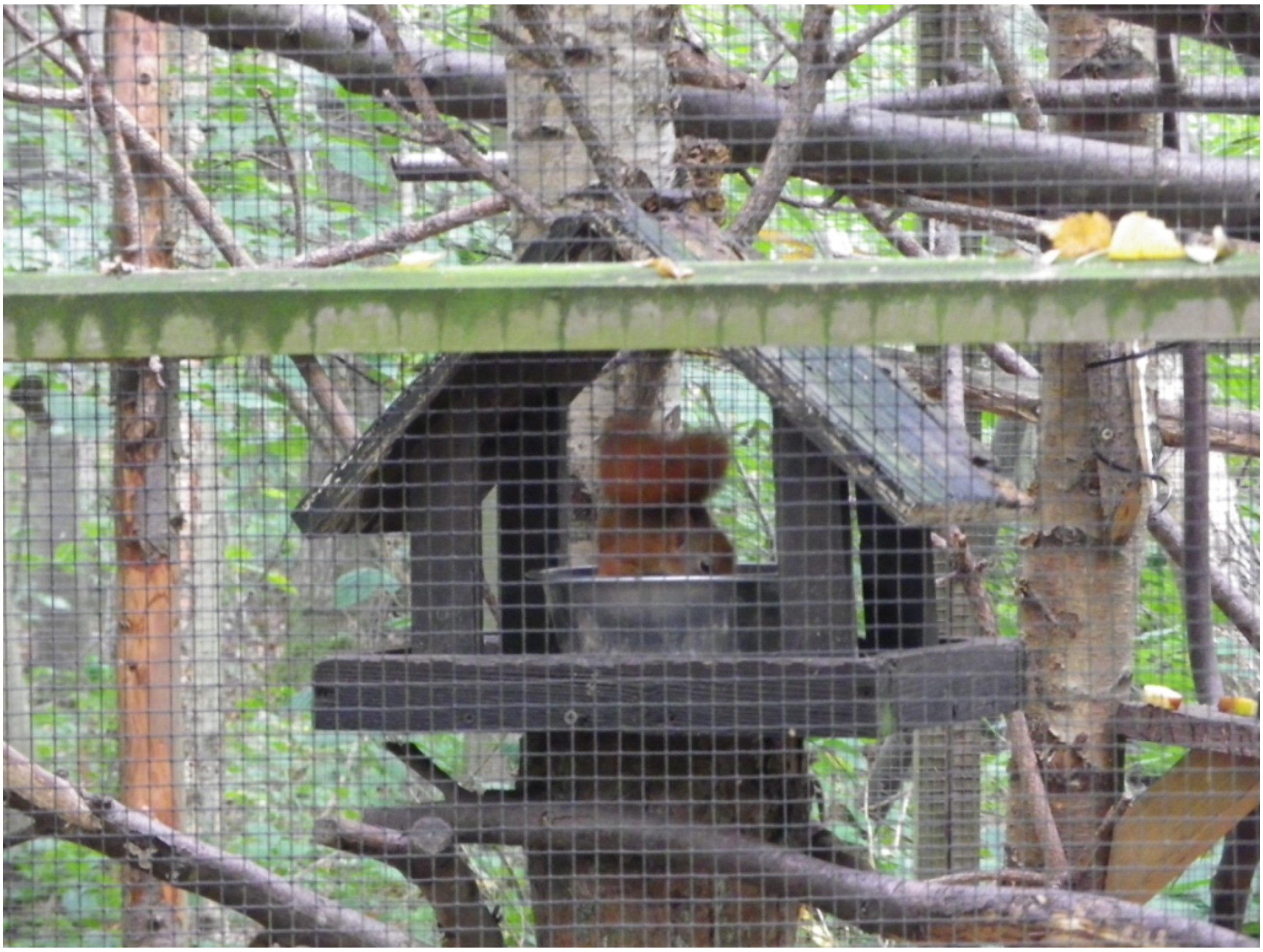

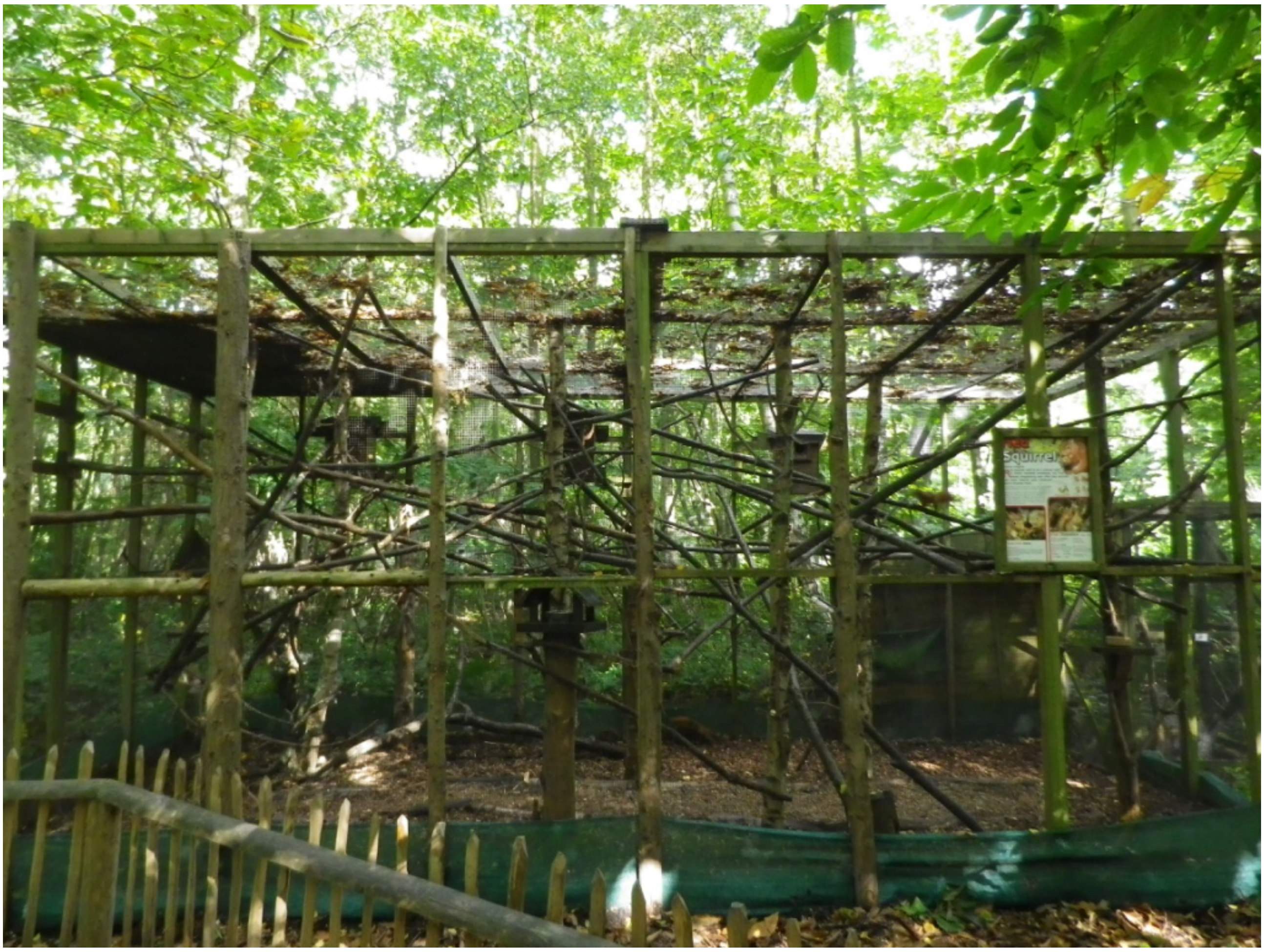

Pine marten (*Martes martes*)

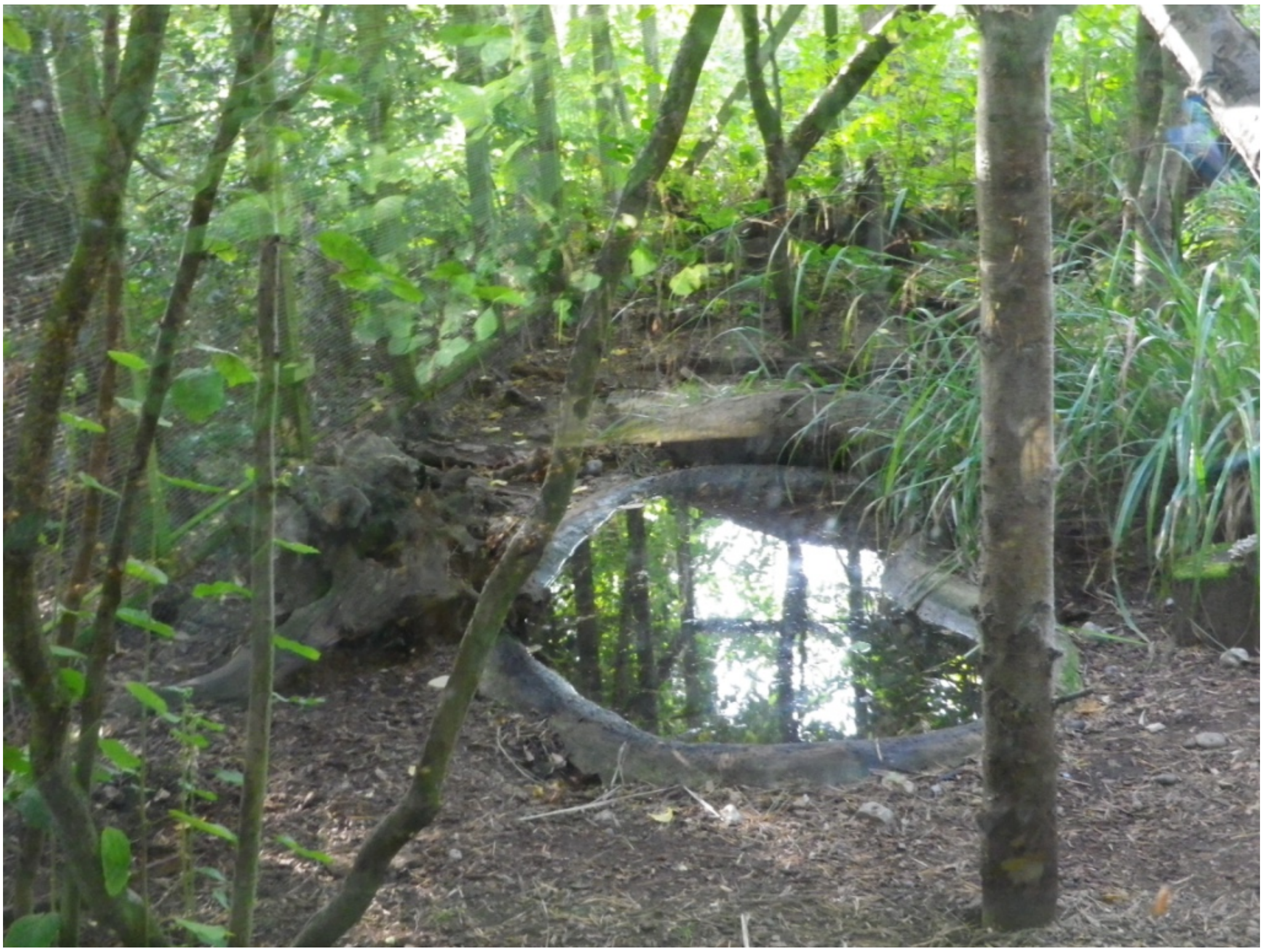

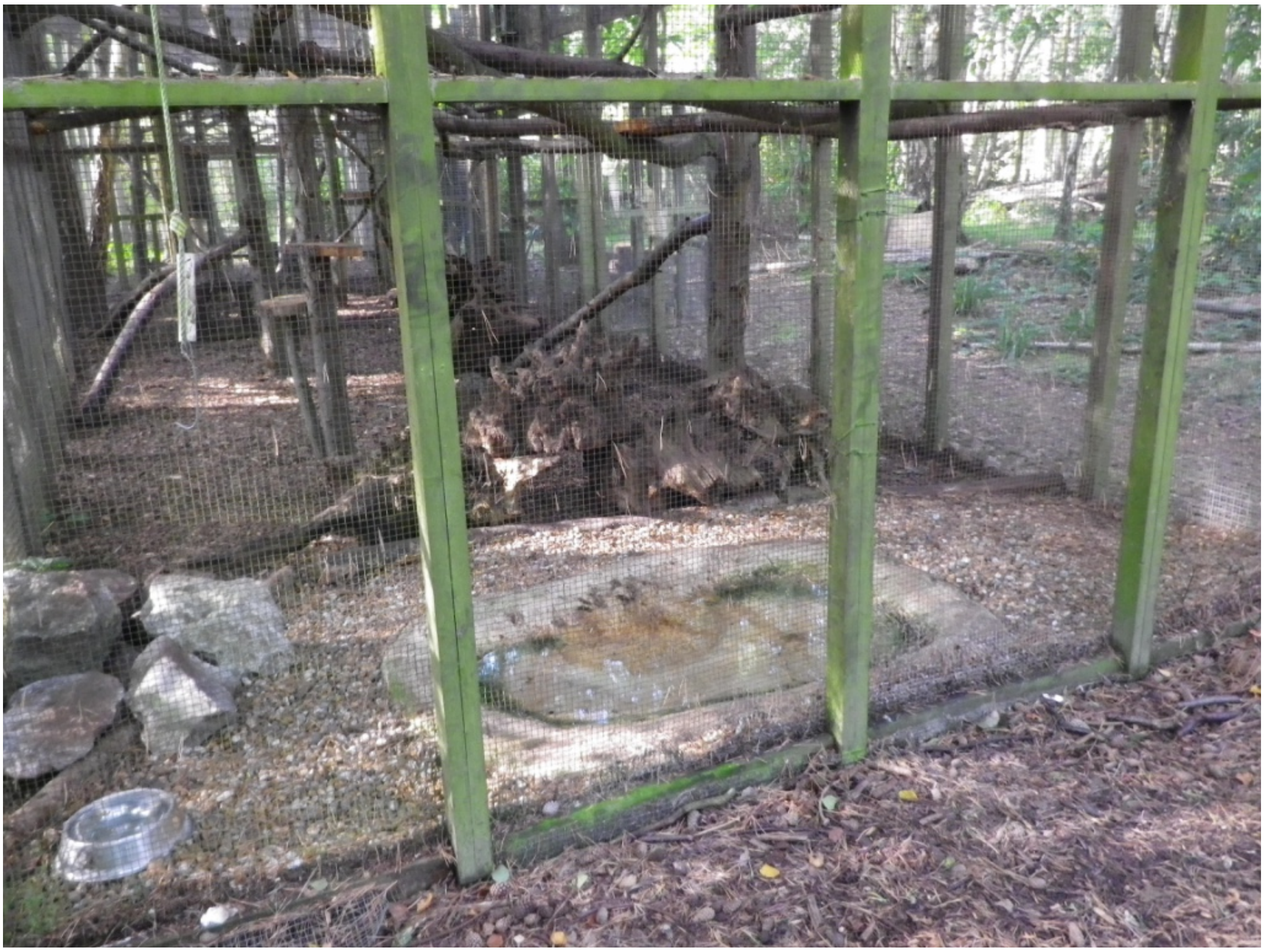

**Highland Wildlife Park, Kingussie, Scotland**

Water bath in woods

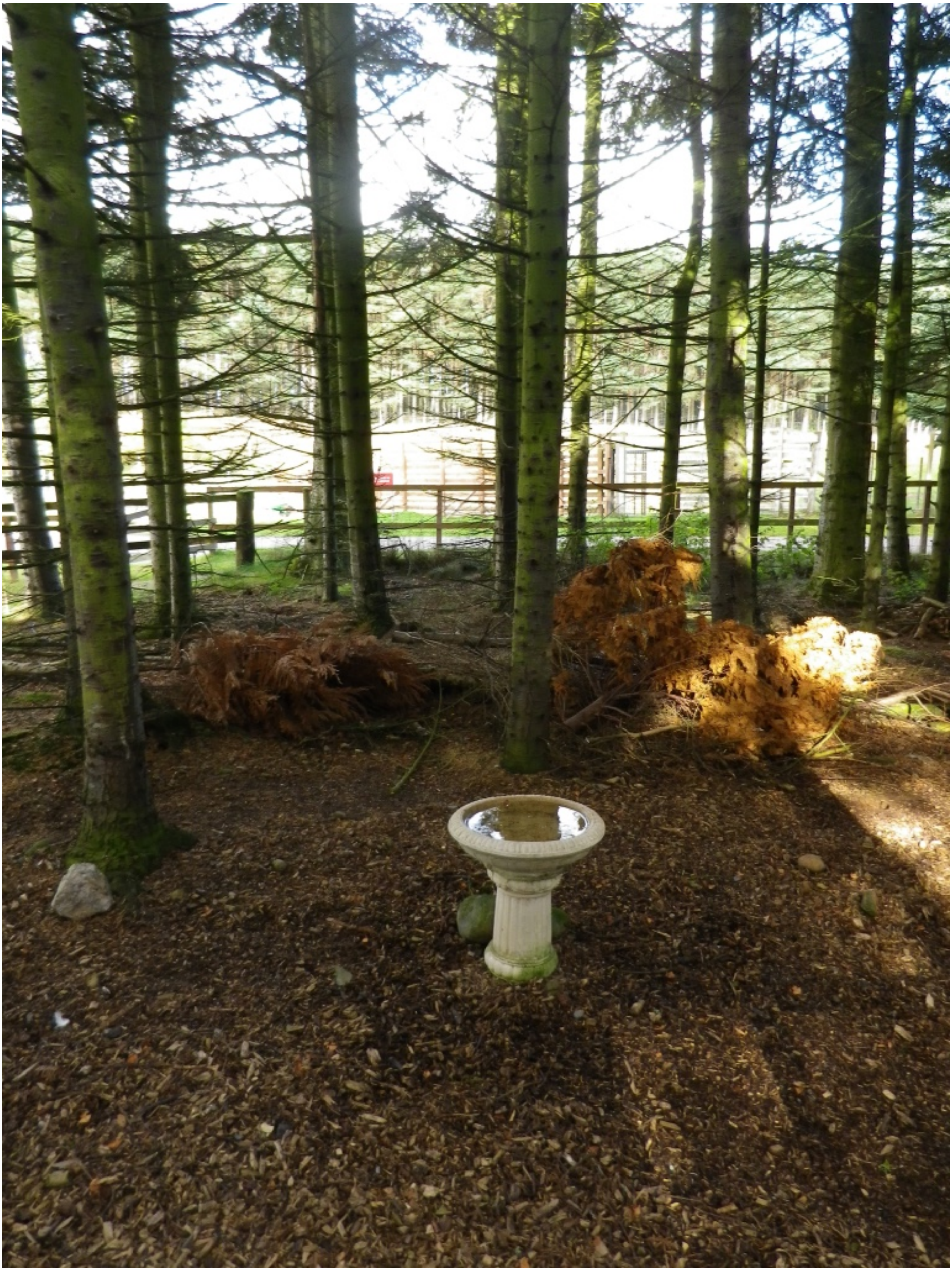

Lynx (*Lynx lynx*)

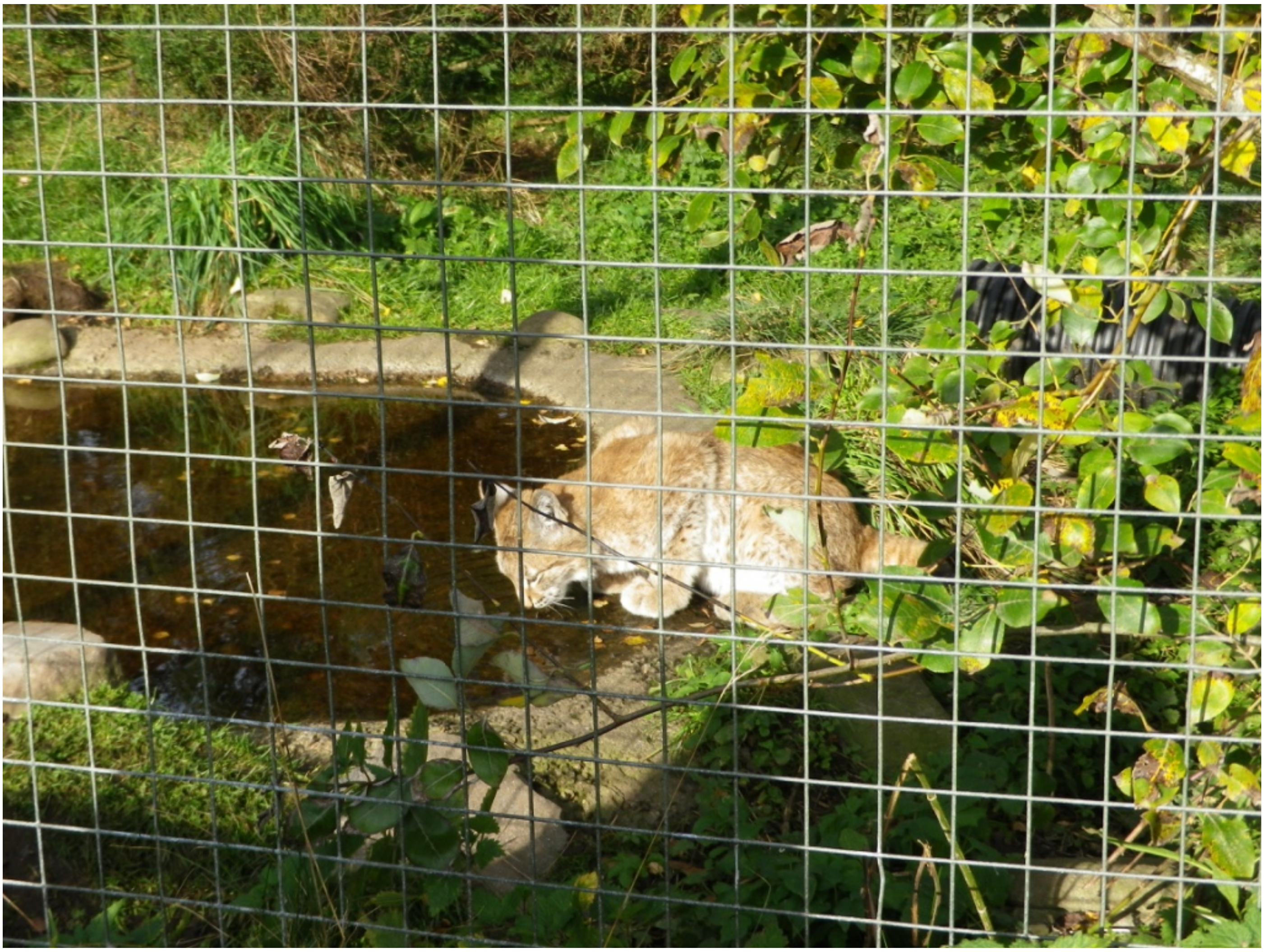

Red deer (*Cervus elaphus*)

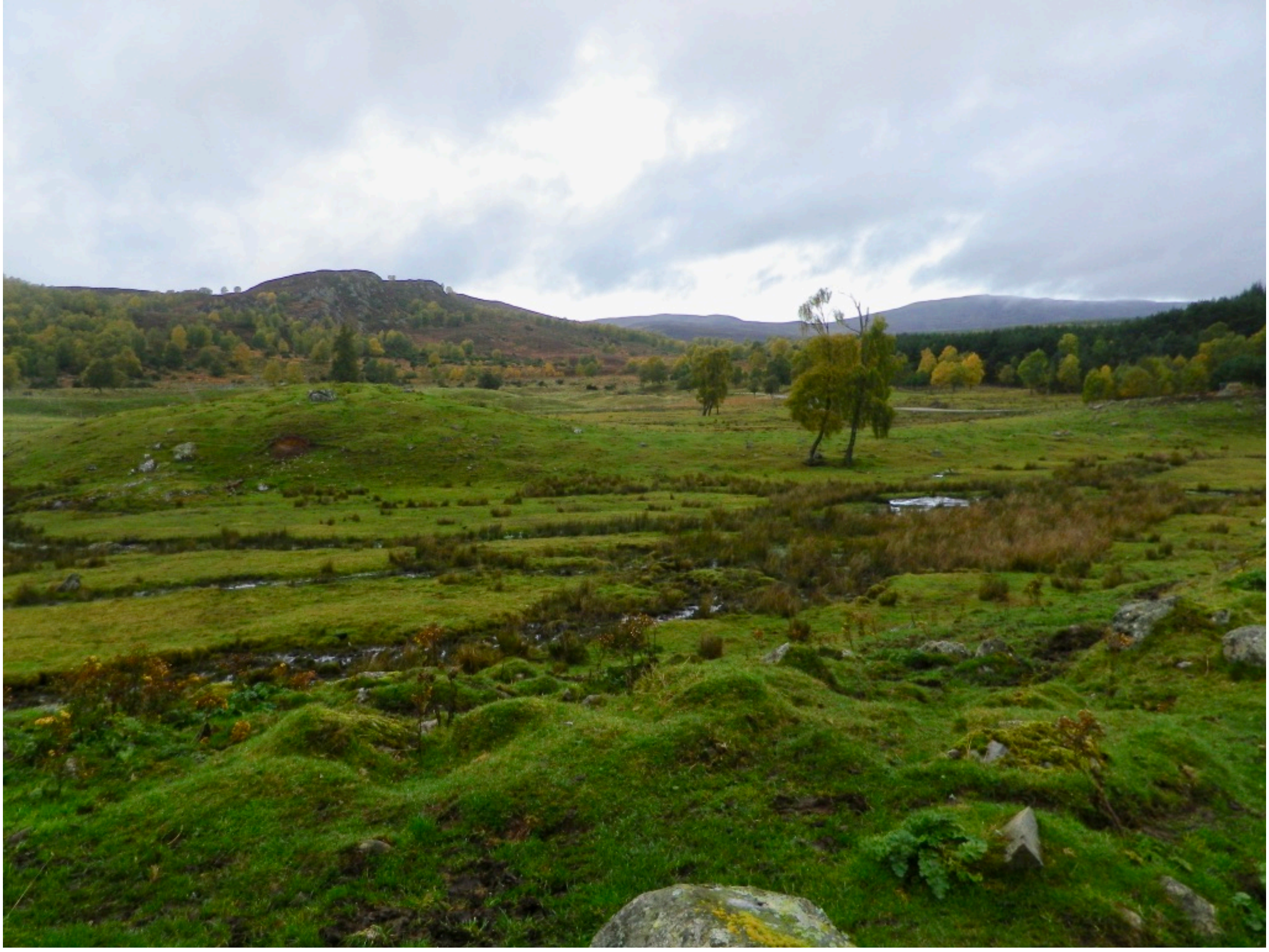

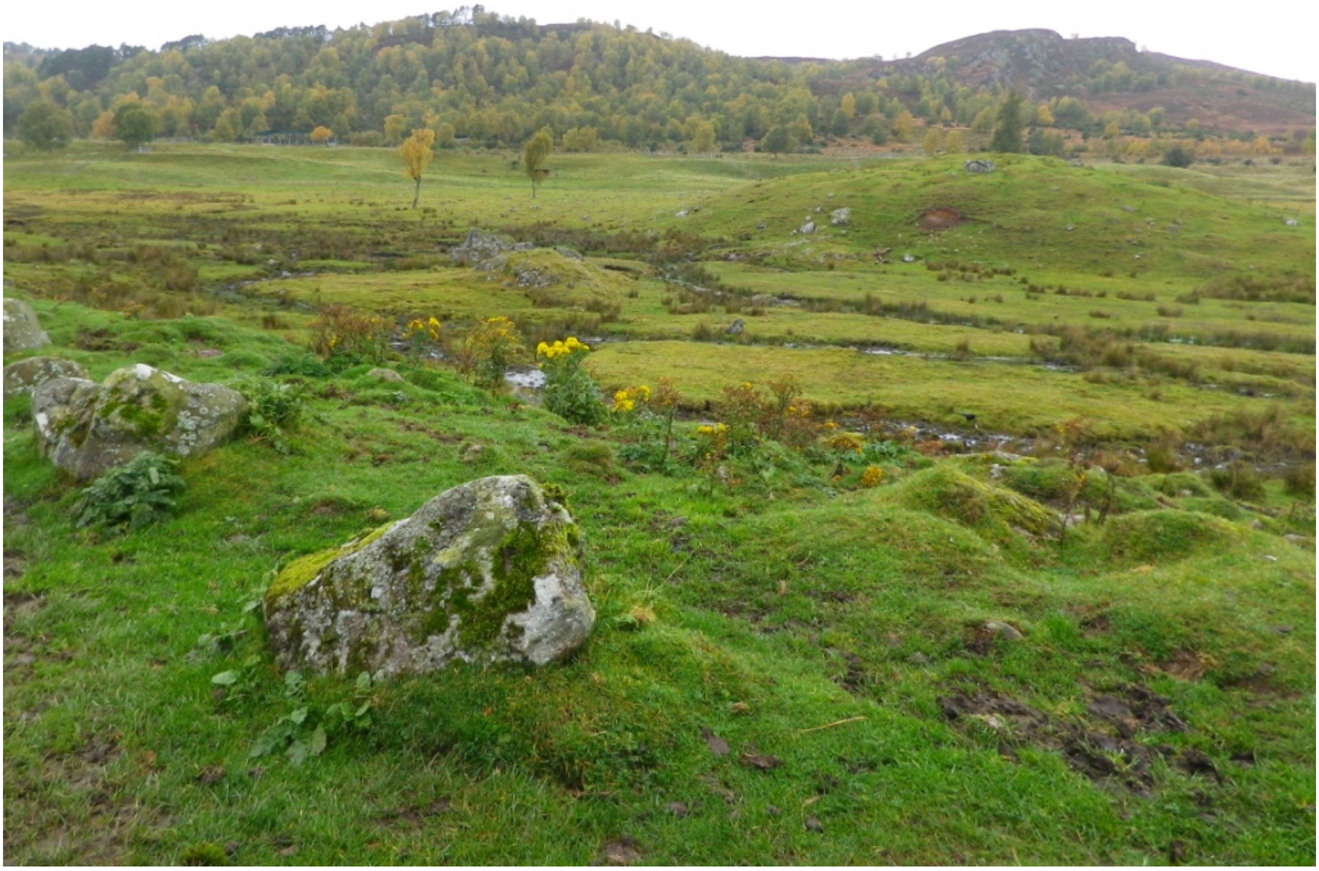

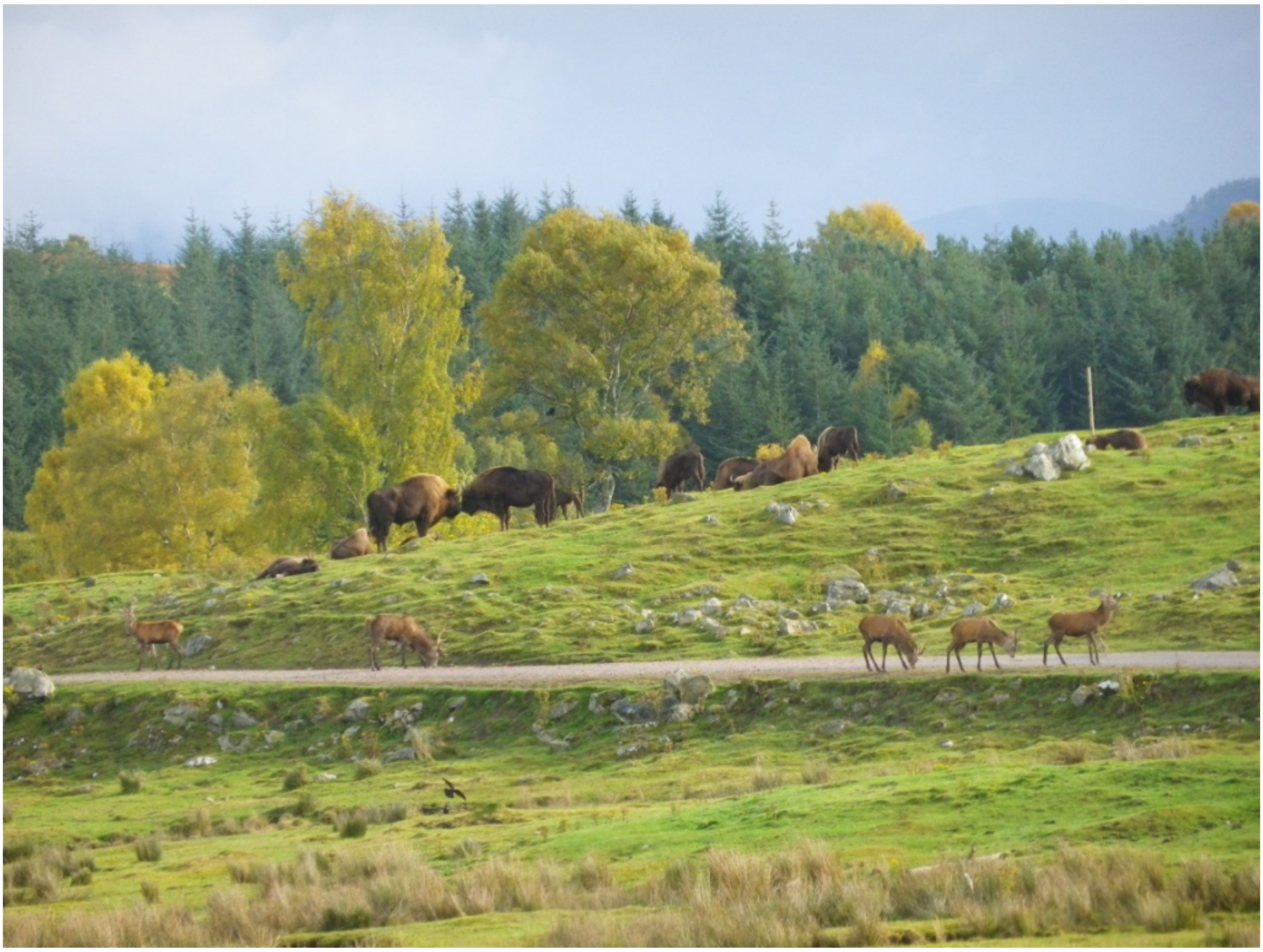

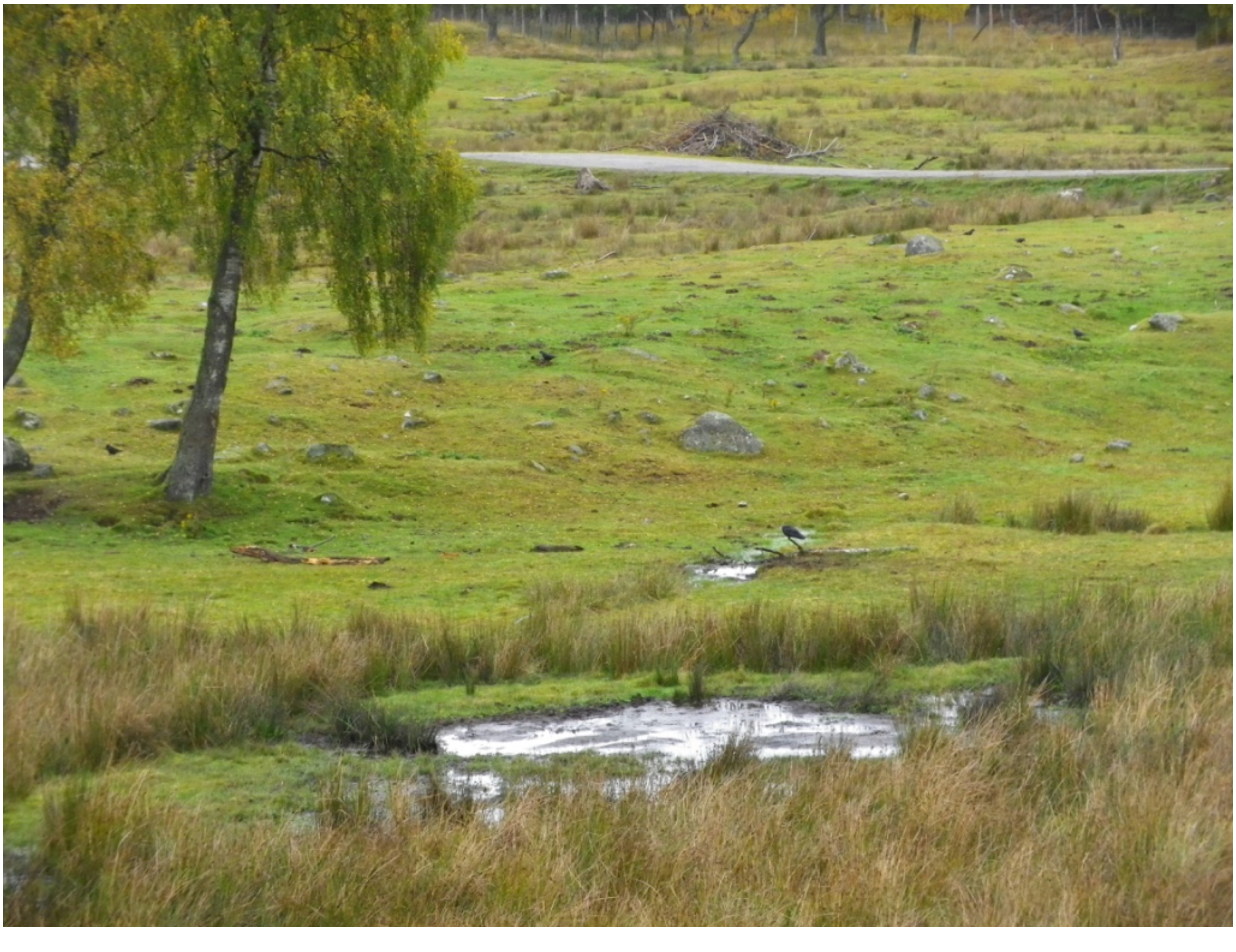

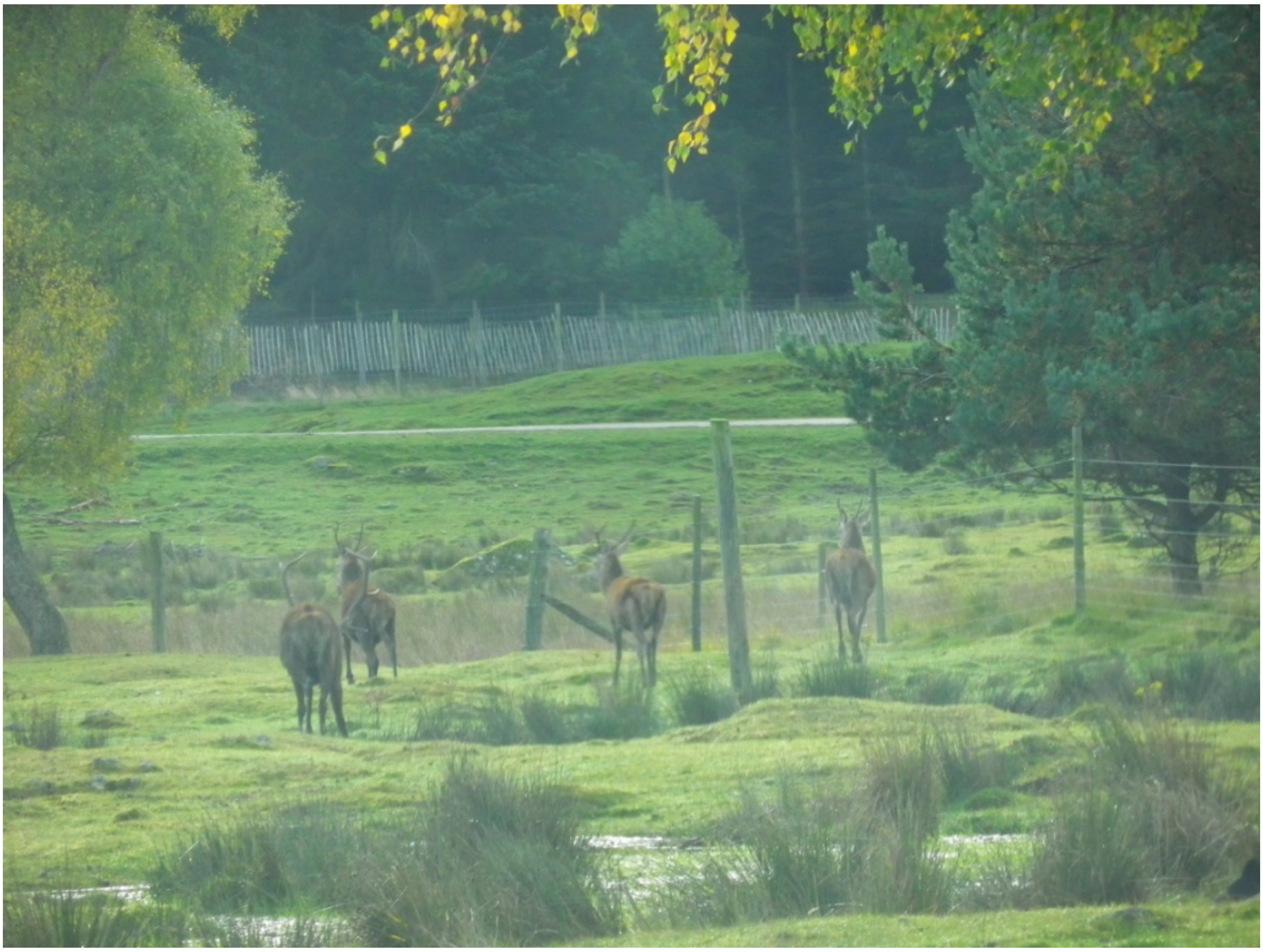
